# SeMOE allows for quantitative glycan perception and exhibits anti-cancer potentiality

**DOI:** 10.1101/2023.05.08.539922

**Authors:** Xiao Tian, Lingna Zheng, Changjiang Wang, Yida Han, Yujie Li, Tongxiao Cui, Jialin Liu, Chuanming Liu, Guogeng Jia, Lujie Yang, Chen Zeng, Lijun Ding, Chu Wang, Bo Cheng, Meng Wang, Ran Xie

## Abstract

Metabolic oligosaccharide engineering (MOE) is a classical chemical approach to perturb, profile and perceive glycans in physiological systems, but probes upon bioorthogonal reaction require accessibility and background signal readout makes it challenging to achieve absolute glycan quantification. Here we develop SeMOE, a selenosugar-based metabolic oligosaccharide engineering strategy that combines elemental analysis and MOE to enable the absolute quantification and mass spectrometric imaging of glycome in a concise procedure. We demonstrate that SeMOE probes allow for perception, absolute quantification and visualization of glycans in diverse biological contexts. We demonstrate that chemical reporters on conventional MOE can be integrated into a bifunctional SeMOE probe to provide multimodality signal readouts. We further show the anti-cancer potentiality of SeMOE probes. SeMOE thus provides a convenient and simplified method to “see more” of the glyco-world.

---

Metabolic oligosaccharide engineering (MOE) is the most appealing chemical glycobiological approach to probe, perturb and perceive glycans^1, 2^. Cells or organisms are treated with new-to-nature carbohydrate precursors/analogs that are modified with an organic chemical reporter (*e.g.,* azide, alkyne). Such compounds are converted into glycoenzyme substrates within the cell, and metabolically incorporated into newly-synthesized glycoconjugates. Subsequent ligation on the reporter groups via the Nobel Prize-winning bioorthogonal reaction enables the dynamic and versatile glycan visualization, as well as glycoconjugate isolation and -omic analysis (Fig. 1a)^3, 4^. So far, derivatizations on *N*-acetylmannosamine (ManNAc), sialic acid (Sia), *N*-acetylgalactosamine (GalNAc), *N*-acetylglucosamine (GlcNAc), and fucose (Fuc), with or without acetylation on hydroxyl groups for better plasma membrane diffusion, has been realized as the MOE probes^5^. Although intricate protocols have been developed for MOE, stoichiometric attachment of fluorophores or tags to glycans is inevitably hampered by their accessibility, as well as the background signal readout, making it challenging to achieve glycan quantification by MOE. Endeavors have been made by altering readout modalities via click reaction^6, 7^, yet a simple and direct glycan quantification methodology is still lacking.

**Fig. 1.**
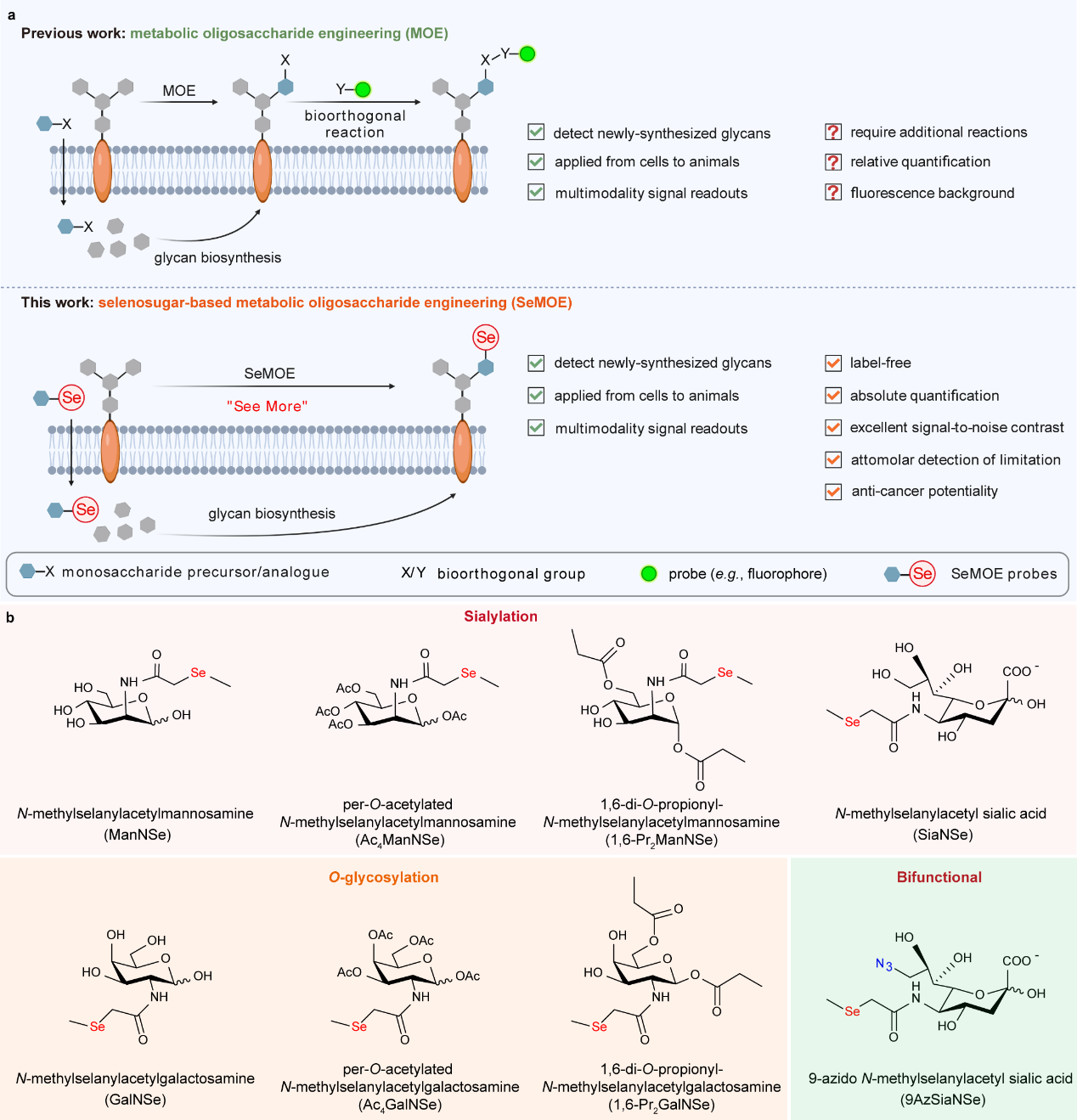
| Schematic of SeMOE methodology. **a,** Comparison of SeMOE and classical MOE. Instead of using monosaccharide reporter bearing bioorthogonal functionalities (e.g., azide) in standard MOE procedure, selenium-containing probes were metabolically incorporated for label-free glycan detection based on elemental analysis. **b,** Selenosugars used in this study.

To expand the applicability of MOE, herein we report a selenosugar-based metabolic oligosaccharide engineering strategy (SeMOE) that combines elemental analysis and MOE to enable the absolute quantification and mass spectrometric imaging of glycome in a concise procedure (Fig. 1a). We demonstrate that the label-free SeMOE probes allow for perception, absolute quantification and visualization of glycans in diverse biological contexts. In addition, chemical reporters on conventional MOE can be integrated into a bifunctional SeMOE probe to provide multimodality signal readouts. Finally, the anti-cancer potentiality of SeMOE probes is also demonstrated.

## Development of SeMOE probes

Selenium is an essential trace nutrient for cellular functions in the physiological environment, mostly in the form of unusual amino acids selenocysteine and selenomethionine, to form selenoproteins^8^. Selenium can be quantitatively analyzed and imaged by elemental analysis methods, such as inductively coupled plasma mass spectrometry (ICP-MS), a powerful multi-element trace analysis and isotope ratio measurement at the trace and ultra-trace level in life sciences^9^. We chose to introduce the selenium element in the form of *N*-methylselanylacetyl side chain for sialic acid, mannosamine and galactosamine, because previous literatures has solidified that the glycan biosynthetic machinery can tolerate the subtle chemical modification on these analogs, and effectively install them into cellular sialoglycans and *O*-glycans in the same way as its natural counterpart, respectively^10^. We therefore synthesized a total of 8 selenium-containing metabolic monosaccharide precursors/analogs, which we termed “selenosugars” or “SeMOE probes” thereafter (Fig. 1b, **Supplementary Note 1**). Of note, per-*O*-acetylated, partially *O*-propionylated, as well as unprotected versions of selenosugars were all used in this work to ensure successful metabolic incorporation via SeMOE. Once incorporated, a selenium spike should be observed upon minimal-to-none background elemental signal, without the need for secondary labeling in classical MOE, with guaranteed signal-to-noise contrast (Fig. 1a). And the unique property of selenium, such as redox response, can be exploited instead to interrogate glycan functions in their native surroundings.

## SeMOE enables label-free glycan detection and absolute quantification

We started systematically evaluating SeMOE strategy feasibility. A previous work reported that 1β-methylseleno-*N*-acetylgalactosamine is the key metabolite for selenium detoxication, implying that selenosugar is non-toxic^11, 12^. As expected, CCK-8 assays on the synthesized selenosugars exhibited cytotoxicities comparable with native monosaccharide precursor or azido sialic acid at the range of 0-5 mM in various mammalian cell lines (**Supplementary Fig. 1**). To verify the incorporation of selenosugars, HeLa cells were incubated with selenosugars varying from 50 μM to 2 mM, lyzed for protein extraction, and analyzed using ICP-MS. We observed concentration- and time-dependent increase in the total Se level with minimal background noise, as normalized by the ^78^Se standard curve (Fig. 2a, Extended Data Fig. 1, **Supplementary Fig. 2**). Se readouts displayed selenosugar type-correlated difference, with a Se signal: Ac_4_ManNSe>1,6-Pr_2_ManNSe>SiaNSe>9AzSiaNSe>ManNSe, consistent with the order for classical MOE incorporation efficacy (*i.e,* azido expression measured via bioorthogonal reaction)^13^, thus implying selenosugars remain intact under protein glycosylation processes. Impressively, the label-free SeMOE provided nanomolar detection sensitivity. The titration experiment on HeLa cells with 1,6-Pr_2_ManNSe showed a signal-to-noise ratio at 6.2 at a concentration as low as 1 nM, up to 5 orders of magnitude better compared with the corresponding fluorescent detection for the azido counterpart (1,6-Pr_2_ManNAz) (Fig. 2b, **Supplementary Fig. 3**).

**Fig. 2.**
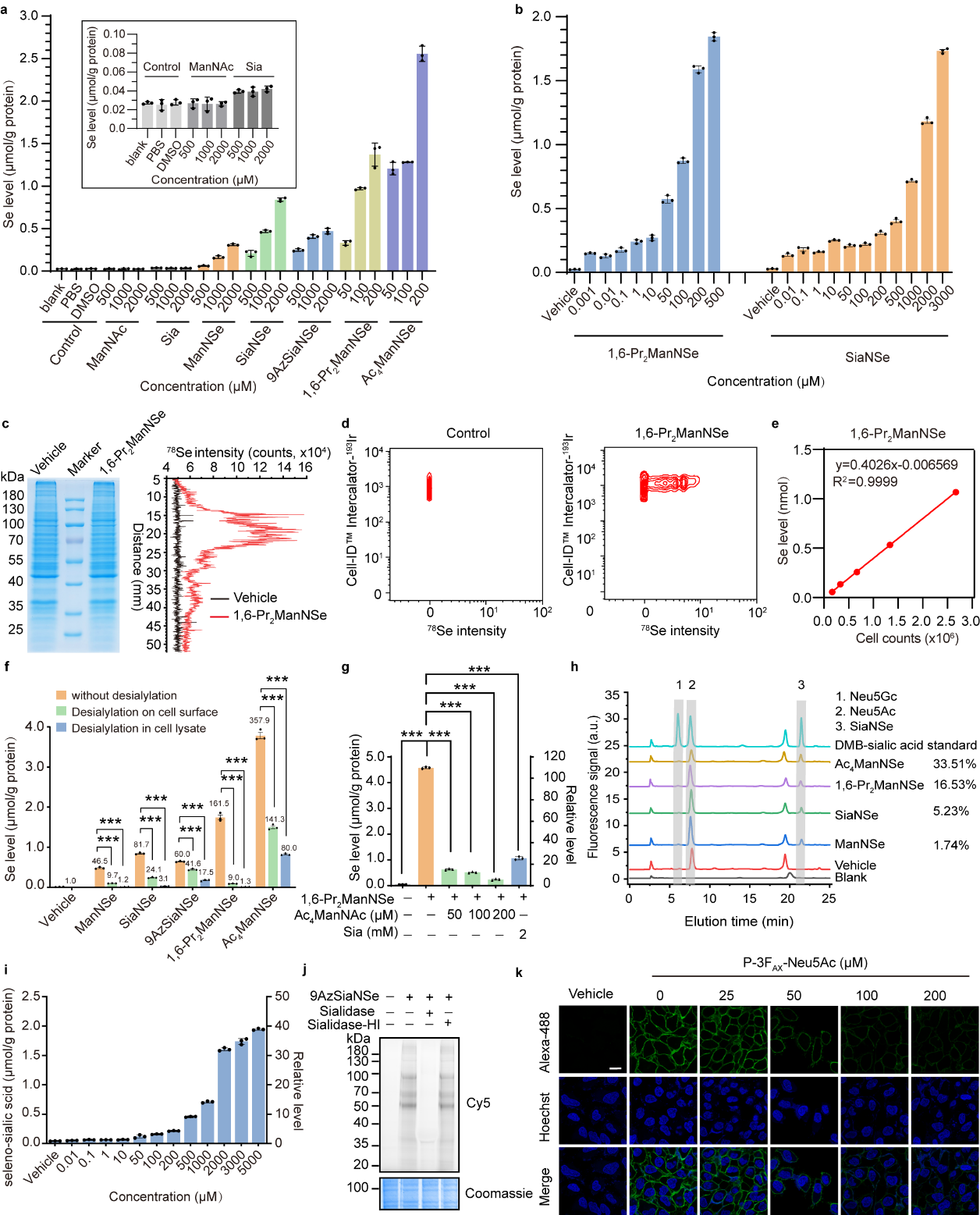
| Label-free glycan detection and dual-modality glycan perception by SeMOE. **a,** Se levels of glycoproteins from HeLa cells treated with respective monosaccharides for 48 h. **b,** Se levels of glycoproteins from HeLa cells treated with 1,6-Pr_2_ManNSe or SiaNSe at varied concentrations for 48 h. **c,** ^78^Se intensity analysis of HeLa cell proteins. HeLa cells were treated with 200 μM 1,6-Pr_2_ManNSe or vehicle for 48 h. Whole-cell proteins were resolved and transferred on PVDF membrane, and detected by LA-ICP-MS. **d,** CyTOF analysis of 293T cells treated with PBS or 200 μM 1,6-Pr_2_ManNSe for 48 h. Live cells were stained by Cell-ID^TM^ Intercalator-^193^Ir, and ^78^Se intensities were adopted. **e,** Linear correlation between cell number and Se level. **f,** Se levels of glycoproteins from respective selensugar-labeled HeLa cells treated with or without sialidase. **g,** Se levels of glycoproteins from HeLa cells treated with 200 μM 1,6-Pr_2_ManNSe and metabolic competitors (Ac_4_ManNAc or Sia) for 48 h. **h,** DMB-derived sialic acid analysis of glycoproteins from HeLa cells treated with indicated selenosugars for 48 h. Peak 1, 2 and 3 represent Neu5Gc, Neu5Ac and SiaNSe, respectively. The metabolic incorporation rate is calculated as the peak area ratio between peak 3 and the sum of peak 1, 2, and 3. **i,** Seleno-sialic acid levels of HeLa cells treated with vehicle or 9AzSiaNSe at varied concentrations. **j,** In-gel fluorescence scanning of 9AzSiaNSe-labeled HeLa cell proteins treated with sialidase or heat-inactivated sialidase (sialidase-HI). **k,** Confocal fluorescence imaging of HeLa cells treated with 9AzSiaNSe and the indicated concentrations of P-3F_AX_-Neu5Ac for 48 h. Scale bar:10 μm. All data are from at least three independent experiments. Error bars in all statistical data represent mean ± s.d. *P < 0.05, **P < 0.01, ***P < 0.001, ns, not significant (two-way ANOVA).

ICP-MS has now evolved into a versatile platform technology when integrated with technological advances such as cytometric system (*i.e.,* cytometry by time of flight, CyTOF)^14^ and laser ablation system (*i.e.,* laser ablation inductively coupled plasma mass spectrometry, LA-ICP-MS)^15^. Our hypothesis is that SeMOE probes are also compatible with these characterization approaches. To ascertain, whole-cell proteins treated with a panel of SeMOE probes were resolved on SDS-PAGE, transferred to PVDF membrane, and subjected to LA-ICP-MS linear scan analysis (Fig. 2c). Robust ^78^Se signals were observed in the protein bands when cells were treated with SeMOE probes, whereas only basal level of selenium was detected in the vehicle group (Fig. 2c). We confirmed that CyTOF can effectively and quantitatively measure the newly-synthesized selenium-containing glycome *in vitro* (Fig. 2d, **Supplementary Fig. 4**). Heretofore, all sialo-SeMOE probes demonstrated a linear correlation between cell number and total Se level (Fig. 2e, **Supplementary Fig. 5**).

Furthermore, we sought to validate that the Se signals indeed represent glycans under physiological conditions. To do so, new-to-nature sialo-SeMOE probes were added to HeLa cells. Treatment of Se-labeled cells with *Streptococcus pneumoniae* neuraminidase A (sialidase) significantly reduced the residence of Se signal on the cell-surface milieu (Fig. 2f)^16^. Similar results were obsereved for whole-cell proteins treated with sialidase (Fig. 2f). Alternatively, treatment of glycoproteins with PNGase F, which cleaves the asparagine side chain amide bond between glycoproteins and *N*-glycans^17^, revealed similar signal decrease (Extended Data Fig. 2a). Likewise, using of P-3F_AX_Neu5Ac, a cell-permeable metabolic inhibitor of sialic acid biosynthesis^18^, also reduced the signal in a dose-dependent manner (Extended Data Fig. 2b). We also confirmed the antagonization between sialo-SeMOE probes and their native or azido-functionalized counterparts (Fig. 2g, Extended Data Fig. 3). To quantitatively measure the metabolic conversion rate of sialo-SeMOE probes in glycoproteins, we performed sialic acid compositional analysis based on the fluorogenic 1,2-diamino-4,5-methylenedioxybenzene (DMB) derivatization. As expected, the SiaNSe peaks originated from cellular samples were in coincide with the synthetic standard substance, demonstrating that selenosugars, either biosynthetically converted from *N*-acetylmannosamine precursors or directly from sialic acid analogs, were metabolically incorporated into glycoproteins, and presented as one of the selenium speciations (Fig. 2h). The incorporation rate of Ac_4_ManNSe, 1,6-Pr_2_ManNSe, SiaNSe and ManNSe were 33.51%, 16.53%, 5.23% and 1.74%, respectively, in alignment with the observation in Fig. 2a. We also noticed a dose-dependent increase in other selenosugar incorporations by the DMB procedures (Extended Data Fig. 4).

**Fig. 3.**
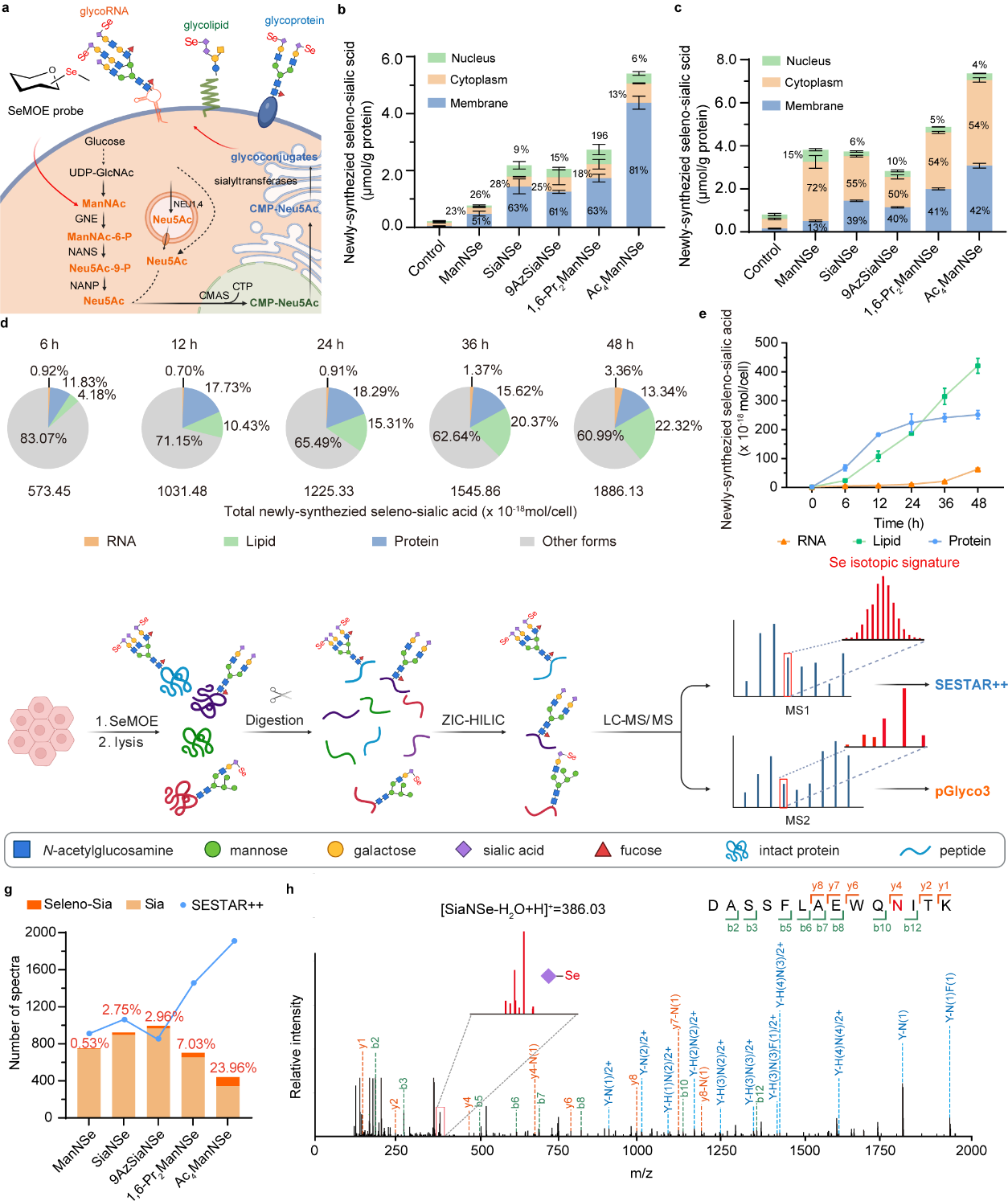
| Interpretation of the dynamic cellular fate of newly synthesized sialic acids by SeMOE. **a,** Schematic of the sialic acid biosynthesis pathway in cellular metabolism. **b-c,** Newly-synthesized seleno-sialic acid levels in different subcellular proteins **(b)** or subcellular fractions **(c)**. **d,** Newly-synthesized seleno-sialic acids in different glycoconjugates from HeLa cells treated with 200 μM 1,6-Pr_2_ManNSe for indicated time. **e,** Dynamic quantification of newly-synthesized seleno-sialic acids in glycoconjugates of HeLa cells treated with 200 μM 1,6-Pr_2_ManNSe for the indicated time. **f,** Workflow of intact *N*-glycoproteomics. *N*-glycopeptides from selenosugar-treated HeLa cells were enriched by ZIC-HILIC column and analyzed by LC-MS/MS. Raw data were analyzed by pGlyco3 or SESTAR++. **g,** Numbers of spectra (MS2) for glycopeptides containing natural sialic acid (Sia), or seleno-sialic acid (Seleno-Sia) identified by pGlyco3, and the number of spectra (MS1) for peptides matched in SESTAR++ searches. The percentage in orange represents the ratio between seleno-sialic acid-containing peptides and total sialopeptides, calculated according to spectra numbers. **h,** Example of a MS2 for the intact *N*-glycopeptide DASSFLAEWQNITK, showing a seleno-sialic acid with the Se isotopic signature. Hexose (H), HexNAc (N). All data are from at least three independent experiments. Error bars in all statistical data represent mean ± s.d. *P < 0.05, **P < 0.01, ***P < 0.001, ns, not significant (two-way ANOVA).

**Fig. 4.**
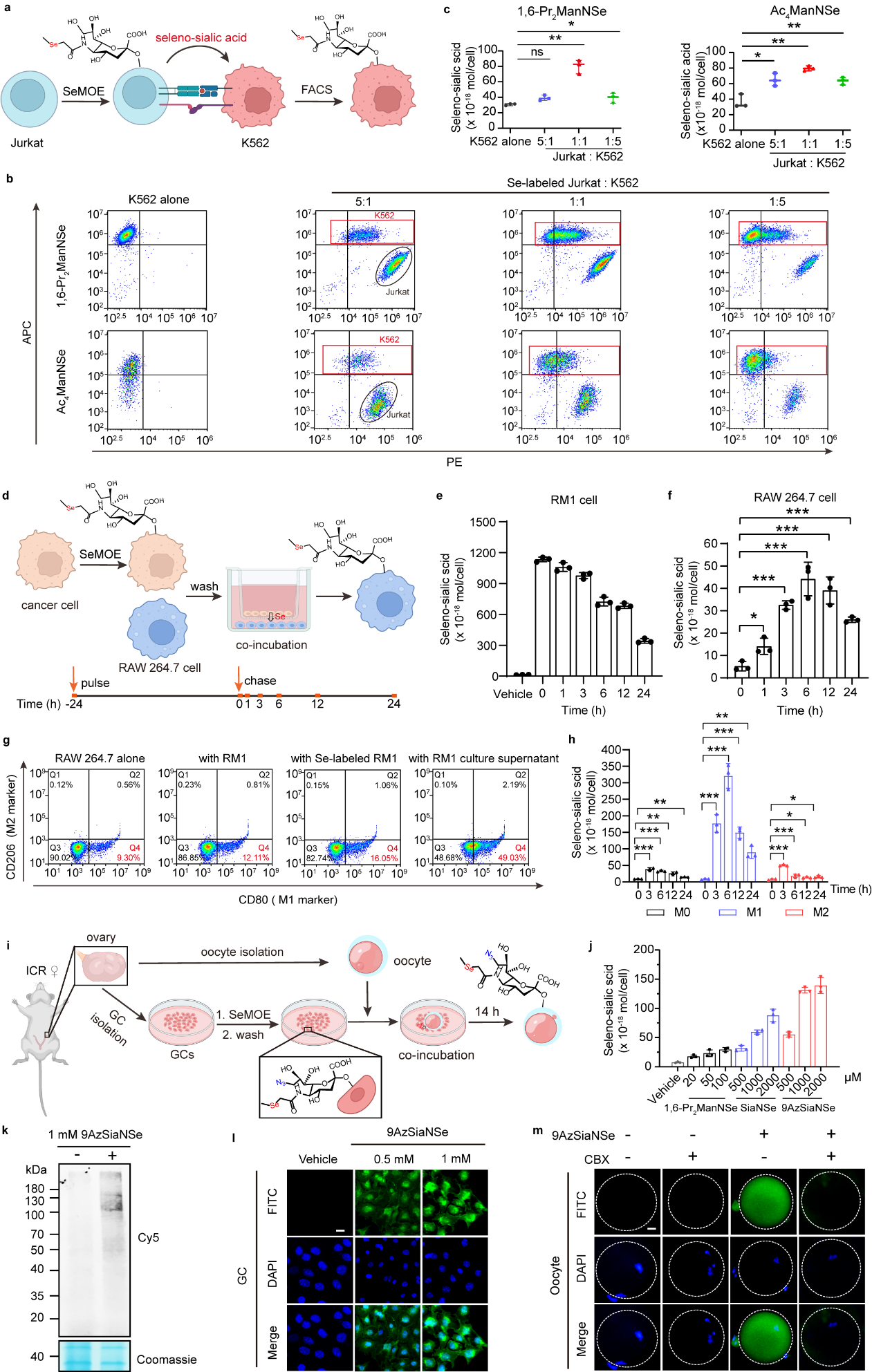
| Glycan transfer tracking in various cell-cell interactions. **a,** Schematic of glycan transfer during Jurkat-K562 trogocytosis. Selenosugar-treated Jurkat cells were stained by Dil-PE, washed and then co-incubated with Did-APC-stained K562 cells at 37℃ for 2 h, followed by sorting and ICP-MS analysis. **b,** Flow cytometry analysis of Jurkat-K562 trogocytosis. **c,** Seleno-sialic acid transfer from 1,6-Pr_2_ManNSe or Ac_4_ManNSe-treated Jurkat cells to K562 cells. **d.** Schematic of cancer cell-RAW 264.7 communication. Selenosugar-treated cancer cells were washed, and co-incubated with RAW 264.7 cells in a 0.4 μm-sized transwell culture system for varied time, followed by cell counting and ICP-MS analysis. **e,** Seleno-sialic acid levels of Se-labeled RM1 cells during co-incubation with RAW 264.7 cells. **f,** Seleno-sialic acid transfer from RM1 to RAW 264.7 cells. **g,** M1 polarization of RAW 264.7 cell induced by RM1 cell or RM1 cell culture supernatant. **h,** Seleno-sialic acid transfer from RM1 to RAW 264.7 cells in different polarization states. **i,** Schematic of GC labeling and co-incubation with oocytes. Selenosugar-treated freshly isolated GCs were washed, and co-incubated with oocytes for 14 h, followed by confocal imaging or ICP-MS analysis. **j,** Seleno-sialic acid levels of GCs treated with respective selenosugars at indicated concentrations for 48 h. **k,** In-gel fluorescence scanning of GCs treated with or without 9AzSiaNSe for 48 h, followed by reaction with alkyne-Cy5. **l,** Confocal fluorescence imaging of 9AzSiaNSe-labeled GCs. Scale bar: 20 μm. **m,** Confocal fluorescence imaging of oocytes after co-incubation with 9AzSiaNSe-labeled GCs with or without 100 μM CBX. Scale bar: 20 μm. All data are from at least three independent experiments. Error bars in all statistical data represent mean ± s.d. *P < 0.05, **P < 0.01, ***P < 0.001, ns, not significant (two-way ANOVA).

Previous reports revealed S-glyco-modification, an undesired side reaction when peracetylated unnatural azido hexoses were used in MOE^19^. Selective acylation on the C-1 and C-6-positions of azido hexoses effectively reduced such side reaction^20, 21^. Interestingly, the same rules applied for our chalcogen-containing unnatural sugars (**Supplementary Fig. 6)**. These data, collectively, prove that these label-free selenium-based carbohydrate probes can intercept the glycan biosynthetic pathway, faithfully depict and quantify the incorporated glycoconjugates, with ideal sensitivity and limit of detection.

## Dual-modality glycan perception with bifunctional SeMOE probes

Simultaneous integration of multimodality within one monosaccharide probe is the next research goal we want to pursue. We envisioned that the facile introduction of both bioorthogonal functional group (*i.e.,* azide) and ICP-MS responsive element (*i.e.,* selenium) would provide orthogonal signal readouts (fluorescence labeling vs. mass spectrometric analysis) for a more complex biological scenario. To synthesize, we installed azide at the C-9 position of a *N*-methylselanylacetyl sialic acid (SiaNSe), to generate bifunctional 9-azido *N*-methylselanylacetyl sialic acid (9AzSiaNSe)^22^ (Fig. 1b). Characterization of azide expression based on flow cytometry, confocal fluorescent microscopy and in-gel fluorescence showed that 9AzSiaNSe would tolerate the glycan biosynthetic pathway to be incorporated onto sialome, in a time-and dose-dependent manner (Extended Data Fig. 5a-e). Simultaneously, complementary detection of 9AzSiaNSe using the ICP-MS modality not only confirmed the concentration-dependent trend in its metabolism, but also provided meticulous and quantitative information at nanomolar to sub-micromolar concentration range (Fig. 2i). Cells treated with sialidase, sialic acid biosynthetic inhibitor, as well as a competition assay with native sialic acid, all strengthened the fact that 9AzSiaNSe indeed serve as a dual-modal sialic acid analog in SeMOE strategy (Fig. 2j, k, Extended Data Fig. 5f, g).

**Fig. 5.**
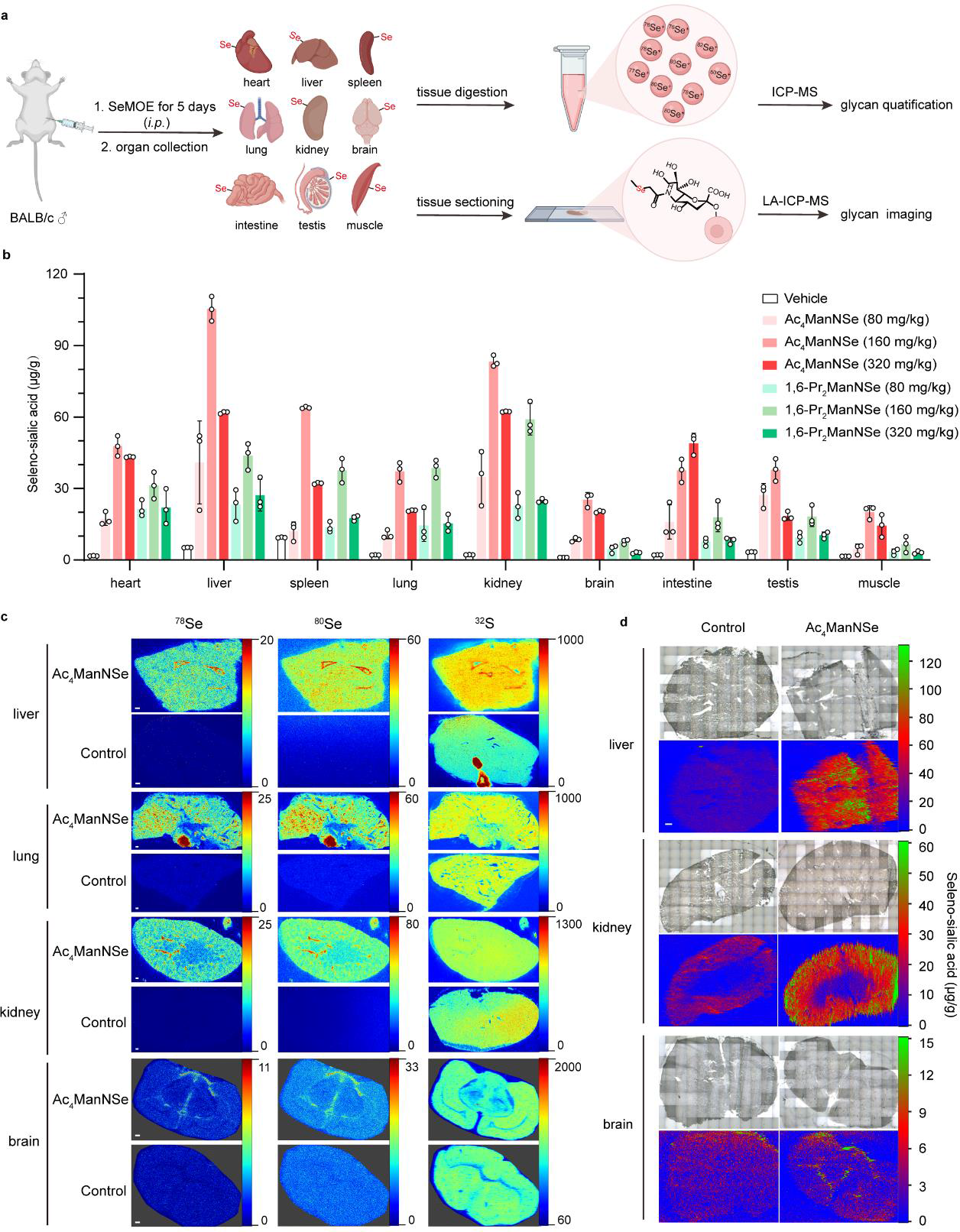
| In situ visualization and quantification of mouse tissue sialoglycans. **a,** Schematic of SeMOE *in vivo* and *in situ* tissue glycan imaging and quantification. Male BALB/c mice were once-daily, intraperitoneally injected with selenosugars at varied dosages, while control mice received 70% DMSO (vehicle) alone. On day 5, the mice were euthanized and perfused. The organs were digested for ICP-MS analysis, or sectioned for LA-ICP-MS imaging. **b,** Seleno-sialic acid levels of tissues from the mice treated with Ac_4_ManNSe or 1,6-Pr_2_ManNSe at the indicated concentration. **c,** In situ sialoglycan imaging of mouse liver, lung, kidney and brain by LA-ICP-MS. Scale bar: 200 μm. **d,** In situ sialoglycan quantification of mouse liver, kidney and brain by LA-ICP-MS. Scale bar: 200 μm. All data are from at least three independent experiments. Error bars in all statistical data represent mean ± s.d. *P < 0.05, **P < 0.01, ***P < 0.001, ns, not significant (two-way ANOVA).

## SeMOE probe quantitatively recapitulated the fate of sialic acid in cellular metabolism

In the Roseman-Warren-Bertozzi biosynthetic pathway, exogenous supplied sialic acid precursor or sialic acid, both in monosaccharide free form, were taken up by cells into the cytosol, converted to cytidine-5′-monophospho-*N*-acetylneuraminic acid (CMP-sialic acid) in the nucleus, trafficked into the Golgi apparatus, where they were utilized by sialyltransferases (Fig. 3a)^23^. Subsequent translocations of the sialylated glycoconjugates (*e.g.,* sialoprotein, sialolipid) were then accomplished via the secretory pathway, and ultimately displayed on the cell membrane. We reasoned that the quantitative feature of SeMOE would open up an avenue to delineate the biodistribution as well as the cellular fate of newly-synthesized sialoglycans. Therefore, we fractionated the major compartmentalization of HeLa cells treated with SeMOE probes after 48 h, and then determined the newly-synthesized sialic acid level in either precipitated proteins or isolated fraction buffer, using ICP-MS (Fig. 3b, c). The majority of newly-synthesized sialylated proteins were localized in the cell membrane (60%∼80%), showcasing the intrinsic function of sialoproteins as extracellular scaffolds (Fig. 3b)^24^. Signal resided in the nucleus (∼10%) and cytoplasmic matrix (∼20%) implied selenosugars bound to proteins vis glycosidic linkages may also present in these cellular regions (Fig. 3b), in accordance with the previous observation^25^. Notably, for the group without protein precipitation treatment, a large proportion of selenoglycan signal (50%-75%) was found in the cytosolic fraction rather than in the cell membrane, presumably due to the inclusion of selenosugars in free or nucleotide-activated form (Fig. 3c). Alternatively, we conducted a methodical evaluation on the cellular fate of selenosugars in newly-synthesized glycoconjugates, as embodied by glycoproteins, glycolipids, and glycoRNAs^26, 27^.

Major glycoconjugate components at various time points after SeMOE probe incubation were isolated for ICP-MS analysis. We monitored a gradual accumulation of selenium in glycoconjugates over time, with different time-dependent slope changes (Fig. 3d, **e**). Because substantial selenosugar signals were found in the metabolic flux pool, the selenoglycan substrates are thus not the limiting step (Fig. 3d). Hence, SeMOE quantitatively reflected the averaged sialylation turnover rate, and the absolute selenoglycan expression on major sialome classes.

By exploiting the isotopic signatures of certain elements, innovational techniques such as stable isotope labeling by/with amino acids in cell culture (SILAC)^28^ and isotope-targeted glycoproteomics (IsoTaG)^29^, have brought quantitative chemical glycoproteomics into the realm of possibility. Traditionally, bioorthogonal conjugation of isotope-containing affinity tags is used to realize the glycopeptide enrichment and isotopic mass pattern prediction^30^. The isotopic envelope pattern of selenium is guaranteed by its six stable isotopes with distinctive distribution (^74^Se, 0.89%; ^76^Se, 9.37%; ^77^Se, 7.63%; ^78^Se, 23.77%; ^80^Se, 49.61%; ^82^Se, 8.73%). We postulated that the effective incorporation of selenosugar in glycoproteins would achieve direct isotopic pattern prediction for glycopeptides, without the need for secondary tagging procedures. Consequently, we performed comparative studies using a previously reported computational algorithm, the updated version of selenium-encoded isotopic signature targeted profiling (SESTAR++)^31, 32^, and a glycan-first glycopeptide search engine, pGlyco3^33^ to comprehensively analyze the intact *N*-glycopeptides in cells treated with SeMOE probes (Fig. 3f, **Supplementary Table 5,6**). The substitution rate of selenosugars at a range of 0.53%-23.96% of total *N*-glycoprotein was observed, in concert with the DMB-labeled sialic acid analysis (Fig. 3g, Fig. 2h, **Supplementary Table 7**). Moreover, as expected, seleno-sialic acid peaks with Se isotopic signature in MS2 were found (Fig. 3h). However, owing to the microheterogeneity of *N*-glycan, the variation in glycan substitution rate, along with the intricate convolution of selenium isotopic distribution, the pattern recognition complexity in SESTAR++ will inevitably grow. Identified glycopeptides < 3 kDa exhibited expected the isotopic pattern in SESTAR++, while selenium-containing *N*-glycopeptides > 3 kDa showed a better match hit in pGlyco3 (**Supplementary Fig.7**). Nevertheless, the comprehensive utilization of SESTAR++ and pGlyco3 would mutually improve the detection of selenium-encoded glycopeptides in chemical glycoproteomics from complex samples.

*In toto*, SeMOE grants a pan-scale paradigm to monitor sialylation processes and the sialic acid metabolism.

## Tracking glycan transfer in cell-cell interaction and communication using SeMOE

Intercellular crosstalk and interaction orchestrate various biological processes such as organismal development and immune interaction, and are essential for cell viability^34^. Both contact-dependent communication (*e.g.,* via gap junction or cell-surface ligand-receptor interaction) and contact-independent communication (*e.g.,* via secreted molecules signaling) require diverse biomolecules to serve as the information carrier^35, 36^. However, the oligosaccharide moiety of glycoconjugates, or secreted sugar chains that participate in the cell-cell interaction are unconsciously overlooked, due to challenges in glycan characterization and quantification. To demonstrate the application of SeMOE as the evaluation platform for glycans upon cellular interaction, we selected two *in vitro* models as the exemplification: (a) contact-dependent trogocytosis assay^37^, and (b) contact-independent cancer cell-macrophage communication^38^. For the trogocytosis assay, we screened and selected 200 μM 1,6-Pr_2_ManNSe or 50 μM Ac_4_ManNSe as the ideal probe for T lymphocyte Jurkat cells (*i.e.,* effector cell) (**Supplementary Fig. 8**). SeMOE-treated Jurkat cells were co-incubated with K562 cells (*i.e.,* target cell) at 37℃ for 2 h, with different effector-to-target (E:T) ratios. K562 cells were sorted and the nibbled selenoglycans from Jurkat were then quantified using ICP-MS (Fig. 4a, **Supplementary Note 2**). A plethora of sialoglycoconjugates from Jurkat cells (1.6-4.5 femtomole selenosugar per Jurkat cell) were transferred to K562 cells (8.3-64.7 attomole selenosugar per K562 cell), with a glycoconjugate transfer percentage at a range from 0.1% to 3.1% as E:T ratio and selenosugar type varied (Fig. 4b,c, Extended Data Fig. 6a,b). A similar glycan exchange of incorporated 9AzSiaNSe between Jurkat and K562 cells was observed (Extended Data Fig. 6c-e).

**Fig. 6.**
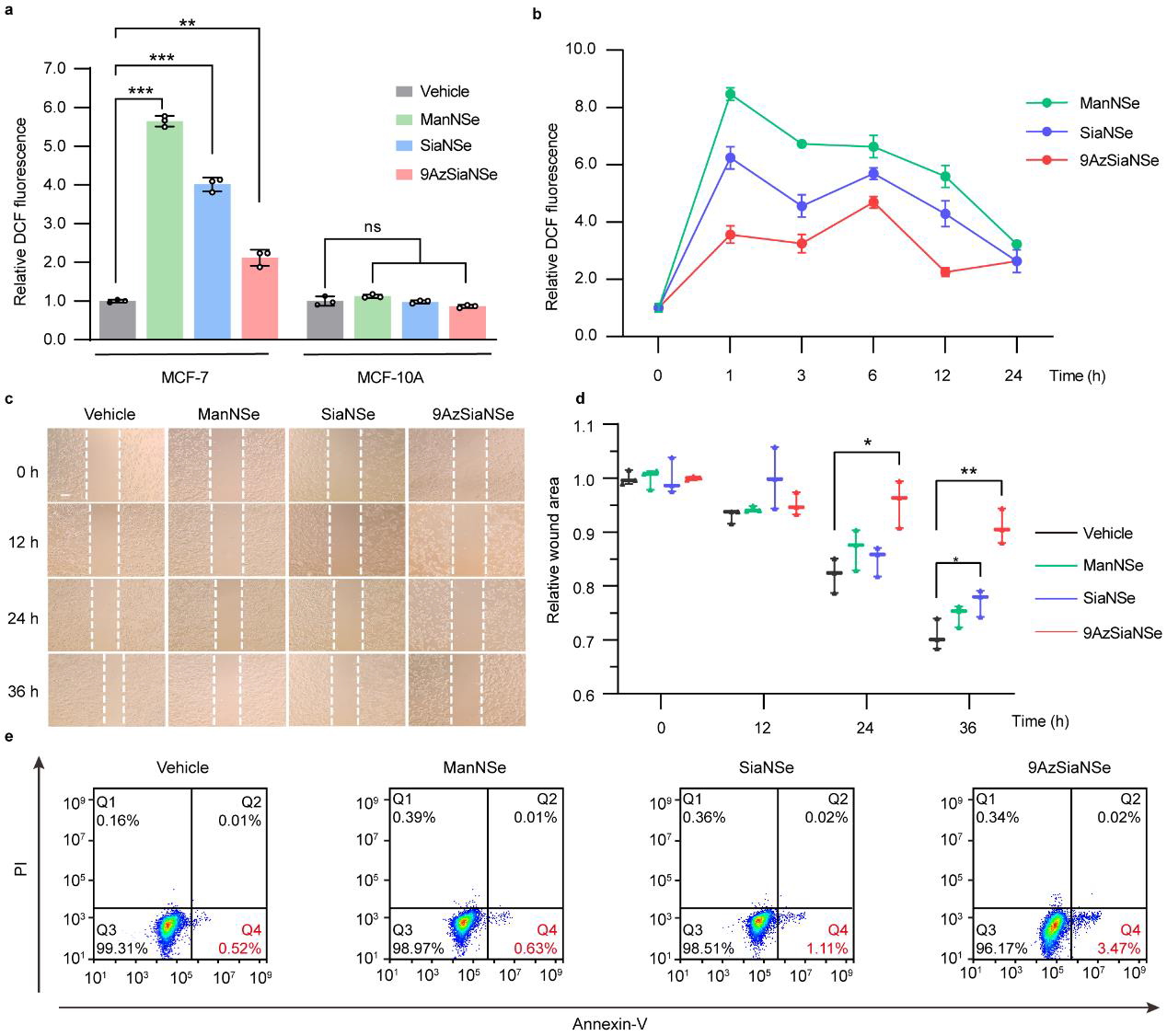
| Potential anticancer effect of the selenosugars. **a,** ROS levels of MCF-7 and MCF-10A cells treated with respective selenosugars at indicated concentrations for 24 h. **b,** ROS levels of MCF-7 cells treated with 10 μM ManNSe, SiaNSe or 9AzSiaNSe for varies time, respectively. **c,** Inhibitory effects of 1 mM ManNSe, SiaNSe or 9AzSiaNSe on MCF-7 cell migration. Scale bar: 100 µm. **d,** Relative wound area of MCF-7 cells of c. **e,** Flow cytometry analysis of MCF-7 cells treated with 1 mM ManNSe, SiaNSe, or 9AzSiaNSe by Annexin-V/PI assay. All data are from at least three independent experiments. Error bars in all statistical data represent mean ± s.d. *P < 0.05, **P < 0.01, ***P < 0.001, ns, not significant (two-way ANOVA).

For the cancer cell-macrophage communication studies, we adopted a transwell assay^39^ to investigate the glycosylated mediators between cancer cells and non-activated macrophage (M0) RAW 264.7 cells, using a pulse-chase experiment setting (Fig. 4d, **Supplementary Fig. 9**). A continuous glycan signal downfall was observed from RM1 cell (a carcinoma cell line) within 24 h, which was gradually channeled into seleno-sialoglycan signals observed in RAW 264.7 cells (Fig. 4e,f). We attribute the inflection point after 6 h to RAW 264.7 cellular division (Fig. 4f). Similar sialoglycan transfer trend was observed when 4T1 cell was used instead (**Supplementary Fig. 10**). We also demonstrated that participants in the indirect cellular communication were also *O*-glycosylated among a myriad of carcinoma cell lines when interacted with RAW 264.7 cells, and exhibited similar a fluctuation pattern (Extended Data Fig. 7). To examine whether selenoglycans would alter the macrophage state in response to indirect signaling, we characterized the polarization state of RAW 264.7 cells with or without selenosaccharide treatment on RM1 (Fig. 4g, Extended Data Fig. 8a). As a result, soluble signaling enacted by glycosylated, secreted molecules triggered the functional differentiation of macrophages from M0 to classically activated macrophages (M1)^40^, as evidenced by flow cytometric M1 biomarker staining and the reactive oxidation species (ROS) assay (Fig. 4g, Extended Fig. 8a,b). Reciprocally, we explored the incorporation of RM1-released, selenosugar-containing signal molecules in pre-induced macrophage phenotypes M0, M1, and alternatively activated macrophage (M2), corresponding to the normal, the pro-inflammatory, and the tumor-associated state of macrophages, respectively^41^. M1-type macrophage illustrated better signal molecule incorporation when compared to M2-type counterparts after proliferation normalization (Fig. 4h, Extended Data Fig. 8c), suggesting that selenosaccharides were readily poised as the signal molecule tracker in distinguishing macrophage phenotypes.

Furthermore, we applied SeMOE methodology in monitoring the sialic acid transfer between mouse primary granulosa cells (GCs) and oocytes. In mammals, oocyte maturation is closely related to its abundant *N*-glycosylation and the interaction with GCs^42^. Notably, nutrients such as glucose, pyruvate, nucleotides, and amino acids are transferred from GCs to oocytes through gap junctions^43^. Inhibition of gap junction or glycosylation is detrimental to oocyte maturation and GC development^44^. GCs were isolated from ovaries of 3-4 week ICR female mice, treated with 9AzSiaNSe, washed, co-incubated with oocytes for 14 h, and visualized or quantified using confocal fluorescent imaging or ICP-MS analysis (Fig. 4i). 9AzSiaNSe achieved dual-module characterization of GCs at both protein and cellular levels (Fig. 4j-l, **Supplementary Fig. 11a**). As expected, glycan transfer between GCs and oocytes was visualized by azide labeling (Fig. 4m, **Supplementary Fig. 11b**), manifesting the SeMOE applicability in primary cells. Unfortunately, we failed to quantify the absolute amount of glycans transferred into oocytes via ICP-MS mode, due to limitation in sample size. This is the first evidence that cell-surface sialoglycans participate in the substance exchange event in granulosa-oocyte interaction. Moreover, the treatment of gap junction inhibitor carbenoxolone (CBX) during co-incubation processes resulted in observable sialic acid transfer inhibition (Fig. 4m)^45^. The above-mentioned data conjointly proved the versatility and universality of the SeMOE strategy in interrogating cellular glycan exchange.

## SeMOE warrants quantitative in situ visualization of sialoglycans in biological tissues

We envisioned that coupling SeMOE technology with imaging mass spectrometric techniques such as LA-ICP-MS would fulfill in situ visualization of newly-synthesized glycan biodistribution in biological samples, with both high sensitivity and molecular specificity. Wild-type male BALB/c mice were intraperitoneally injected with Ac_4_ManNSe or 1,6-Pr_2_ManNSe at various dosages once daily for five days. On the sixth day, mice were euthanized, the organs were either homogenized for ICP-MS, or sectioned for LA-ICP-MS imaging (Fig. 5a). The seleno sialic acid incorporation in major organs exhibited a dosage-dependent manner, with the highest ingestion level of SiaNSe at 105.32 μg/g in the liver (Fig. 5b). Interestingly, organs such as brain also achieved distinguishable sialoglycan biodistribution, which are challenging to analyze using classical MOE strategy when blood-brain barrier (BBB) arises. Thin tissue sections on major organs were investigated with respect to the elemental distribution of ^78^Se, ^80^Se, ^32^S, ^56^Fe, ^63^Cu and ^66^Zn (Fig. 5c, Extended Data Fig. 9a,b). Strikingly, newly-synthesized sialoglycans demonstrated anatomical structure-specific biodistribution in major organs. For example, intensified selenium signals were found in the liver and lung lobes, the tracheal junction, surrounding tissues of the kidney, the contour of the thalamus, hippocampus and corpus callosum in the brain (Fig. 5c). In situ spatial imaging and absolute quantification of selenoglycan was realized and calibrated by using the Se standard substance (Fig. 5d, Extended Data Fig. 9c). To sum up, the label-free SeMOE, in tandem with LA-ICP-MS, effectively circumvent the secondary labeling used in classical MOE, and portrayed as an avant-garde tool to visualize and quantify newly-synthesized sialoglycans in animals with elemental precision.

## Potential anticancer effect of SeMOE probes

Selenium-containing entities including 1,4-anhydro-4-seleno-D-talitol (SeTal)^46^, selenadiazole derivatives (SeDs)^47^, and anti-inflammatory drug Ebselen provided therapeutic opportunities because of the unique redox property of selenium. Finally, we scrutinized the potential biological effects of SeMOE probes on cancer cells, in the aspect of ROS generation, cell migration and cell apoptosis. Intriguingly, SeMOE probes facilitated the rapid ROS generation in a variety of cancer cell lines rather than in normal cells (Fig. 6a, Extended Data Fig. 10a,b, **Supplementary Fig. 12**). To illustrate, MCF-7 cells treated with 10 μM ManNSe within 1 h already produced ROS escalation (Fig. 6b, Extended Data Fig. 10c). In addition, treatment of MCF-7 cells with ManNSe, SiaNSe or 9AzSiaNSe all inhibited the cell migration (Fig. 6c,d), and promoted cell early apoptosis in a dose-dependent manner (Fig. 6e, **Supplementary Fig. 13**). Overall, the precise function of SeMOE probes remains equivocal, and will be benefited from its mechanistic elucidation.

## Discussion

The core concept of SeMOE is minimalism. Revolutionary MOE has emerged as an organic chemistry-based methodology in the perturbation, profiling and perception of newly-synthesized glycans in living organisms. SeMOE developed in this work provides a simplified, yet versatile and convenient protocol for absolute quantification during glycan-mediated processes based on the feature of the non-metal element Se, with minimal background, attomolar detection of limitation for glycans, and excellent signal-to-noise contrast. Of note, SeMOE is applicable from the molecular to the tissue level, the isotopic pattern engraved in selenosugar shows the potential in developing the next-generation algorithm for glycoproteomic identification and analysis, while the development of LA-ICP-MS opens up a challenging path for multi-elemental imaging analysis with a subcellular resolution for a comprehensive view of element distribution over the whole organism. The interdisciplinary thought collision is vivified by the bifunctional seleno sialic acid, which enables simultaneous visualization and quantification of sialoglycans using different modalities. The information lost in static mass spectrometric analysis can be well compensated by the dynamic feature of fluorescence imaging or glycoprotein enrichiment. Furthermore, SeMOE strategy is readily expanded to other metabolic chemical reporters, including but not limited to glycans, lipids, proteins, and other post-translational modifications (*e.g.,* acetylation). Methodological adaption from chemoenzymatic glycan labeling or Se-click reaction will further augment the application of selenosugars in a constellation of biological scenarios. In addition, potential utilization of selenosugar as nutritional supplement, or regulatory agent in cancer therapy is of great interest. We believe the SeMOE strategy can serve as the Occam’s razor for glycan perception, to “See More” of the fascinating glyco-world.

## Acknowledgements

We thank Prof. Xing Chen (Peking University) for the guidance with the glycan labeling and glycoproteomics. We gratefully acknowledge support from the National Natural Science Foundation of China (No. 2207070006, No. 22107005), the Natural Science Foundation of Jiangsu Province (No. BK20202299), and the Programs for High-level Entrepreneurial and Innovative Talents Introduction of Jiangsu Province (Individual and Group Program), the Fundamental Research Funds for the Central Universities (No. 021414380508), the Chinese Academy of Sciences (No.YJKYYQ20210025), the STI2030-Major Projects (No. 2022ZD0211804), and National Key R&D Program of China (No. 2022YFA1505600)

## Author contributions

R.X. and M.W. supervised the project. R.X. and X.T. conceived the study, designed the experiments and analyzed the data. X.T. conducted most of the experiments unless specified otherwise. L.Z. performed the ICP-MS and LA-ICP-MS experiments under the supervision of M.W.. X.T., Y.H., B.C. and T.C. contributed to the chemical synthesis. X.T. and C.W. (in Nanjing University) performed the SeMOE *in vivo*. X.T., Y.H. and Y.L. performed the selenosugar cytotoxicity assay, cell subcellular fraction isolation and RNA (or lipid) sample preparation. X.T. and J.L. conducted the intact *N*-glycopeptide enrichment and pGlyco3 analysis. C.L., X.T. and C.W. (in Nanjing University) conducted the co-incubation of GC-oocyte under the supervision of L.D.. X.T. and G.J. performed the SESTAR++ searches under the supervision of C.W. (in Peking University). L.Y. and C.Z. contributed to the CyTOF analysis. R.X. and X.T. wrote the manuscript with contributions from all the authors.

## Methods

### Cell lines and mice

Cell lines HeLa, A549, 293T, Hep G2, SW620, 4T1, CT26, B16-F10, RAW 264.7, MCF-7, MCF 10A, Jurkat E6-1, K562, RM-1, T24, CHO, HMC3, MRC-5, SV-HUV-1 and HK-2 were all purchased from the American Type Culture Collection (ATCC). HCCC-9810 cells were purchased from the Institute of Biochemistry and Cell Biology, Shanghai Institutes for Biological Sciences, Chinese Academy of Sciences, Shanghai, China. HeLa, A549, 293T, Hep G2, SW620, 4T1, CT26, B16-F10 and RAW 264.7 cells were cultured in DMEM (Gibco) supplemented with 10% (v/v) fetal bovine serum (FBS) (Gibco), 100 U ml^−1^ penicillin and 100 μg ml^−1^ streptomycin (Gibco). MCF-7, MCF 10A, HCCC-9810, Jurkat E6-1, K562, RM-1 and T24 cells were cultured in RPMI 1640 (Gibco) supplemented with 10% (v/v) FBS (Gibco), 100 U ml^−1^ penicillin and 100 μg ml^−1^ streptomycin (Gibco). CHO, HMC3 and MRC-5 cells were cultured in α-MEM (Gibco) supplemented with 10% (v/v) FBS (Gibco), 100 U ml^−1^ penicillin and 100 μg ml^−1^ streptomycin (Gibco). SV-HUV-1 and HK-2 were cultured in DMEM/F-12 (Gibco) supplemented with 10% (v/v) FBS (Gibco), 100 U ml^−1^ penicillin and 100 μg ml^−1^ streptomycin (Gibco). Cultures were grown in T25 or T75 flasks (Thermo Fisher) and maintained at 37 °C with 5% CO _2_. Cells in all experiments were within 20 passages and free of mycoplasma contamination. Wild-type (WT) BALB/c and ICR mice were purchased from GemPharmatech Co. Ltd., Nanjing, China, and kept under specific pathogen-free (SPF) conditions with free access to standard food and water. All animal experiments were approved by the institutional Animal Care and Use Committee of Nanjing University.

### Antibodies and reagents

Antibodies included streptavidin-Alexa Fluor 488 (Thermo, S32354, 1:2,000), streptavidin-Alexa Fluor 555 (Thermo, S32355, 1:2,000), Hoechst 33342 (Thermo, S32354, 1:1,000), NucBlue™ Fixed Cell ReadyProbes™ Reagent (DAPI) (Thermo, R37606), HRP-conjugated anti-rabbit IgG(H+L) (Beyotime, A1092, 1:1000), Did-APC (Beyotime, C1039, 1 mM stock solution, 1:400), Dil-PE (Beyotime, C1991, 1:400), anti-mouse CD80-PE (Biolegend, 104707, 1:200), anti-mouse CD206-APC (Biolegend, 141707, 1:200). Alkyne-biotin (catalog no. 1137), alkyne AZDye 488 (catalog no. 1277), alkyne-Cy5 (catalog no. TA116), alkyne AZDye 647 (catalog no. 1301), and 2-(4-((bis((1-tert-butyl-1H-1,2,3-triazol-4-yl)methyl)amino)methyl)-1H-1,2,3-triazol-1-yl) acetic acid (BTTAA) (catalog no. 1236) were purchased from Click Chemistry Tools. Iodoacetamide and 4,5-methylenedioxy-1,2-phenylenediamine dihydrochloride (DMB) were purchased from Sigma Aldrich. P-3F_AX_-Neu5Ac (catalog no.5760) was purchased from Tocris Bioscience. LPS and IL-4 were purchased from Proteintech. Neuraminidase A (sialidase) from *Streptococcus pneumoniae* (NanA) was expressed and purified as previously described^16^. PNGase F was purchased from New England Biolabs (NEB). All organic agents were of at least analytic grade, obtained from commercial suppliers and used without further purification. *N*-acetylmannosamine (ManNAc), *N*-acetylneuraminic acid (Neu5Ac), *N*-glycolylneuraminic acid (Neu5Gc) and per−*O*−acetylated *N*-acetylmannosamine (Ac_4_ManNAc) were purchased from Carbosynth. *N*-azidoacetylmannosamine (ManNAz), *N*-azidoacetylneuraminic acid (SiaNAz), per−*O*−acetylated *N*- azidoacetylmannosamine (Ac_4_ManNAz), and 1,6-di-*O*-propionyl-*N*-azidoacetylmannosamine (1,6-Pr_2_ManNAz) were synthesized as reported^21, 49, 50^. Reagents for CyTOF analysis were purchased from Fluidigm. Dithiothreitol (DTT) used in LC–MS/MS analysis was purchased from J&K Scientific. Sequencing-grade modified trypsin and trypsin resuspension buffer were purchased from Promega. Se standard solution (100 ppm) was obtained from the National Sharing Platform for Reference Materials, China.

### SeMOE in vitro and in vivo

Ac_4_ManNSe and Ac_4_GalNSe were made to 200 mM stock solution (1,000 x) in sterile dimethyl sulfoxide (DMSO). 1,6-Pr_2_ManNSe and 1,6-Pr_2_GalNSe were made to 200 mM stock solution (1,000 ×) in sterile water. ManNSe, SiaNSe and 9AzSiaNSe were made to 500 mM stock solution (250 ×) in sterile water. All stock solutions were stored at −20℃.

In cell labeling experiments, the cells were treated with indicated unnatural selenosugars at varied concentrations for 48 h. The cells were collected by centrifugation (400 *g*, 5 min), washed three times with PBS, counted by the automated cell counter (Invitrogen), and then used for HNO_3_ digestion and ICP-MS analysis.

In the mouse metabolic labeling experiment, Ac_4_ManNSe or 1,6-Pr_2_ManNSe was diluted to 20 mg/mL in 70% DMSO (v/v) or in PBS, respectively. Male BALB/c mice (8 to 10 weeks old) were once-daily, intraperitoneally injected with 200 μL of Ac_4_ManNSe (160 mg selenosugar/kg/day), while control mice received the corresponding vehicle alone. On day 5, mice were euthanized. The mice were perfused with PBS, and major organs were collected and washed three times with PBS, frozen in liquid nitrogen-isopentane, and stored at −80°C until used for nitric acid digestion or tissue section preparation.

### Enzymatic treatment of live cells and protein samples

Neuraminidase A (sialidase) from *Streptococcus pneumoniae* (NanA), was expressed and purified as previously described^16^. Expressed protein was concentrated to 7.0 mg/mL. In the live cell experiment, 5 × 10^6^ HeLa cells were washed twice with PBS, treated with 10 mM ethylenediaminetetracetic acid disodium (EDTA Na2) in PBS, and centrifuged at 400 *g* for 4 min at room temperature, followed by three washes with PBS. Cells were resuspended in sialidase buffer (500 μL HBSS buffer, 10 μL sialidase and 2.5 μL of 1 M MgCl_2_) and incubated at 37℃ for 30 min.

In the protein lysate experiment, whole cell protein lysates were obtained as described above, and adjusted to 2 mg/mL using BCA assay. Protein samples were digested with sialidase or PNGase F, respectively. For sialidase-treated protein samples, 200 μg of glycoprotein (100 μL) from HeLa cells were mixed with 0.5 μL of 1 M MgCl_2_ and 10 μL sialidase, and reacted for 2 h at 37℃. For PNGase F treatment, 100 μg of glycoprotein (50 μL) from HeLa cells was mixed with 6 μL 10 × denaturing buffer and 4 μL deionized water, and denatured for 10 min at 100 ℃. Then, 10 μL GlycoBuffer 2 (10 ×) (NEB), 10 μL 10% NP-40 (NEB), 10 μL PNGase F (NEB) and 10 μL water were added into the reaction system. PNGase F cleavage occurred for 2 h at 37℃.

### Inhibition and metabolic competition assays

For inhibition of sialyltransferases, HeLa cells were treated with 0, 25, 50, 100 or 200 μM P-3F_AX_-Neu5Ac (Tocris), along with indicated concentration of unnatural selenosugar for 48 h. In the metabolic competition experiment, the natural monosaccharides, as well as their azido-counterparts, were used along with SeMOE probes for metabolic competition. HeLa cells were treated with these carbohydrate precursors/analogs at varied ratios as indicated for 48 h. The cells were subjected to cell fluorescence imaging or ICP-MS analysis afterwards.

### Isolation and validation of subcellular fractionation

The Membrane, Cytosol and Nuclear Protein Extraction Kit (KGBSP002, KeyGEN BioTECH) was used on selenosugar-labeled HeLa cells. After SeMOE, cells were washed three times with ice cold PBS, scraped off and pelleted. Cell pellets were suspended in Extraction Buffer A (supplemented with protease inhibitor) and lysed in a glass homogenizer, followed by incubation for 20 min at 4℃ and centrifugation (12,000 rpm, 10 min, 4℃). The supernatant was collected as cytoplasmic protein lysate. After washed twice with Extraction Buffer A, the remaining pellets were resuspended in Extraction Buffer B (supplemented with protease inhibitor), incubated for 10 min at 4℃ and centrifuged at 12,000 rpm for 10 min at 4 °C. This supernatant was collected as nuclear protein lysate. Then Extraction Buffer C (supplemented with protease inhibitor) was subsequently added to the remaining pellets, followed by incubation for 10 min at 4℃ and centrifugation (12,000 rpm, 10 min, 4℃). The supernatant was collected as membrane protein lysate. For analysis of selenoglycans covalently bound to proteins, lysates were precipitated by three volumes of alcohol at −80℃ overnight. The precipitated proteins were collected by centrifugation (12,000 *g*, 4 ℃, 10 min) and washed three times with 75% ethanol (v/v). For analysis of total selenoglycans, lysates were lyophilized. The above-mentioned subcellular fractions were digested in nitric acid and hydrogen peroxide (2:1, v/v) overnight, and analyzed by ICP-MS.

### Extraction of protein, lipid and RNA

MCF-7 cells were treated with or without 200 μM 1,6-Pr_2_ManNSe for varied time (6 h, 12 h, 24 h, 36 h, 48 h). Then, cells were collected by centrifugation (400 *g*, 5 min), washed three times with PBS, and counted by the automated cell counter (Invitrogen). All experiments were done with three replicates.

In protein isolation experiment, cells were suspended in cold RIPA buffer (1% Nonidet P-40 (v/v), 1% sodium deoxycholate (w/v), 150 mM NaCl, 0.1% SDS (w/v), 50 mM triethanolamine and EDTA-free Roche protease inhibitor, pH 7.4). After lysed by sonication, the cell lysates were centrifuged at 12,000 *g* for 10 min at 4°C for removal of debris, and the protein concentration was determined by BCA protein assay kit (Thermo Fisher Scientific). Then proteins were precipitated by adding into a MeOH/CHCl_3_ mixture (aqueous phase/MeOH/CHCl_3_ = 4:4:1, v/v/v), the isolated proteins were collected by centrifugation (20,000 *g*, 10 min, 4°C), and washed twice with cold methanol.

To extract cell lipids, 5 × 10^6^ cells were suspended in 150 μL cold water and lysed by sonication. Next, 400 µL methanol and 200 µL chloroform were added to the cell lysate and the mixture was stirred for 30 min at room temperature. After stirring, phase separation was initiated by adding 200 µL chloroform. The mixture was vortexed, and spinned at 2,800 *g* for 10 min at room temperature. The subnatant fraction (containing the lipid extracts) was carefully transferred into a fresh tube. Then the remaining mixture was mixed with 100 µL chloroform, stirred for 30 min at room temperature, and then centrifugated at 2,800 *g* for 10 min at room temperature. The subnatant fractions were collected and combined, and evaporated to dryness under N_2_.

RNA samples were isolated as previously described^27^. Samples were suspended in RNase-free water, and the RNA concentrations were determined by Nanodrop (Thermo Scientific). RNA samples were freeze-dried overnight. All samples isolated above were stored at −80°C until digestion by nitric acid for ICP-MS analysis.

### Acid digestion and ICP-MS quantitative analysis

After SeMOE treatment, the cells were collected, washed and counted as described above. For sample analysis of major biomacromolecules, protein, lipid and RNA extracts were collected as described above. To prepare acid digestion samples, 1 × 10^5^∼ 5 × 10^6^ cells, or 0.01∼1.0 mg biomacromolecule samples were mixed with 44 μL nitric acid and 22 μL 30% hydrogen peroxide, and digested for 24 h at room temperature. After digestion, Millipore water was added dropwise until a final nitric acid concentration of 2%, with 1.5 mL as the final volume.

For mouse organ samples, mouse organs were perfused, washed, frozen and stored as described above. Mouse organs were cut into pieces, lyophilized overnight and accurately weighed before mixed with 2 mL nitric acid and 1 mL 30% hydrogen peroxide. Samples were digested at 150℃ for ∼1 h in an airtight glass container until they became clear solutions. After that, digested solutions were heated at 100℃ to eliminate excess nitric acid, and 2% nitric acid was then added for samples to redissolve in 5.0 mL as the final volume. The sample solutions were stored at 4℃ until ICP-MS analysis. Se standard solutions with different concentrations (0, 1, 10, 50, 100, and 333 ppb), as well as sample solutions, were analyzed by solution nebulization ICP-MS (PerkinElmer, NexION 300D, USA). The parameters of ICP-MS solution analysis are shown in **Supplementary Table 1**. The isotopes ^78^Se were adopted for analysis, and the calibration curves of Se of ICP-MS solution analysis are shown in **Supplementary Fig. 2**. Se concentrations were absolutely quantified according to the standard curve. Se or selenoglycan levels were normalized according to the cell number or sample weight.

### CyTOF analysis

HEK293 cells were treated with vehicle or selenosugar (s) indicated (2 mM SiaNSe, 2 mM 9AzSiaNSe, 200 μM 1,6-Pr_2_ManNSe or 200 μM 1,6-Pr_2_GalNSe) at 37℃ for 48 h. After SeMOE, 3 × 10^6^ cells were collected and resuspended in 1 mL metal contaminant-free PBS (Rockland) supplemented with 0.5 μM cisplatin (Fluidigm), mixed immediately, and incubated at room temperature for 2 min, followed by centrifugation at 1,500 rpm for 5 min at room temperature. Cell pellets were washed twice with 2 mL CyFACS (Fluidigm), incubated with 125 nM Intercellular-Ir2 (Fluidigm) in FIX and PERM Buffer (Fluidigm) at 4℃ overnight, and then centrifuged at 2,000 rpm for 5 min and discard the supernatant. After that, cells were washed with 1 mL CyFACS and 1 mL Ultra-Water (Fluidigm). The cells were resuspended in Cell Acquisition Solution (Fluidigm) containing 10% EQ Four Calibration Beads (Fluidigm). Sample concentration was adjusted to acquire at a rate of 200 - 300 events/sec using a wide-bore (WB) injector on a CyTOF instrument (Fluidigm). The CyTOF data were exported as *.fcs files. Pre-gating was performed in FlowJo software (BD Biosciences) to filter the data to consist only of live, intact, single cells.

### DMB assay for sialic acid detection

Seleno-sialic acids and native sialic acids on protein were derivatized with 4,5-methylenedioxy-1,2-phenylenediamine dihydrochloride (DMB) and detected via reverse phase HPLC. In brief, HeLa cells were treated with or without selenosugars at indicated concentrations for 48 h. After SeMOE treatment, 2 × 10^6^ cells were washed three times with cold PBS, and centrifuged at 400 *g* for 10 min at 4℃. Cell pellets were resuspended in cold RIPA buffer (1% Nonidet P-40 (v/v), 1% sodium deoxycholate (w/v), 150 mM NaCl, 0.1% SDS (w/v), 50 mM triethanolamine and EDTA-free Roche protease inhibitor, pH 7.4) and incubated for 30 min at 4℃ to ensure sufficient lysis. Lysates were centrifuged at 12,000 *g* for 10 min at 4℃ to remove cell debris. Three volumes of ice-cold alcohol was added into the homogeneous cell lysates to precipitate the cellular proteins, and the mixture was kept at −80℃ overnight. Whole cell protein precipitates were collected by centrifugation (12,000 *g* for 10 min at 4℃) and washed four times with 75% alcohol (v/v) to discard the free sialic acids or CMP-sialic acids. Protein samples (or sialic acid standard) were dispersed in 2 M acetic acid and incubated for 3 h at 80℃ to release the protein-bonded sialic acids, cooled to room temperature, and filtered through 10 kDa MWCO filters (Millipore) by centrifugation (15,000 *g*, 15 min). The filtrate was collected and used for DMB derivatization directly. For sialic acid DMB derivatization, deionized H_2_O, DMB, β-mercaptoethanol and Na_2_S_2_O_4_ were added to make 100 μL as the final volume and adjust the final concentration of acetic acid, DMB, β-mercaptoethanol, Na_2_S_2_O_4_ at 1.4 mM, 7 mM, 0.75 M and 18 mM, respectively. Derivatization was performed in the dark at 50℃ for 2 h, cooled on ice for 10 min, and neutralized with NaOH solution (0.2 M, 25 μL). Samples were diluted 500 × with Millipore water and analyzed by RP-HPLC (Aglient 1260, XDB-C18 column, 5 µm, 4.6 × 250 mm) with a fluorescence detector (λ_ex_=373 nm, λ_em_ =448 nm). The flow rate was 0.8 mL/min and the elution gradient was: T (0 min) 84% H_2_O + 9% CH_3_CN + 7% CH_3_OH; T (14 min) 84% H_2_O + 9% CH_3_CN + 7% CH_3_OH; T (22 min) 64% H_2_O + 18% CH_3_CN + 18% CH_3_OH; T (28 min) 64% H_2_O + 18% CH_3_CN + 18% CH_3_OH; T (29 min) 84% H_2_O + 9% CH_3_CN + 7% CH_3_OH; T (30 min) 84% H_2_O + 9% CH_3_CN + 7% CH_3_OH. The incorporation efficiency of seleno-sialic acid into glycoproteins was quantified by the integration of peak areas.

### S-glyco-modification

HeLa cells were washed three times with cold PBS, lysed in cold PBS by sonication, and the debris was removed by centrifugation (20,000 *g*, 10 min, 4 °C). The total protein concentration was measured using the BCA protein assay kit (Thermo Fisher Scientific). 50 μL cell lysates (2 mg/mL) were incubated with indicated monosaccharides at varied concentrations at 37 °C for 2 h. After precipitation by adding 150 µL methanol, 37.5 µL chloroform and 100 µL Millipore water, the aqueous phase was removed by centrifugation (20,000 *g*, 5 min). The precipitated proteins were washed twice with cold methanol, stored at −80℃ until nitric acid digestion and ICP-MS analysis, or resuspended in 50 µL 0.4% SDS (w/v) in PBS for click reaction and in-gel fluorescence imaging.

### In-gel fluorescence scanning

In the click reaction setting of cell lysates, 100 µM alkyne-Cy5 or alkyne-Cy3, premixed BTTAA-CuSO_4_ complex solution (50 µM CuSO _4_, BTTAA/CuSO_4_ 2:1) and 2.5 mM freshly prepared sodium ascorbate were added to protein lysates (2 mg/mL in RIPA buffer), and vortexed at room temperature for 2 h. Cy5- or Cy3-labeled protein samples were resolved by SDS-PAGE, imaged by Typhoon FLA 9500 (GE) and analyzed by image Lab 3.0. The gels were stained by Coomassie Brilliant Blue (CBB) as the loading control.

### Cell confocal fluorescence microscopy imaging

Cells were seeded into Lab-Tek Chambered Coverglass (NUNC, 155409) and treated with vehicle or indicated metabolic precursors for 48 h. After metabolic labeling, cells were fixed with 4% formaldehyde (w/v) in PBS for 15 min, and washed three times with PBS. For click reaction, cells were incubated with 50 µM alkyne AZDye 488, premixed BTTAA-CuSO_4_ complex (50 µM CuSO _4_, BTTAA/CuSO_4_ 6:1) and 2.5 mM freshly prepared sodium ascorbate in PBS at room temperature for 10 min, followed by three washes with PBS. For nucleus staining, cells were incubated with 5 μg/mL Hoechst 33342 or DAPI at room temperature for 20 min, and washed three times with PBS. For cellular imaging, cells were imaged by Zeiss LSM 700 laser scanning confocal system equipped with a ×40 or ×63 oil immersion objective lens. FIJI/ImageJ (https://fiji.sc, NIH) was used for processing and visualization of all the images in this study.

### Flow cytometry analysis

Cells were seeded into 6-well plates and treated with vehicle or the metabolic precursors as indicated for 48 h. The cells were collected with 10 mM EDTA in PBS, transferred into 96-well tissue culture plates, centrifuged (400 *g*, 5 min, 4 °C) and washed three times with 1% FBS (v/v) in cold PBS. The cells were then resuspended in PBS containing 0.5% FBS (v/v), 50 µM alkyne AZDye 488, premixed BTTAA-CuSO_4_ complex (50 µM CuSO_4_, BTTAA/CuSO_4_ 6:1) and 2.5 mM freshly prepared sodium ascorbate, followed by reaction at 4℃ for 10 min. After three washes with 1% FBS (v/v) in cold PBS, cells were incubated with 1 µg/mL Streptavidin Alexa Fluor 488 conjugate at 4℃ for 30 min. The cells were washed three times with 1% FBS (v/v) in cold PBS, and applied to BD LSRFortessa Flow Cytometer system or ACEA NovoCyte benchtop flow cytometer (Acea Biosciences, USA). Flow cytometry data were analyzed using FlowJo.

### *N*-glycoproteomics and SESTAR++ analysis

For protein sample preparation, HeLa cells were treated with vehicle or indicated selenosugar (2 mM SiaNSe, 2 mM 9AzSiaNSe, 200 μM 1,6-Pr_2_ManNSe or 200 μM 1,6-Pr_2_GalNSe), collected, washed, re-suspended in 4% SDS (m/v) in PBS containing protease inhibitor, and lysed by sonication, followed by heating at 95℃ for 10 min and centrifugation (20,000 *g*, 10 min, 18℃) to remove debris. Protein concentrations were determined by BCA protein assay kit (Thermo Fisher Scientific). Then, cell proteins were precipitate by eight volumes of cold methanol at −80°C overnight. Precipitated proteins (1 mg) were denatured in 8 M urea, reduced in 10 mM dithiothreitol (DTT) at 37℃ for 1 h, alkylated in the dark by 20 mM iodoacetamide (IAA) at room temperature for 30 min, and diluted in 50 mM NH_4_HCO_3_ to a final urea concentration of 0.8 M. After that, trypsin (Promega) was added to a final enzyme-to-substrate ratio of 1:50 and incubated at 37 °C for 18 −20 h, followed by the centrifugation (20,000 *g*, 20 min). The supernatants were added 0.5% trifluoroacetic acid (v/v) to make the pH less than 3. All digested samples were desalted using the C18 column (Waters), dried by vacuum centrifugation and stored at −20 °C for further use. Then, glycopeptides were enriched using ZIC-HILIC (Merck Millipore). Briefly, desalted peptides (1-2 mg) were dissolved in 100 μL of 80% acetonitrile (v/v) containing 0.1% trifluoroacetic acid (v/v), loaded onto an in-house ZIC-HILIC micro-column containing 30 mg of ZIC-HILIC particles packed onto a C8 disk. The flow through was collected and passed back through the column for four additional times. Then, the column was washed seven times with 200 μL 80% acetonitrile (v/v) containing 0.1% trifluoroacetic acid (v/v) to remove hydrophobic peptides. Enriched glycopeptides were eluted twice with 70 μL 0.1% trifluoroacetic acid, collected, and dried by vacuum centrifugation.

For LC-MC/MS analysis, all above enriched glycopeptide samples were resuspended in 0.1% FA in water, separated over a 50 cm EasySpray reversed phase LC column (75 µm inner diameter packed with 2 μm, 100 Å, PepMap C18 particles, Thermo Fisher Scientific). The solvents (A: water with 0.1% formic acid and B: 80% acetonitrile with 0.1% formic acid) were driven and controlled by a Dionex Ultimate 3000 RPLC nano system (Thermo Fisher Scientific). The gradient was 6 h in total:1-7% solvent B in 10 min, followed by an increase from 7-35% solvent B from 11 to 311 min, an increase from 35-44% solvent B from 311 to 353 min, and an increase from 44% to 99% solvent B from 353 to 356 min. The MS parameters for glycopeptide analysis was: (1) MS: resolution = 120,000; AGC target = 500,000; maximum injection time = 50 ms; included charge state = 2–6; dynamic exclusion duration = 15 s; each selected precursor was subject to one sount HCD-MS/MS; (2) scout HCD-MS/MS: isolation window = 2; resolution = 15,000; AGC target = 500,000; maximum injection time = 250 ms; collision energy = 30%; stepped collision mode on, energy difference of ± 10%. RAW data of MS1 and MS2 were then searched by SESTAR++ or pGlyco3, respectively, as previously described^31, 33^ . Peptide spectrum matches in the pGlyco3 searches, MS1 matches in the SESTAR++ searches, and intact *N*-glycopeptides identified are shown in **Supplementary Tables 5-7**.

### Trogocytosis analysis of Jurkat cell and K562 cell

Jurkat cells were firstly treated with 50 μM Ac_4_ManNSe or 200 μM 1,6-Pr_2_ManNSe for 48 h. The cells were collected by centrifugation (400 *g*, 5 min), washed three times with 2% FBS (v/v) in cold PBS, and stained with Dil-PE staining buffer (5 μM Dil-PE in complete culture medium) in the dark for 10 min at room temperature, and washed three times with complete culture medium. Alternatively, Jurkat cells were treated with 1 mM 9AzSiaNSe for 48 h. The cells were collected by centrifugation (400 *g*, 5 min), washed three times with 2% FBS (v/v) in cold PBS, and followed by covalent attachment of biotin via click reaction. Briefly, the cells were resuspended at 5 × 10^6^ cells/mL in a click reaction buffer containing PBS, 0.5% FBS (v/v), 50 µM alkyne-biotin, premixed BTTAA-CuSO_4_ complex (50 µM CuSO _4_, BTTAA/CuSO_4_ 6:1) and 2.5 mM freshly prepared sodium ascorbate. The cells were reacted at 4℃ for 10 min, and washed three times with 2% FBS (v/v) in cold PBS.

Before co-culture experiment, K562 cells were stained with Did-APC staining buffer (5 μM Did-APC in complete culture medium) in the dark for 10 min at room temperature, and washed three times with complete culture medium.

For co-culture experiment, Dil-PE or biotinylated labeled Jurkat cells, were co-cultured with Did-APC labeled K562 cells (2 × 10^6^ cells/mL) in complete culture medium, respectively. Co-incubations were set up in 6-well U-bottom plates at varied cell number ratios (Jurkat:K562 at 5:1, 1:1, 1:5), with total cells at a cell number of 8 x 10^6^ per well. Cell mixtures were centrifuged at 150 *g* for 30 s to favor cell contact, and co-incubated for 2 h at 37℃. After co-incubation, cells were centrifuged (400 *g*, 4 min, 4℃), washed three times with 2% FBS (v/v) in cold PBS. Then, the co-incubation of Dil-PE labeled Jurkat cell and Dil-APC labeled K562 cells were directly used for flow cytometry analysis and sorting, while the co-culture of biotinylated Jurkat cells and Dil-APC labeled K562 cells were additionally incubated with 1 µg/mL streptavidin-Alexa Fluor 555 (PE) in the dark at 4℃ for 30 min. After three washes with 2% FBS (v/v) in cold PBS, cells were applied to for flow cytometry analysis and sorting. Flow cytometry analysis was conducted on a ACEA NovoCyte benchtop flow cytometer (Acea Biosciences, USA), and flow cytometry sorting was conducted on a BD FACSAria™ III Sorter. K562 cells were sorted in APC positive gate according to the flow cytometry gating strategy (**Supplementary Note 2**), accurately counted, digested with nitric acid, and analyzed by ICP-MS.

### Transwell assay of mouse cancer cells and RAW 264.7 cells

All transwell co-incubation experiments were performed in 0.4 μm-sized transwell inserts (Coring, USA). Before co-incubation, mouse cancer cells were treated with 200 μM 1,6-Pr_2_ManNSe or 1,6-Pr_2_GalNSe for 24 h in 6-well plates, while RAW 264.7 cells were incubated with or without 100 ng/mL LPS or 20 ng/mL IL-4 for 36 h in 6-well plates. SeMOE-treated cancer cells were collected, seeded into inner transwell inserts (1 × 10^5^ cells/insert), and incubated with 200 μM 1,6-Pr_2_ManNSe or 1,6-Pr_2_GalNSe overnight to guarantee SeMOE efficacy. In the meantime, RAW 264.7 cells were collected, seeded into 12-well plates (5 × 10^5^ cells/well), and incubated with or without 100 ng/mL LPS or 20 ng/mL IL-4 overnight to maintain the desired macrophage polarization status. After cell adherence, cancer cells and RAW 264.7 cells were washed three times with PBS, and co-cultured in a transwell setting at 37℃ for 24 h. Both cancer cells and RAW 264.7 cells were collected at various time points (1 h, 3 h, 6 h, 12 h, 24 h), washed three times with cold PBS, counted by the automated cell counter (Invitrogen), and subjected to macrophage polarization detection, ROS measurement or ICP-MS analysis.

For the effect of cancer cell-culture medium on macrophage polarization status, RM1 cell supernatant was collected, added to the culture medium of RAW 264.7 cells at a ratio of 50% or 100% (v/v), and incubated for 48 h. After that, RAW 264.7 cells were collected, washed three times with cold PBS, and used for macrophage polarization detection. RAW 264.7 cells were collected, washed three times with cold PBS containing 2% FBS (v/v), resuspended at 5 × 10^6^ cells/ mL in cold PBS containing 2% FBS (v/v), anti-mouse CD80-PE (Biolegend, 104707, 1:200) and anti-mouse CD206-APC (Biolegend, 141707, 1:200). After incubation in the dark at 4℃ for 30 min, cells were washed three times with 2% FBS (v/v) in cold PBS, and analyzed by NovoCyte benchtop flow cytometer in PE and APC channel.

### Isolation and co-culture of mouse GCs and oocytes

GCs and oocytes were isolated from ovaries of 3-4 week-old ICR female mice as previously described^51^. Briefly, mouse ovaries were collected, washed twice with complete DMEM/F-12 medium (Gibco) supplemented with 10% FBS (v/v), 100 U/mL of penicillin, 0.1 mg/mL of streptomycin, 1 mM pyruvate, and 2 mM glutamine, followed by removal of excess tissue with tweezers. For GC isolation, these ovaries were punctured with an 18-gauge needle to release GCs. The GCs were filtered through a 40 µm filter to remove the oocyte-cumulus complexes and cellular debris, collected into 15 mL centrifuge tubes (Corning, USA) and centrifuged at 1,000 rpm for 5 min. GC pellets were resuspended in complete DMEM/F-12 medium at a density of 5 × 10^5^ cells/mL, seeded in 6 cm Petri dishes, and cultured in complete DMEM/F-12 medium at 37℃, 5% CO_2_ with saturated humidity. The medium was replaced every 48 h of incubation until 80-90% confluency. To isolate oocytes, mouse ovaries were chopped with a blade to release oocytes. Denuded oocytes were obtained by a series of mouth-controlled micropipettes with successively smaller diameters, and cultured in droplets of M2 medium (Sigma) at 37°C in an incubation chamber (Billups Rothenberg, Del Mar, CA) infused with 5% O_2_ and 5% CO_2_.

For SeMOE of GCs, freshly isolated GCs were seeded into a 48-well plate with cell culture slides at a density of 5 × 10^5^ cells/mL and cultured in complete DMEM/F-12 medium for 48-72 h. Then, cells were washed twice with PBS, and treated with vehicle or the SeMOE probe indicated for 36-48 h, followed by three washes with PBS. GCs were fixed with 4% formaldehyde (w/v) in PBS for 20 min, washed three times with 0.5% BSA (m/v) in PBS, and incubated with 50 µM alkyne AZDye 488 or alkyne AzDye 647, BTTAA-CuSO_4_ complex (50 µM CuSO _4_, BTTAA/CuSO_4_ 6:1) and 2.5 mM freshly prepared sodium ascorbate in PBS for 10 min at room temperature. Then, GCs were incubated with DAPI for 10 min at room temperature, followed by three washes with 0.5% (m/v) BSA in PBS. For co-culture of GCs and oocytes, GCs were treated with 1 mM 9AzSiaNSe for 36-48 h, and washed three times with PBS. After that, freshly isolated mouse oocytes were added to wells, and co-cultured with 9AzSiaNSe-labeled GCs in M2 medium with or without 100 μM CBX for 14 h. Oocytes were collected by mouth-controlled micropipettes for further glycan confocal fluorescence microscopy imaging. Briefly, oocytes were fixed with 4% formaldehyde (w/v) in PBS for 20 min, washed three times with 0.5% BSA (m/v) in PBS, and permeabilized with 0.5% Triton X-100 (v/v) in PBS for 10 min. After three washes with 0.5% BSA (m/v) in PBS, oocytes were incubated with 50 µM alkyne AZDye 488, BTTAA-CuSO_4_ complex (50 µM CuSO _4_, BTTAA/CuSO_4_ 6:1) and 2.5 mM freshly prepared sodium ascorbate in PBS for 10 min at room temperature. After click reaction, oocytes were incubated with DAPI for 10 min at room temperature, followed by three washes with 0.5% (m/v) BSA in PBS. Fluorescence imaging was performed on Zeiss LSM 700 laser scanning confocal system equipped with a ×40 objective lens.

### Selenium analysis of proteins on the PVDF membrane by LA-ICP-MS

HeLa cells were treated with PBS or 200 μM 1,6-Pr_2_ManNSe for 48 h and lysed. Proteins were separated by SDS-PAGE and transferred to a PVDF membrane. The PVDF membrane was washed three times with deionized H_2_O, dried at room temperature, and subjected to LA-ICP-MS for linear scan analysis. The parameters and conditions of laser ablation system and ICP-MS are showed in **Supplementary Table 2**.

### In situ visualization and quantification of tissue sialoglycans by LA-ICP-MS

The organs of selenosugar-treated mice were collected, frozen in liquid nitrogen-isopentane, and sectioned to 10 μm slides for LA-ICP-MS imaging. For multi-element imaging, an Iridia laser ablation system (Teledyne Photon Machines, Bozeman, USA) with a 193 nm ArF excimer laser and a low-dispersion ablation cell was used. The LA system was coupled to icpTOF 2R ICP-TOFMS (TOFWERK AG, Thun, Switzerland) by the aerosol rapid introduction system (ARIS). High purity helium was used as transport gas to carry ablated sample aerosols from ablation chamber to ICP-TOFMS. For LA-ICP-TOFMS experiments, daily tuning was performed by using a NIST 612 glass standard RM (National Institute for Standards and Technology, Gaithersburg, USA) to obtain high sensitivity for ^59^Co, ^115^In, and ^238^U signal and a low oxide rate (*e.g.*, UO^+^/U^+^ < 3%). In order to obtain the shortest single pulse response, the flow of inner cell and outer cell was optimized. The size of laser spot was set as 20 μm, the distance between lines was set as 20 μm, the dosage was set as 1, the laser frequency was set as 250 Hz. The parameters and conditions of laser ablation system and ICP-TOFMS are showed in **Supplementary Table 3**. HDIP software (Teledyne Photon Machines, Bozeman, USA), TOFware, and laser image viewer software (TOFWERK AG, Thun, Switzerland) were used for imaging.

For quantitative imaging, the Iridia laser system was coupled to NexION 300D ICP-MS (PerkinElmer, Norwalk, USA) by the ARIS. Helium was used as transport gas. For LA-ICP-MS experiments, ICP-MS was tuned by using a NIST SRM612 glass for a maximum ^115^In and a low oxide rate (*e.g.*, UO^+^/U^+^). The isotopes ^78^Se was measured throughout all experiment. To ensure that tissue samples could be fully ablated, the size of laser spot was set as 20 μm, the distance between scanning lines was set as 40 μm. The optimized single pulse response was about 10 ms, the laser frequency was set as 100 Hz. The gelatin calibration standards containing 0, 1, 10, 50, 100 ppm selenium and tissue sialoglycans were analyzed together. The parameters and conditions of laser ablation system and ICP-MS are showed in **Supplementary Table 4**. Microsoft Office Excel 2020 and Origin Lab 2020 was used as data treatment software, and images integration were performed by the Iolite Software (v3.6) on Igor Pro 7 (WaveMetrics, USA).

### Measurement of intracellular ROS levels

The production of ROS induced by selenosugars was detected by a fluorescent ROS indicator, 2’,7’-dichlorofluorescein diacetate (DCFH-DA, ROS assay kit, Beyotime, S0033). Briefly, after SeMOE, cells were washed three times with PBS and incubated with 5 μM DCFH-DA in the dark for 30 min at 37℃. Then, cells were washed twice with PBS, trypsinized, followed by three washes with PBS. The level of intracellular ROS was determined by ACEA NovoCyte benchtop flow cytometer (Acea Biosciences, USA) in FITC channel.

### Cell proliferation assay

Cells were seeded in a 96-well plate (2,000 cells/well), treated with vehicle or selenosugar at varied concentrations for 48 h. The cells were washed three times with PBS and incubated with 100 μL medium containing 10 μL 2-(2-methoxy-4-nitrophenyl)-3-(4-nitrophenyl)-5-(2,4-disulfophenyl)-2H-tetrazolium monosodium salt solution (WST-8, Cell Counting Kit-8, Fdbio Science, FD3788) at 37 °C for 3 h. The absorbance at 450 nm was measured by a Synergy H4 Hybrid Reader (BioTek) and normalized to the vehicle-treated cells.

### Cell migration assay

MCF-7 cells were seeded in 6-well plates and incubated in 1640 medium overnight. The cells were collected, treated with vehicle or selenosugars as indicated for 36 h. The migrated cells were imaged by EVOS™ XL Core Imaging System (Thermo Fisher Scientific) at varied time. The inhibition of cell migration was analyzed by image J and normalized to the vehicle-treated cells.

### Cell apoptosis assay

The effect of selenosugars on cell apoptosis was detected by APC-Annexin V/PI Detection Kit (UElandy, A6030). In brief, after treated with vehicle or selenosugar at varied concentrations for 48 h, MCF-7 cells were collected and washed three times with 2% BSA (m/v) in PBS. 10 μL Binding buffer, 5 µL APC-Annexin V and 5 µL PI were sequentially added to 5 ×10^5^ cells, mixed and incubated in the dark for 15 min at room temperature, followed by the addition of 400 μL Binding buffer. Stained cells were analyzed immediately by ACEA NovoCyte benchtop flow cytometer (Acea Biosciences, USA) in FITC and APC channel.

### Reporting Summary

Further information on research design is available in the Nature Research Reporting Summary linked to this article.

## Extended Data Figures

**Extended Data Fig. 1.**
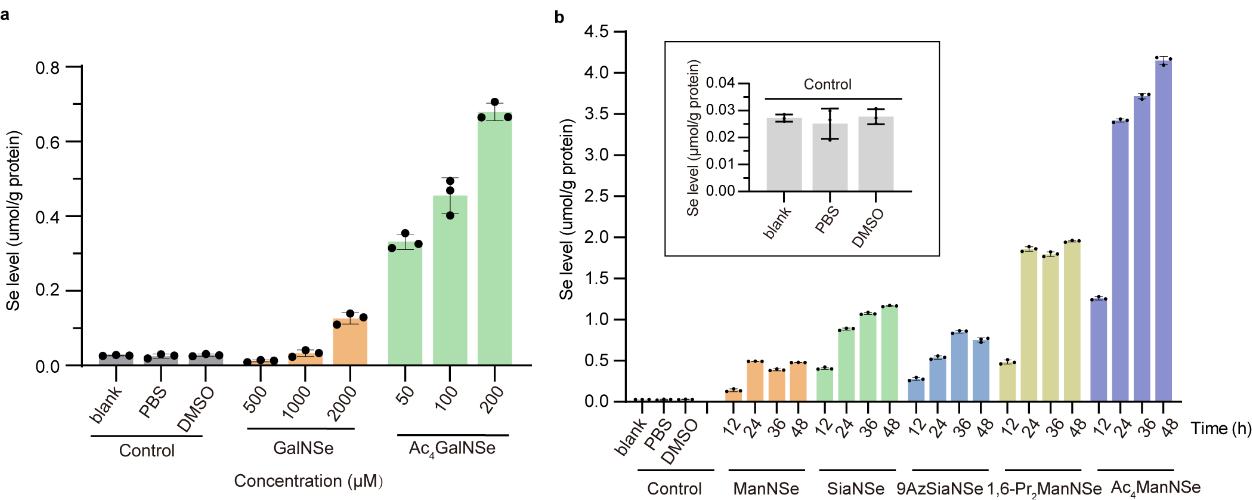
| Se levels of glycoproteins from HeLa cells treated with respective selenosugars. **a,** Se levels of glycoproteins from HeLa cells treated with vehicle, 2 mM GalNSe, or 200 μM Ac_4_GalNSe for the indicated time, respectively. **b,** Se levels of glycoproteins from HeLa cells treated with vehicle, 2 mM ManNSe, SiaNSe, 9AzSiaNSe, 200 μM 1,6-Pr_2_ManNSe or Ac_4_ManNSe for the indicated time, respectively. All data are from at least three independent experiments. Error bars in all statistical data represent mean ± s .d. (Related Fig. 2)

**Extended Data Fig. 2.**
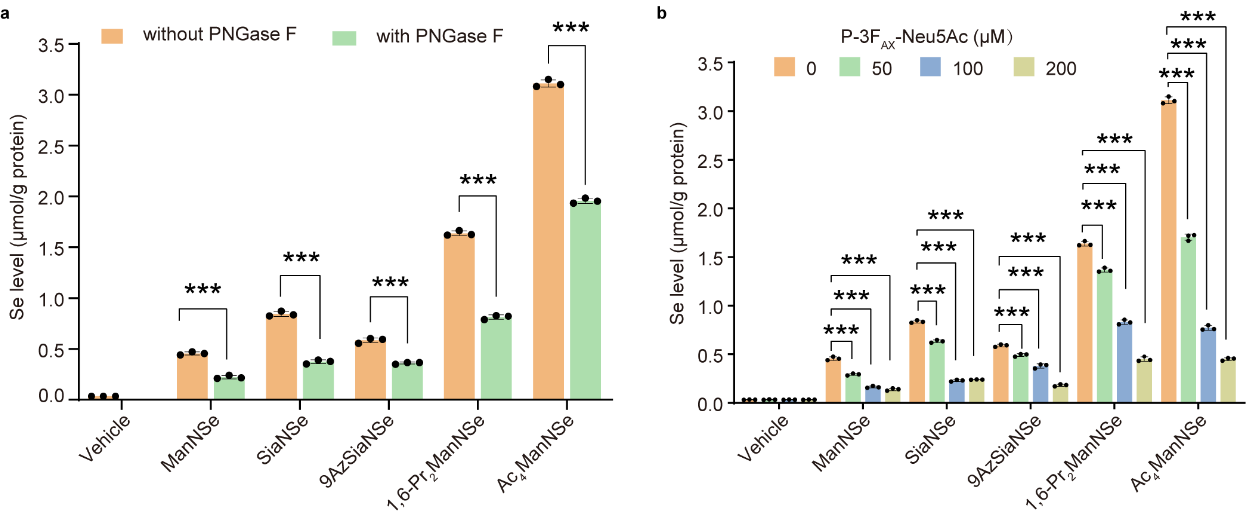
| Se levels of glycoproteins from respective selenosugar-labeled HeLa cells treated with PNGase F **(a)** or the indicated concentrations of P-3F_AX_-Neu5Ac **(b)**. (Related Fig. 2)

**Extended Data Fig. 3.**
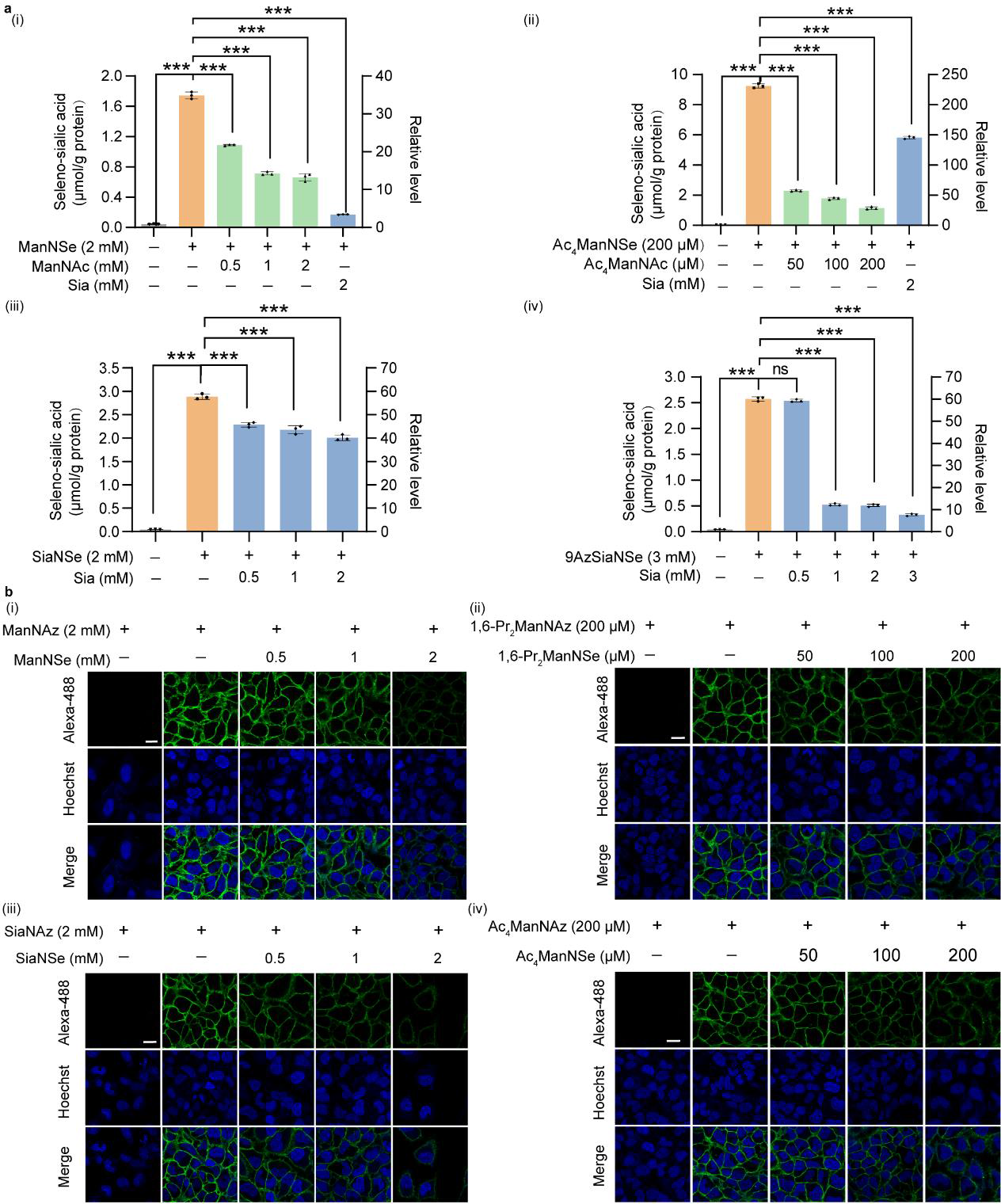
| Metabolic competition between selenosugars and known monosaccharides involved in sialic acid metabolism. **a,** Seleno-sialic acid levels of glycoproteins from Hela cells treated with indicated selenosugars and known monosaccharides involved in sialic acid metabolism for 48 h. **b,** Confocal fluorescence imaging of HeLa cells treated with indicate selenium-based and azide-based unnatural monosaccharides for 48 h. Scale bar: 10 μm. All data are from at least three independent experiments. Error bars in all statistical data represent mean ± s.d. *P < 0.05, **P < 0.01, ***P < 0.001, ns, not significant (two-way ANOVA). (Related Fig. 2)

**Extended Data Fig. 4.**
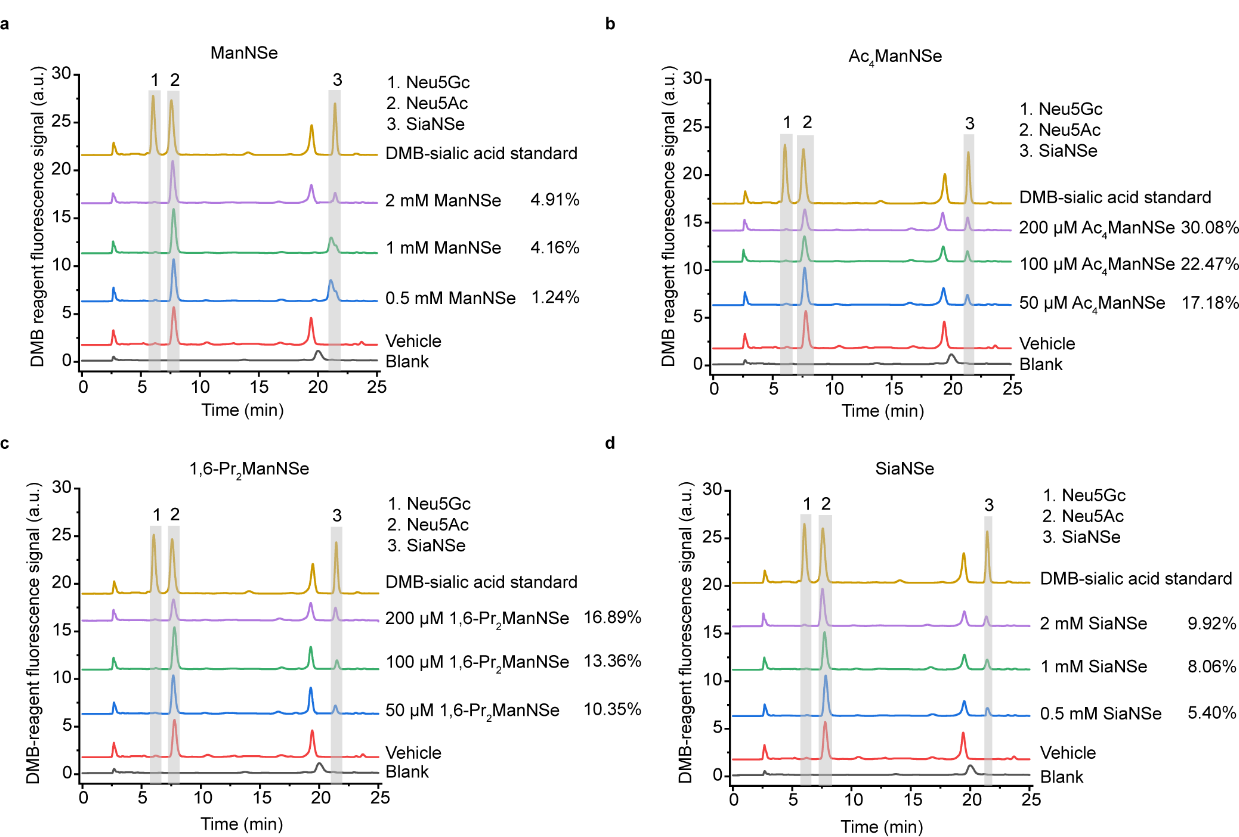
| DMB-derived sialic acid analysis of glycoproteins from HeLa cells treated with indicated selenosugars. **a-d,** Glycoproteins from HeLa cells treated with indicated selenosugars for 48 h were isolated, and reacted with the fluorogenic 1,2-diamino-4,5-methylenedioxybenzene (DMB) probe. HPLC analysis quantified the presence and abundance of specific sialic acids. Peak 1, 2 and 3 represent Neu5Gc, Neu5Ac and SiaNSe, respectively. The metabolic incorporation rate is calculated as the peak area ratio between peak 3 and the sum of peak 1, 2, and 3. (Related Fig. 2)

**Extended Data Fig. 5.**
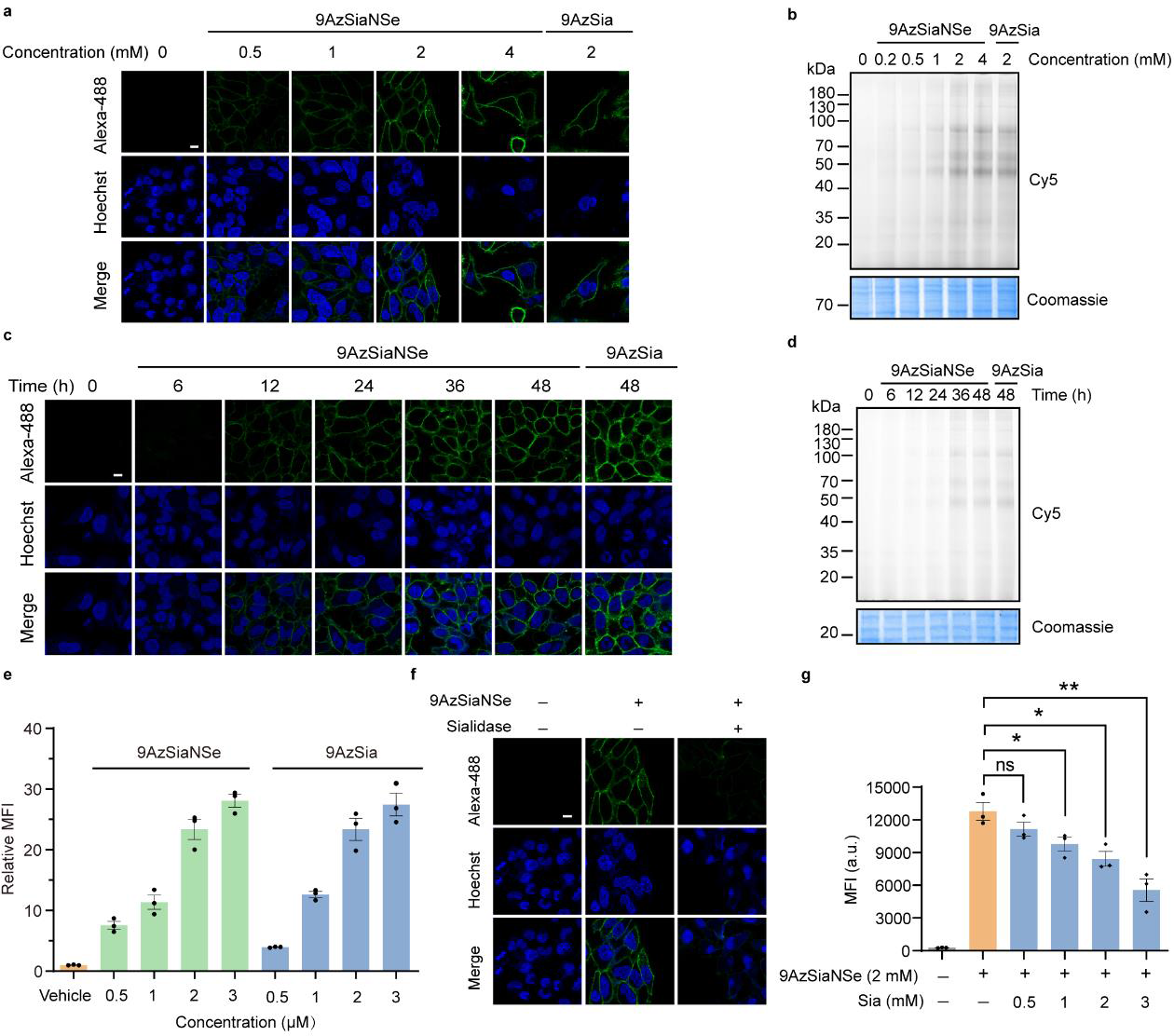
| Metabolic labeling of HeLa cells by 9AzSiaNSe. **a-b,** Confocal fluorescence imaging **(a)** or in-gel fluorescence scanning **(b)** of HeLa cells treated with vehicle or 9AzSiaNSe at indicated concentrations for 48 h, respectively. **c-d,** Confocal fluorescence imaging **(c)** or in-gel fluorescence scanning **(d)** of HeLa cells treated with vehicle or 2 mM 9AzSiaNSe for indicated time. **e,** Flow cytometry analysis of 9AzSiaNSe-treated HeLa cells. **f,** Confocal fluorescence imaging of 9AzSiaNSe-labeled Hela cells treated with or without sialidase, followed by reaction with alkyne-Alexa 488. Scale bar: 10 μm. **g,** Flow cytometry analysis of HeLa cells treated with 9AzSiaNSe and Sia at indicated concentrations for 48 h. All data are from at least three independent experiments. Error bars in all statistical data represent mean ± s.d. *P < 0.05, **P < 0.01, ***P < 0.001, ns, not significant (two-way ANOVA). (Related Fig. 2)

**Extended Data Fig. 6.**
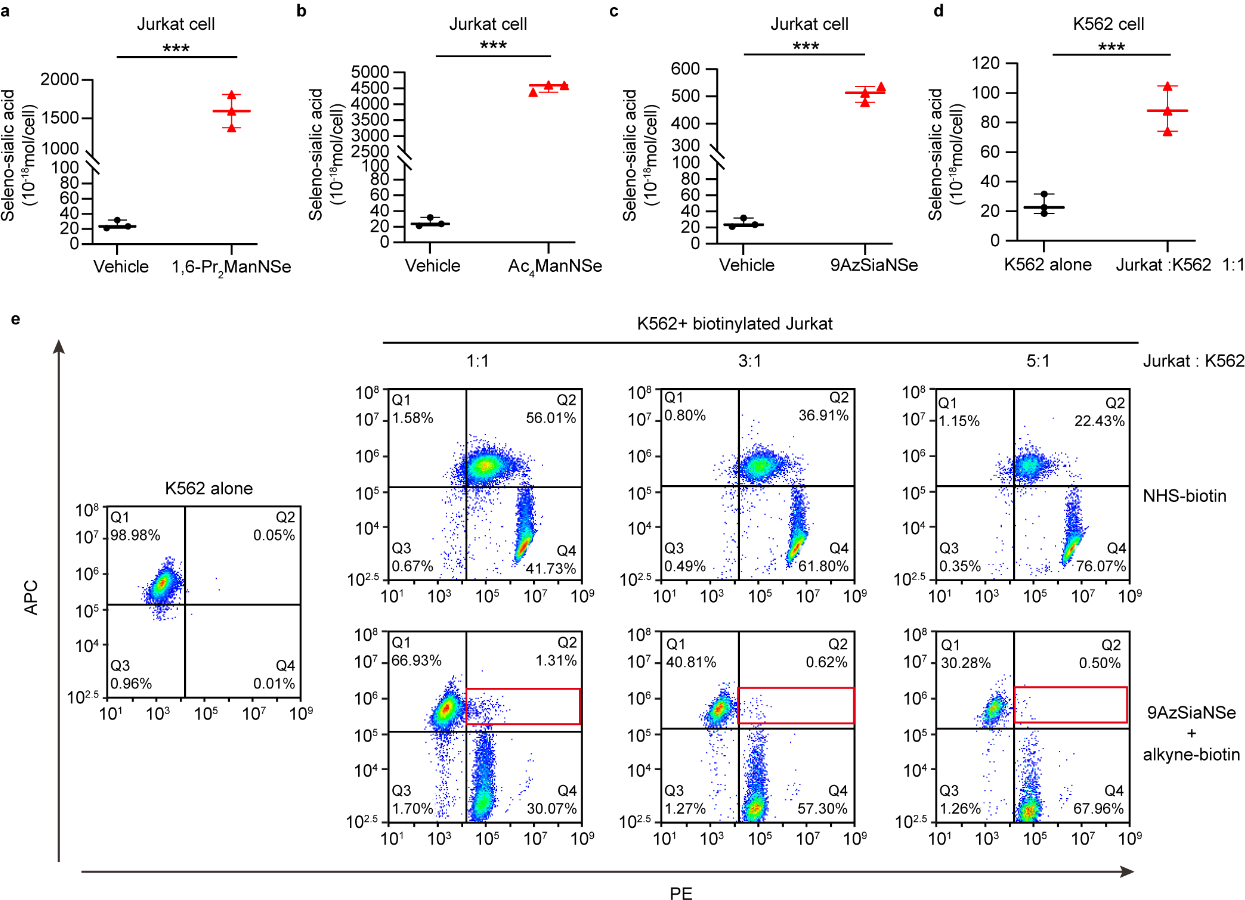
| Glycan transfer quantification during Jurkat cell-K562 cell trogocytosis. **a-b,** Seleno-sialic acid levels of Ac_4_ManNSe-treated **(a)** or 1,6-Pr_2_ManNSe-treated **(b)** Jurkat cells. **c,** Seleno-sialic acid levels of 9AzSiaNSe-treated Jurkat cells. **d,** Seleno-sialic acid transfer from 9AzSiaNSe-labeled Jurkat cells to K562 cells during trogocytosis. **e,** Flow cytometry analysis of protein and sialic acid transfer during Jurkat and K562 cell trogocytosis. Jurkat cells treated with or without 9AzSiaNSe were biotinylated by NHS-biotin or alkyne-biotin, respectively. Did-APC stained K562 cells were co-incubated with biotinylated Jurkat cells in varied ratios at 37℃ for 2 h, followed by incubation with streptavidin-PE at 4℃ for 0.5 min. All data are from at least three independent experiments. Error bars in all statistical data represent mean ± s.d. *P < 0.05, **P < 0.01, ***P < 0.001, ns, not significant (two-way ANOVA). (Related Fig. 4)

**Extended Data Fig. 7.**
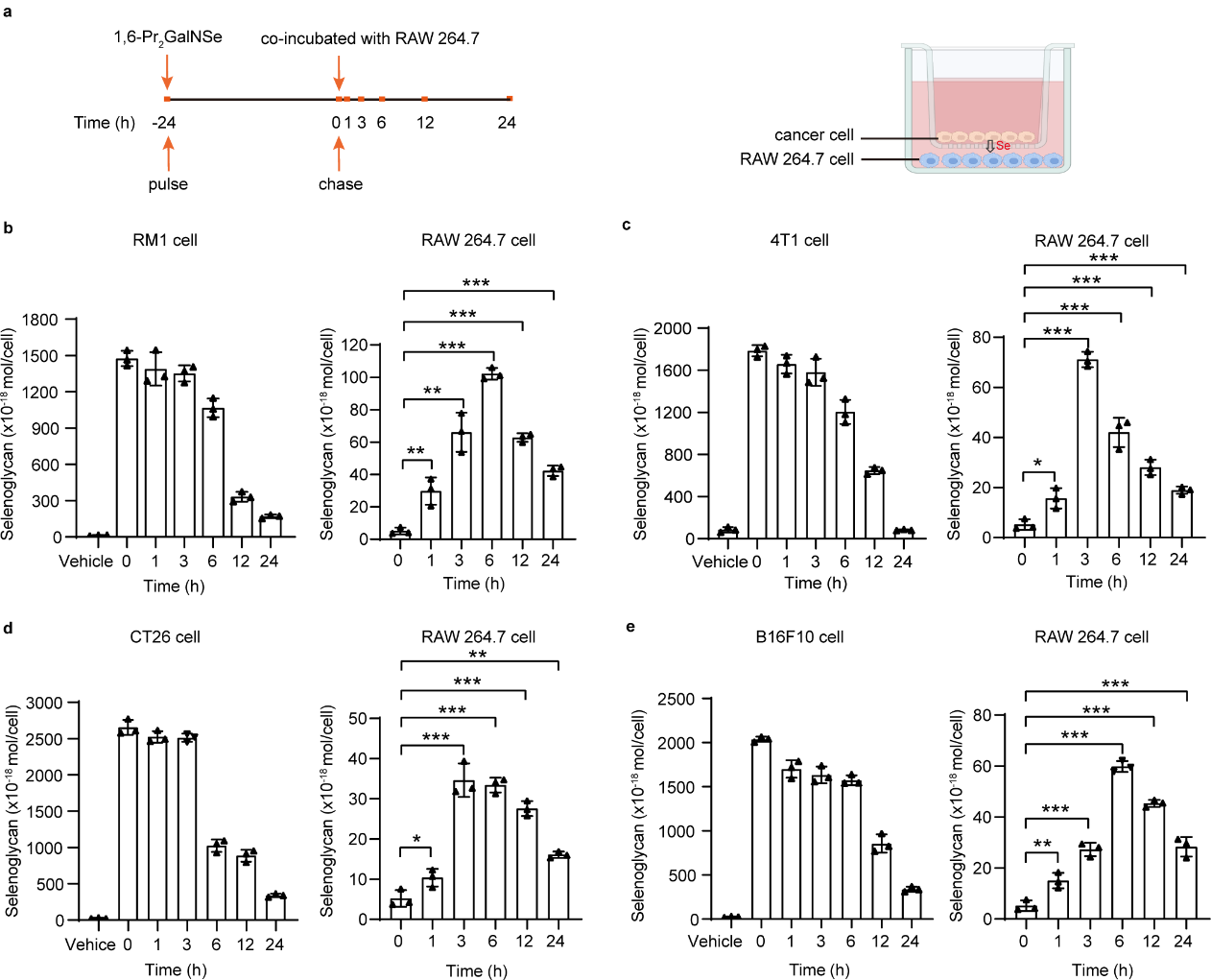
| Glycan transfer during cancer cell and RAW 264.7 cell transwell incubation. **a,** Workflow of the co-incubation of 1,6-Pr_2_GalNSe-treated cancer cells and RAW 264.7 cells. Cancer cells were treated with 200 μM 1,6-Pr_2_GalNSe for 24 h, washed, and then co-incubated with RAW 264.7 cells in a 0.4 μm-sized transwell culture system for varied time. **b-e,** Selenoglycan levels of RM1 cells. **(b)**, 4T1 cells **(c)**, CT26 cells **(d)** and B16F10 cells **(e)** during incubation with RAW 264.7 cells, and selenoglycan transfer from cancer cells to RAW 264.7 cells. All data are from at least three independent experiments. Error bars in all statistical data represent mean ± s.d. *P < 0.05, **P < 0.01, ***P < 0.001, ns, not significant (two-way ANOVA). (Related Fig. 4)

**Extended Data Fig. 8.**
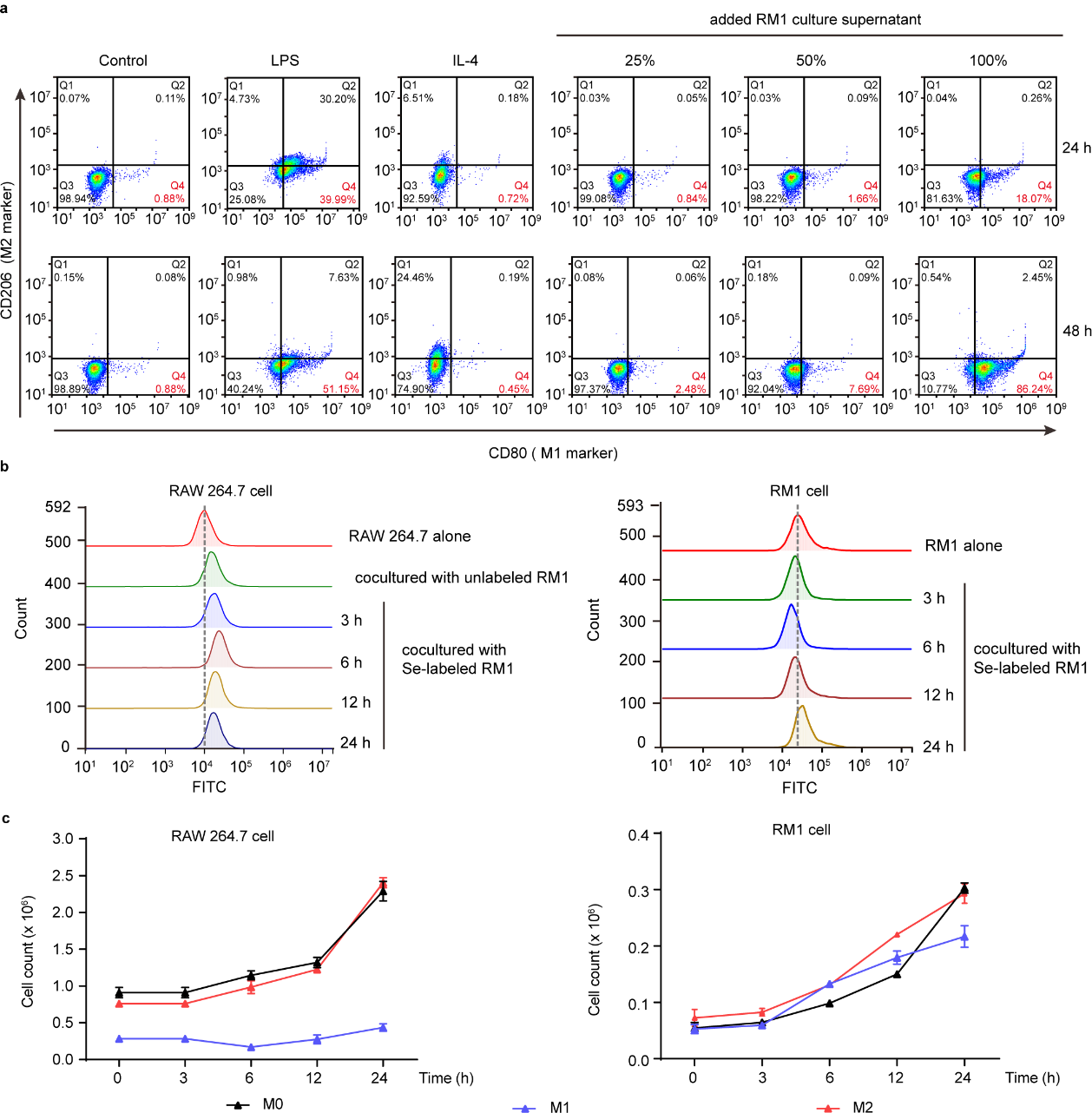
| Biological effects of co-incubation of cancer cells and RAW 264.7 cells. **a,** RM1 cell culture supernatant-induced M1 polarization of RAW 264.7 cells. **b,** ROS assay of RAW 264.7 cells and RM1 cells during co-incubation. **c,** Cell counts of RAW 264.7 cells in different polarization states and RM1 cells during co-incubation. All data are from at least three independent experiments. Error bars in all statistical data represent mean ± s.d. (Related Fig. 4)

**Extended Data Fig. 9.**
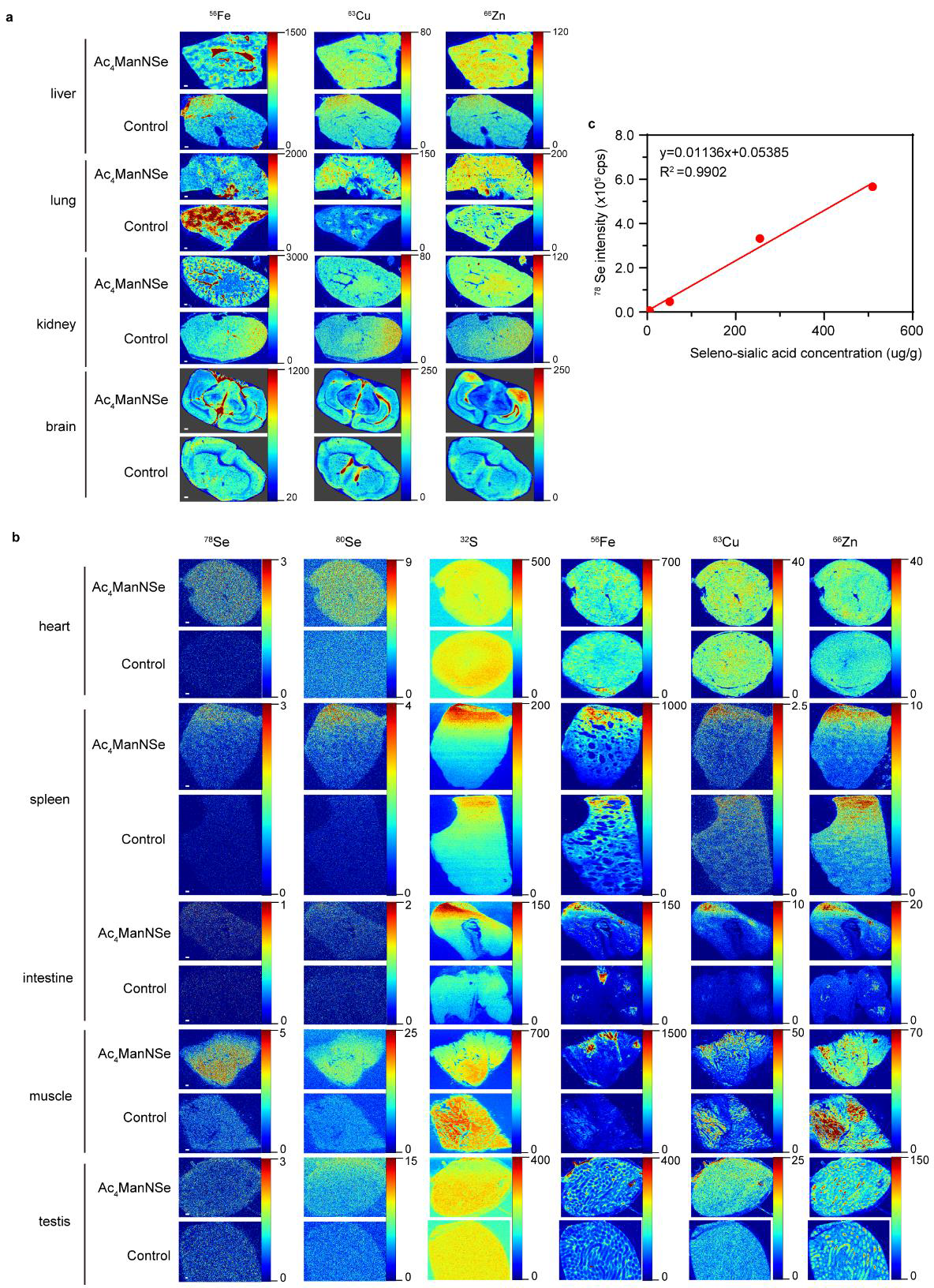
| In situ visualization of sialoglycans in mouse tissues. **a,** In situ LA-ICP-MS imaging of ^56^Fe, ^63^Cu, and ^66^Zn in mouse liver, lung, kidney and brain. **b,** In situ LA-ICP-MS imaging of sialoglycans in mouse heart, spleen, intestine, muscle and testis. Scale bar: 200 μm. **c,** Standard curve for in situ quantification of sialoglycans by LA-ICP-MS. The calibration curve was constructed by gelatin slices containing 0, 1, 10, 50, 100 ppm selenium, respectively, against ^78^Se-signal intensity, on the LA-ICP-MS. (Related Fig. 5)

**Extended Data Fig. 10.**
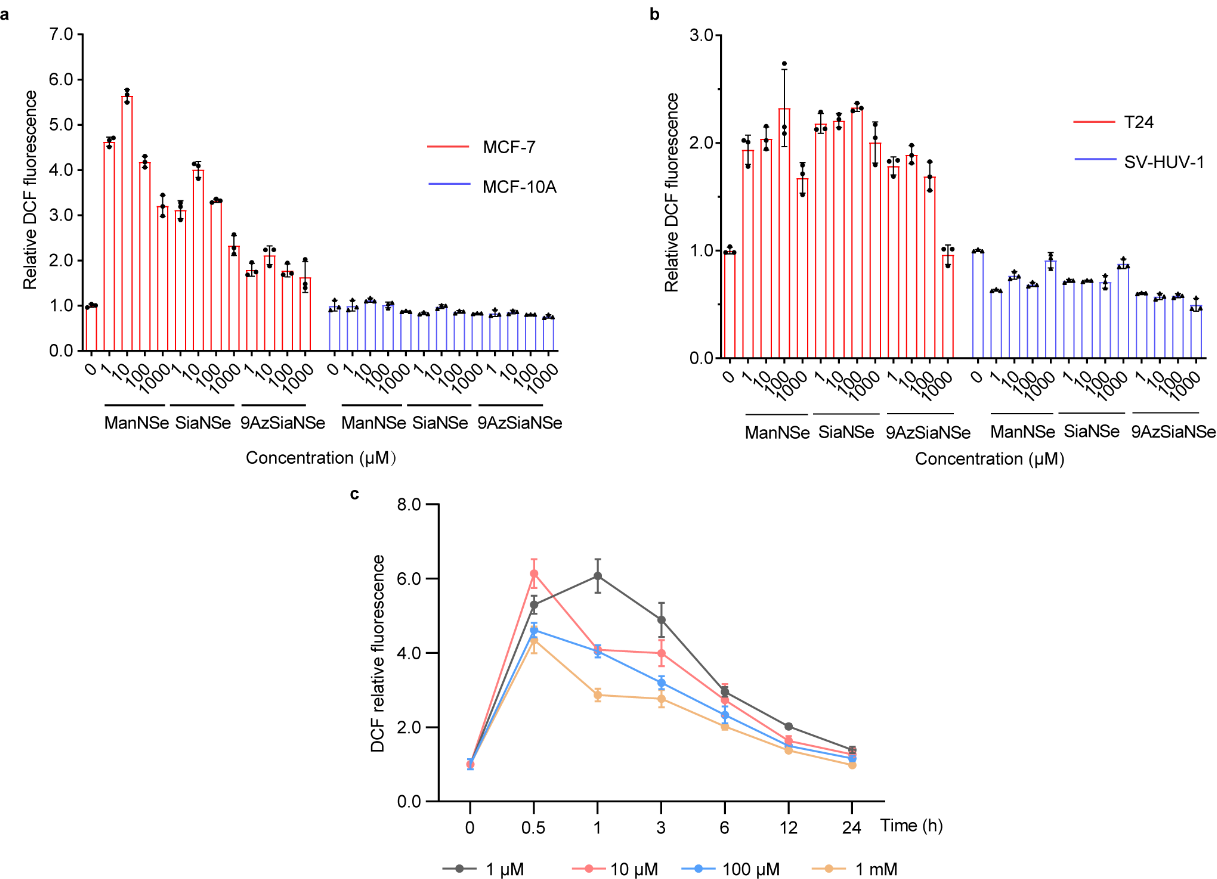
| ROS assay of respective selenosugar-treated cells. **a,** ROS assay of MCF-7 and MCF-10A cells treated with respective selenosugars at indicated concentrations for 24 h. **b,** ROS assay of T24 and SV-HUV-1 cells treated with respective selenosugars at indicated concentrations for 24 h. **c,** ROS assay of MCF-7 cells treated with ManNSe at indicated concentrations for varies time. All data are from at least three independent experiments. Error bars in all statistical data represent mean ± s.d. (Related Fig. 6)

