## Supplementary information for "SeMOE allows for quantitative glycan perception and exhibits anti-cancer potentiality"

This Supplementary Information includes **Supplementary Figs. 1-13**, **Supplementary Notes 1-2**, **Supplementary Tables 1-7**:

**Supplementary Fig. 1:** Cytotoxicity of selenosugars
**Supplementary Fig. 2:** Calibration curve of Se standard solution
**Supplementary Fig. 3:** Concentration-dependent analysis of Se levels in HeLa cells
**Supplementary Fig. 4:** CyTOF analysis of selenosugar-treated 293T cells
**Supplementary Fig. 5:** Correlation between the cell number and Se level
**Supplementary Fig. 6:** S-glyco-modification of selenosugars
**Supplementary Fig. 7:** Glycopeptide comparison between pGlyco3 and SESTAR++
**Supplementary Fig. 8:** Applicability of SeMOE in Jurkat and K562 cells
**Supplementary Fig. 9:** Systematic evaluation of SeMOE in various cancer cell lines
**Supplementary Fig. 10:** Sialic acid transfer assay between 4T1 and RAW 264.7 cells
**Supplementary Fig. 11:** Imaging of sialoglycoconjugates after GC-oocyte interaction
**Supplementary Fig. 12:** ROS assay of various cell lines after SeMOE treatment
**Supplementary Fig. 13:** Apoptosis assay of MCF-7 cells after SeMOE treatment

**Supplementary Note 1:** Synthetic procedures
**Supplementary Note 2:** Flow cytometric gating strategy

**Supplementary Table 1:** Instrumentation and measurement parameters of ICP-MS in solution
**Supplementary Table 2:** Instrumental parameters of LA-ICP-MS for selenoprotein analysis on PVDF membrane
**Supplementary Table 3:** Instrumental parameters of LA-ICP-TOF MS for mouse tissue sialoglycan imaging
**Supplementary Table 4:** Instrumental parameters of LA-ICP-MS for mouse tissue sialoglycan quantification *in situ*
**Supplementary Table 5:** Peptide spectrum matches in the pGlyco3 searches
**Supplementary Table 6:** MS1 matches in the SESTAR++ searches
**Supplementary Table 7:** Intact *N*-glycopeptides identified by SeMOE and pGlyco3

#### Supplementary Figures

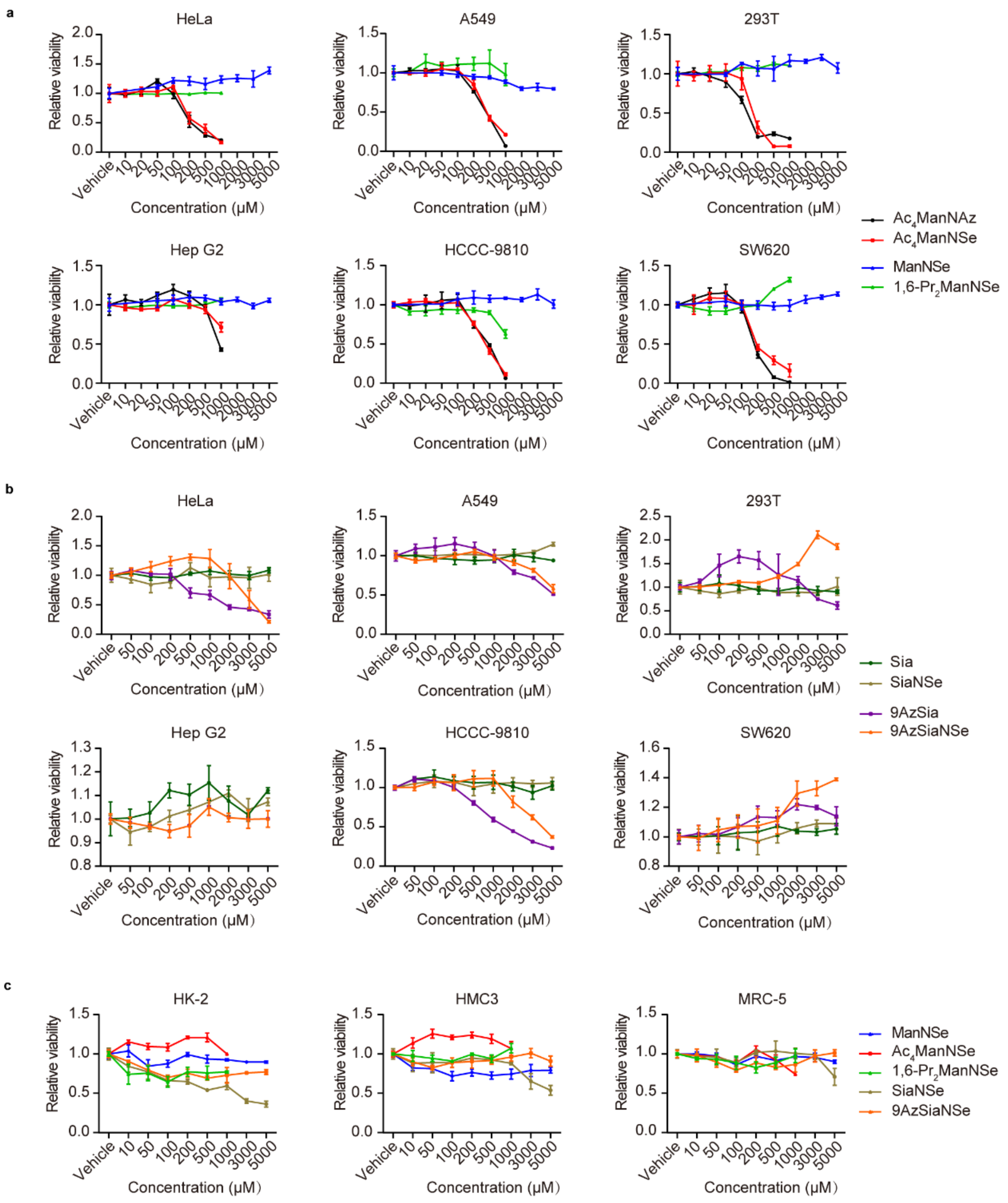


**Supplementary Fig. 1** Cytotoxicity of selenosugars. A variety of cells were incubated with various selenosugars, in a concentration ranging from 0 -5 mM (as designated in each chart), for 48 h. The cells were then analyzed using commercialized CCK-8 cell viability assay. Results are from at least three independent experiments. Error bars represent mean ± s.d. (Related to Fig. 2)

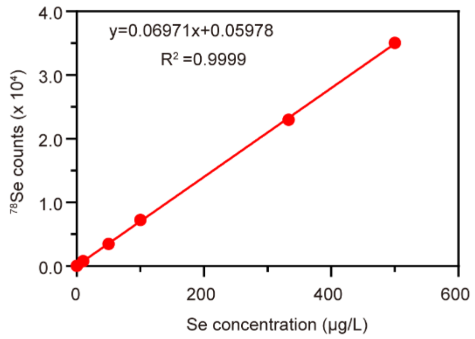


**Supplementary Fig. 2** Calibration curve of Se standard solution. The calibration curve was constructed by Se standard solution of 0, 1, 10, 50, 100, 333 and 500 ppb (μg/L), respectively, against $^{78}$Se-signal intensity, on the solution nebulization ICP-MS (PerkinElmer, NexION 300D, USA). (Related to Fig. 2)

**a**

| 1,6-$Pr_2$ManNSe | | |
|---|---|---|
| Concentration (μM) | Se level (nmol/g protein) | Se relative level |
| 0 | 24.19 ± 1.99 | 1.00 |
| 0.001 | 149.85 ± 4.88 | 6.20 |
| 0.01 | 129.08 ± 13.46 | 5.34 |
| 0.1 | 175.43 ± 17.35 | 7.25 |
| 1 | 241.70 ± 15.53 | 9.99 |
| 10 | 271.20 ± 21.92 | 11.21 |
| 50 | 572.41 ± 29.12 | 23.67 |
| 100 | 873.88 ± 20.96 | 36.13 |
| 200 | 1589.68 ± 26.96 | 65.73 |
| 500 | 1843.87 ± 32.57 | 76.24 |

**b**

| SiaNSe | | |
|---|---|---|
| Concentration (μM) | Se level (nmol/g protein) | Se relative level |
| 0 | 28.20 ± 2.73 | 1.00 |
| 0.01 | 136.59 ± 13.62 | 4.84 |
| 0. 1 | 180.15 ± 22.61 | 6.39 |
| 1 | 159.51 ± 7.03 | 5.66 |
| 10 | 250.43 ± 7.26 | 8.88 |
| 50 | 208.98 ± 11.66 | 7.41 |
| 100 | 219.62 ± 12.68 | 7.79 |
| 200 | 306.57 ± 16.15 | 10.87 |
| 500 | 401.08 ± 21.33 | 14.22 |
| 1000 | 715.21 ± 11.44 | 25.37 |
| 2000 | 1182.05 ± 25.92 | 41.92 |
| 3000 | 1732.64 ± 19.49 | 61.45 |

**Supplementary Fig. 3** Concentration-dependent analysis of Se levels in HeLa cells. HeLa cells were co-incubated with 1,6-$Pr_2$ManNSe **(a)** or SiaNSe **(b)** at varied concentrations (as designated in each table) for 48 h. The cells were lysed for whole protein extraction, and analyzed using ICP-MS. The Se level was calculated based on Se standard curve in Supplementary Fig.2. Results are from at least three independent experiments. Error bars represent mean ± s.d. (Related to Fig. 2)

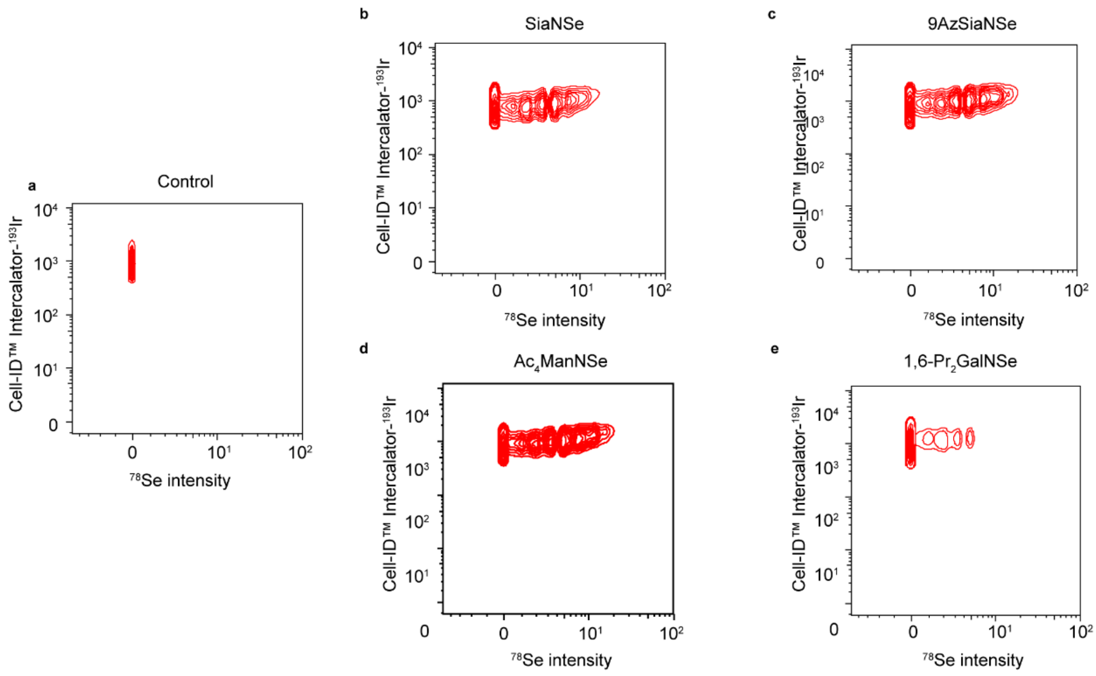


**Supplementary Fig. 4** CyTOF analysis of 293T cells treated with various selenosugars. The 293T cells were treated with vehicle (PBS) **(a)**, 2 mM SiaNSe (**b**), 2 mM 9AzSiaNSe (**c**), 200 µM $Ac_4ManNSe$ (**d**), or 200 µM 1,6-$Pr_2ManNSe$ (**e**) for 48 h, respectively. The cells were washed, stained by Cell-IC$^{TM}$ Intercalator-$^{193}$Ir, and subjected to CyTOF analysis. *m/z* at 78 was used for calculation of $^{78}$Se. (Related to Fig. 2)

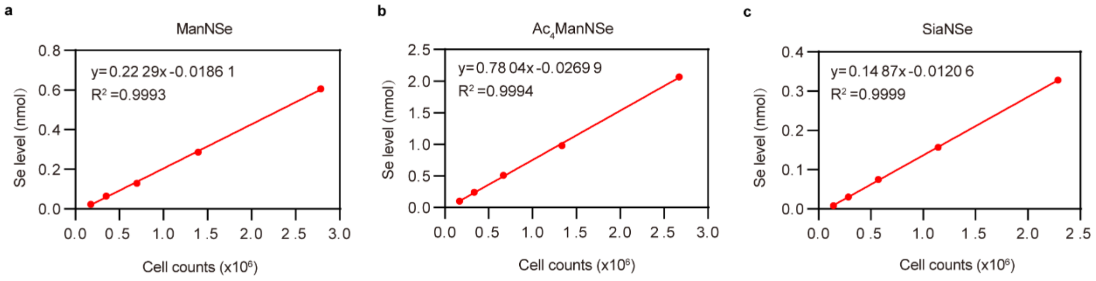


**Supplementary Fig. 5** Linear correlation between the cell number and Se level. HeLa cells were treated with 2 mM ManNSe **(a)**, 200 µM Ac4ManNSe **(b)** or 2 mM SiaNSe **(c)** for 48 h, respectively. The cells were trypsinized, separated based on cell counting numbers ranging from 0 - 2.7 × $10^6$ as various groups. Each group was then subjected to ICP-MS analysis. The linear regression for SeMOE probes is shown in each chart. (Related to Fig. 2)

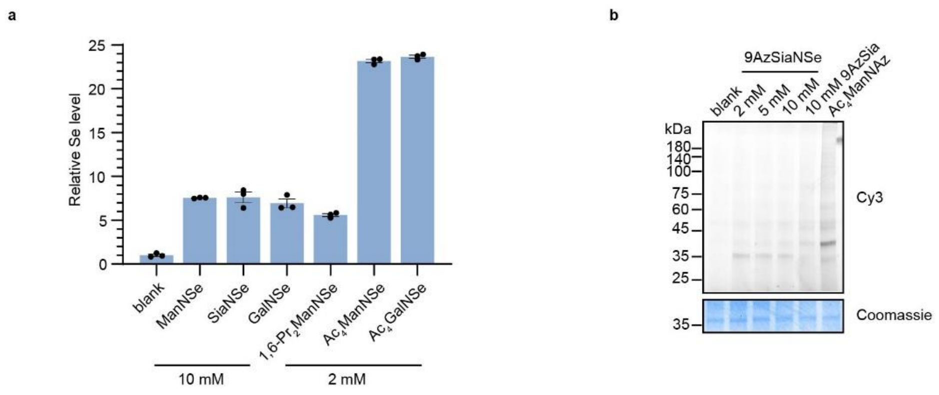


**Supplementary Fig. 6** S-glyco-modification of selenosugars. **a,** ICP-MS analysis of HeLa cell lysates treated with respective unnatural monosaccharides at varied concentrations for 2 h. Results are from at least three independent experiments. Error bars represent mean ± s.d. **b,** In-gel fluorescence scanning showing HeLa cell lysates treated with 9AzSiaNSe at varied concentrations for 2 h, followed by reaction with alkyne-Cy3. (Related to Fig. 2)

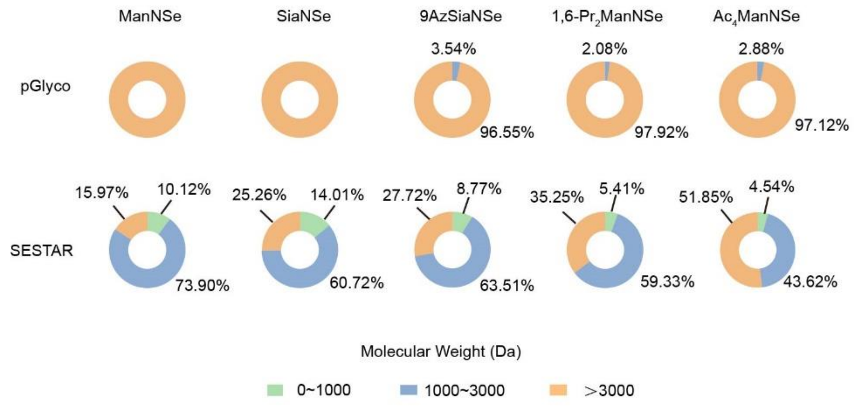


**Supplementary Fig. 7** Comparison of the molecular weight of glycopeptides identified in pGlyco3 and SESTAR++ searches. (Related to Fig. 3)

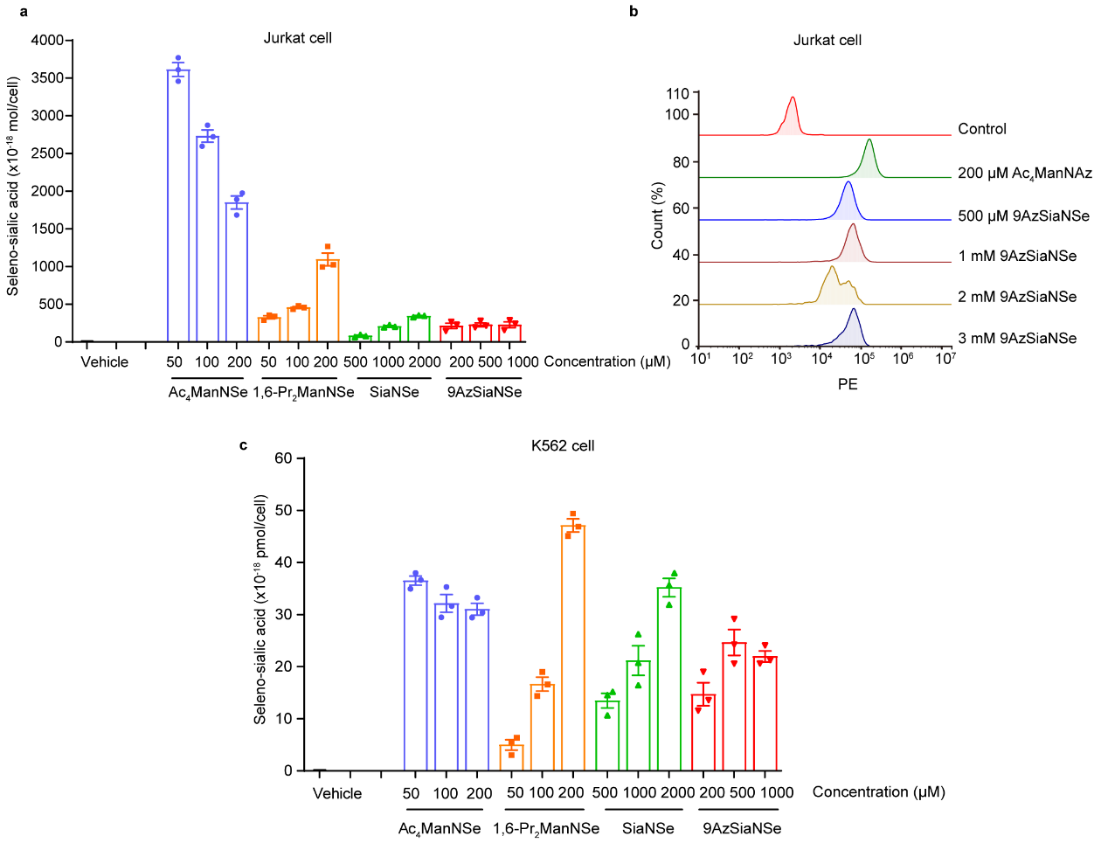


**Supplementary Fig. 8** Applicability of SeMOE in Jurkat and K562 cells. **a,** Seleno-sialic acid levels of Jurkat cells treated with respective selenosugars at indicated concentrations for 48 h were measured by ICP-MS. **b,** Flow cytometry analysis of 9AzSiaNSe-labeled Jurkat cells, followed by reaction with alkyne-Alexa 488. **c,** Seleno-sialic acid levels of K562 cells treated with respective selenosugars at indicated concentrations for 48 h. Results are from at least three independent experiments. Error bars represent mean ± s.d. (Related to Fig. 4)

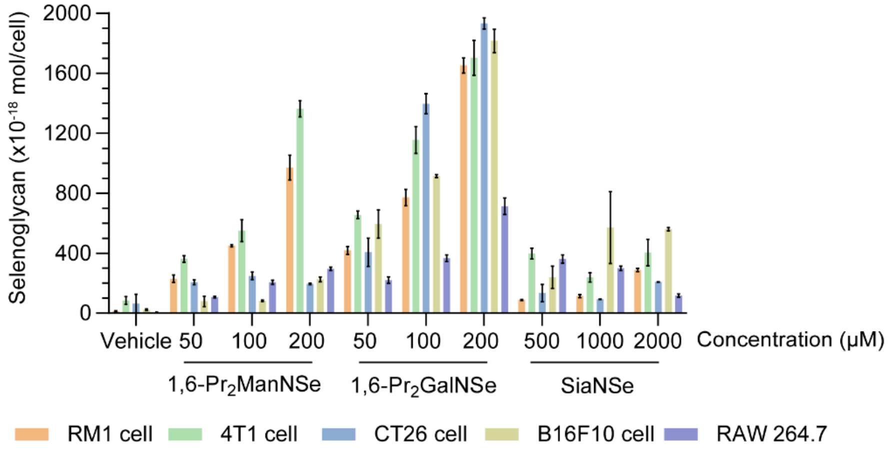


**Supplementary Fig. 9** Systematic evaluation of SeMOE in various cancer cell lines. The cells were treated with respective selenosugars at indicated concentrations for 48 h, and analyzed using ICP-MS. Results are from at least three independent experiments. Error bars represent mean ± s.d. (Related to Fig. 4)

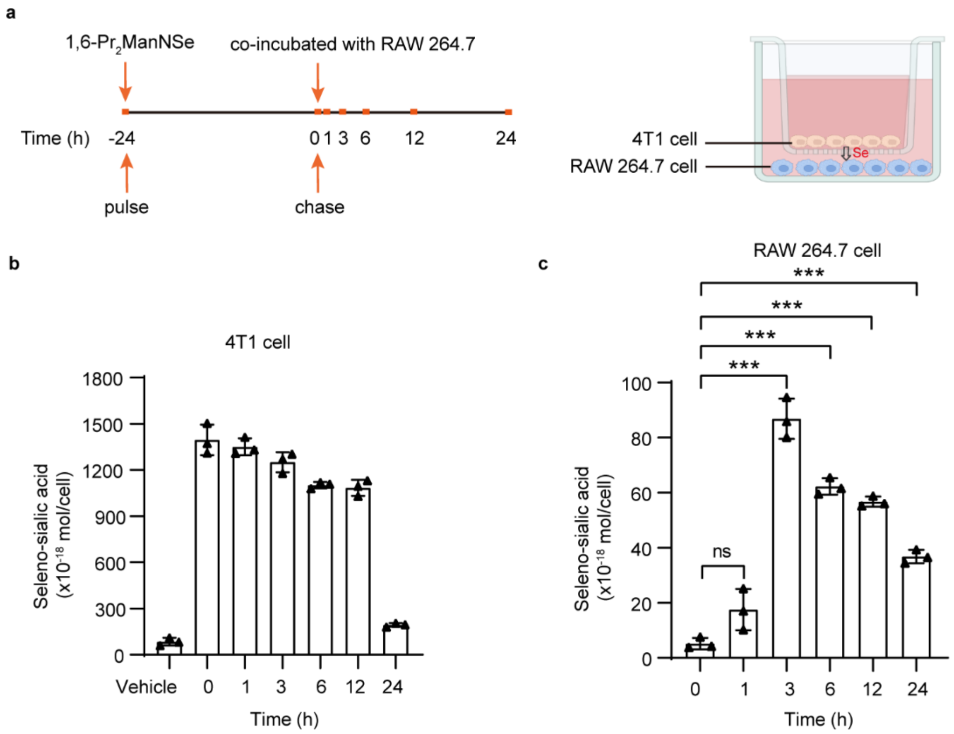


**Supplementary Fig. 10** Sialic acid transfer assay between 4T1 and RAW 264.7 cells. **a,** Schematic workflow of the pulse-chase experiment. 4T1 cells were treated with 200 μM 1,6-$Pr_2$ManNSe for 24 h, washed with PBS, and then co-incubated with RAW 264.7 cells in a 0.4 μm-sized transwell culture system for varied time. **b,** Seleno-sialic acid levels of 4T1 cells during incubation with RAW 264.7 cells. **c,** Seleno-sialic acid transfer from 4T1 cells to RAW 264.7 cells. 0 h represents the initiation of chase experiment. Results are from at least three independent experiments. Error bars represent mean ± s.d. *$P < 0.05$, **$P < 0.01$, ***$P < 0.001$, ns, not significant (two-way ANOVA). (Related to Fig. 4)

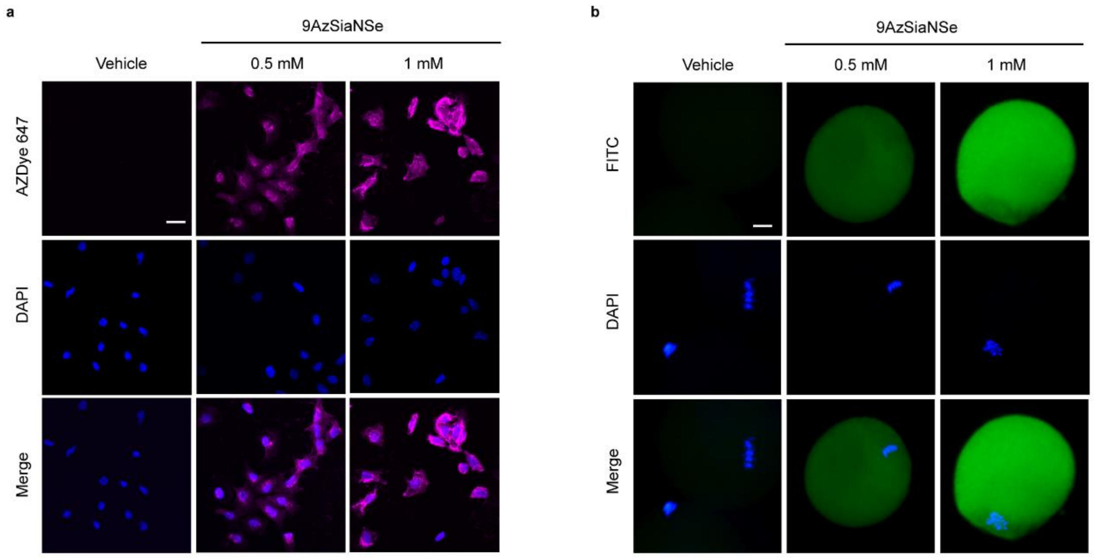


**Supplementary Fig. 11** Confocal fluorescence imaging of 9AzSiaNSe-labeled GCs **(a)** and the oocytes after co-incubation with 9AzSiaNSe-labeled GCs **(b)**. **a,** Mouse GCs were treated with vehicle (PBS) or 9AzSiaNSe at indicated concentrations for 48 h, respectively. The GCs were reacted with alkyne-AZDye 647 via click chemistry. **b,** Mouse primary oocytes were co-incubated with 9AzSiaNSe-labeled or PBS-treated GCs for 14-16 h, respectively. The oocytes were reacted with alkyne-AZDye 488 via click chemistry. The nucleus was stained by DAPI. Scale bar: 20 μm. (Related to Fig. 4)

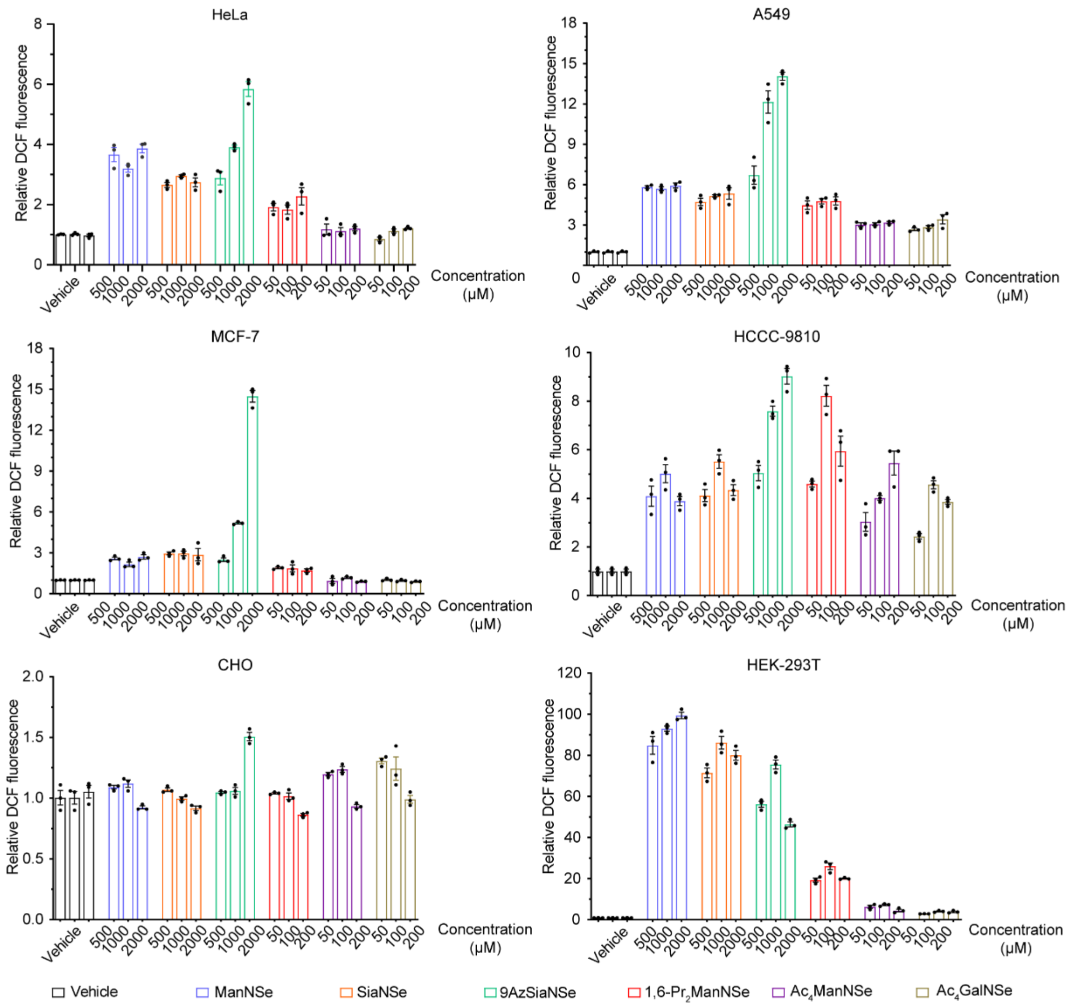


**Supplementary Fig. 12** ROS assay of various cell lines treated with respective selenosugars at indicated concentrations for 24 h. The ROS level was assayed using flow cytometry. Results are from at least three independent experiments. Error bars represent mean ± s.d. (Related to Fig. 6)

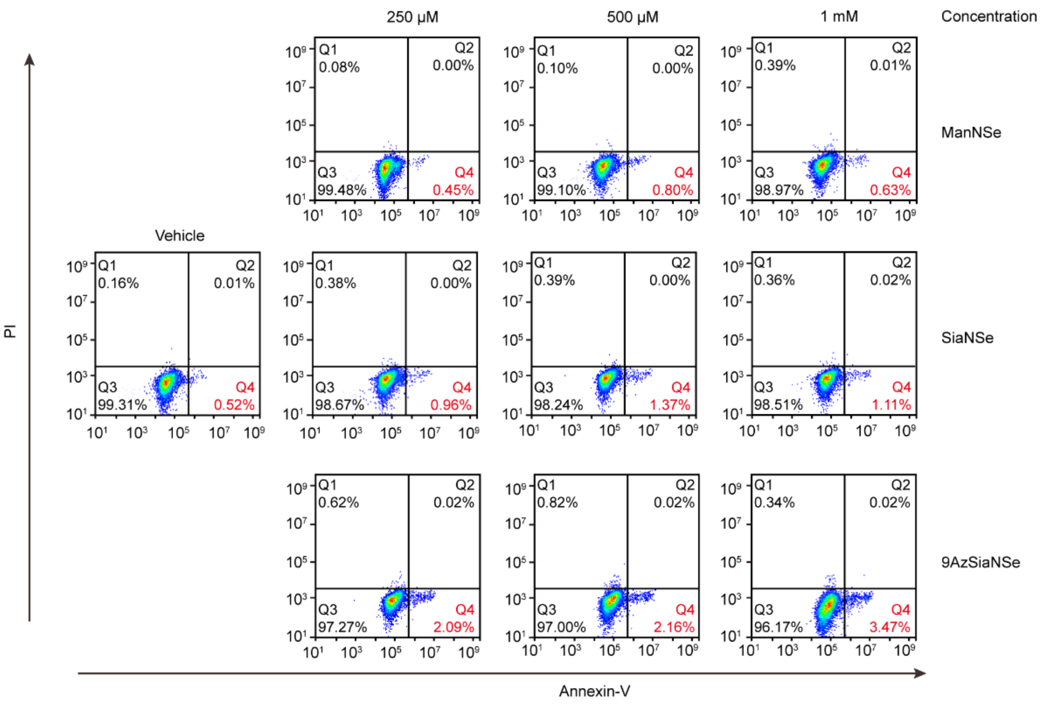


**Supplementary Fig. 13** Apoptosis assay of MCF-7 cells. MCF-7 cells were treated with ManNSe, SiaNSe, or 9AzSiaNSe at indicated concentrations for 48 h. The viability of each cell group was measured using a commercialized Annexin-V/PI assay. (Related to Fig. 6)

#### Supplementary Note 1: Synthetic procedures

#### General synthetic chemistry instrumentation

SiliCycle silica plates (TLG R10011B-624) were used for thin layer chromatography (TLC) with detection of multiband UV-absorption (254 to 365 nm). Column chromatography was carried out using SepaBean™ machine T flash chromatography system with an automated fraction collector. Semi-prep HPLC was carried out using a Waters LC Prep 150 system equipped with a 2998 photodiode array detector, an automatic fraction collector and a XBridge Prep C18 column (19 mm×150 mm, 5 µm) or a XBridge Prep amide column (19 mm×250 mm, 5 µm). Proton nuclear magnetic resonanc ($^1$H NMR) and proton-decoupled carbon-13 nuclear magnetic resonance (13C {1H} NMR) spectra were obtained on a 400 MHz Bruker AVANCE III-400 spectrometer at 25 °C. All chemical NMR analysis was conducted on MestReNova v12.0.3. Data are presented in the form of chemical shift, multiplicity, coupling constants in Hertz (Hz), and integration. Chemical shifts are reported in δ (ppm) relative to the solvent residual peak. Coupling constants are reported in Hz with multiplicities denoted as s (singlet), d (doublet), t (triplet), q (quartet), m (multiplet) and dd (doublet of doublet). High resolution mass spectrometry (HR-MS) was conducted on a Thermo Fisher Q Exactive LC/MS.

#### Synthesis of 2-(methylselanyl)acetic acid NHS ester

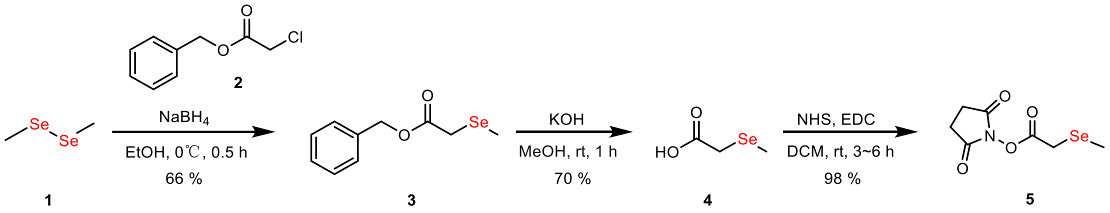


**Benzyl 2-(methylselanyl)acetate (3):** $NaBH_4$ (2.31 g, 61.18 mmol, 2.3 eq.) was added slowly to a stirred solution of **1** (5.0 g, 26.60 mmol, 1.0 eq.) in anhydrous EtOH (150 mL) at 0°C under a nitrogen atmosphere. The reaction mixture was stirred for ~10 min at 0°C until the characteristic diselenide yellow color disappeared. Subsequently, **2** (10.8 g, 58.52 mmol, 2.2 eq.) was added dropwise in 5 min, with the observation of immediate precipitation of a white solid, and the reaction mixture was then stirred for 0.5 h, and monitored using TLC. Deionized water (10 mL) and NaCl (5 g) was added to quench the reaction, the product was extracted with diethyl ether (4 x 150 mL), dried over $Na_2SO_4$, filtered, and concentrated to a yellow oil. Purification via silica gel flash column chromatography eluted with PE gave **3** as a yellow oil (8.54 g, 35.12 mmol, 66%). **TLC** (Rf = 0.3, PE: EA=30:1, stained by phosphomolybdic acid). **$^1$H NMR** (400 MHz, $CDCl_3$) *δ* 7.40 – 7.31 (m, 5H), 5.16 (s, 2H), 3.18 (s, 2H), 2.14 (s, 3H). **$^{13}$C NMR** (101 MHz, $CDCl_3$) δ 171.32, 135.83, 128.67, 128.41, 128.31, 66.98, 23.71, 6.01. **ESI-HRMS:** Calcd for $C_{10}H_{12}O_2SeNa$ $[M+Na]^+$ = 266.9895, found 266.9870

**2-(methylselanyl)acetic acid (4):** To a solution of **3** (8.54 g, 35.12 mmol, 1.0 eq.) in $MeOH/H_2O$ (3/1; 80 mL) at 0°C, KOH (2.76 g, 49.17 mmol, 1.4 eq.) was added slowly. The reaction was warmed to room temperature and stirred for 1 h, diluted with deionized water (50 mL), and EtOAc (4 x 70 mL). The aqueous layer was collected, followed by addition of HCl (1 M aqueous solution) to pH 4. Then, the aqueous layer was extracted with $CH_2Cl_2$ (4 x 200 mL). The pooled organic extracts were dried over $Na_2SO_4$, filtered, and concentrated to obtain a colorless liquid (3.76 g, 24.58 mmol, 70%). **$^1$H NMR** (400 MHz, DMSO-$d_6$) δ 3.12 (s, 1H), 2.08 (s, 1H). **$^{13}$C NMR** (101 MHz, DMSO-$d_6$) δ 172.43, 23.88, 5.21. **ESI-HRMS**: Calcd for $C_3H_5O_2Se^-$ $[M-H]^-$ = 152.9533, found 152.9440

**2,5-dioxopyrrolidin-1-yl 2-(methylselanyl)acetate (5):** To a solution of **4** (3.5 g, 22.87 mmol, 1.0 eq.) in $CH_2Cl_2$ (80 mL), *N*-hydroxysuccinimide (2.76 g, 24.01 mmol, 1.05 eq.) and 1-ethyl(3-dimethylaminopropyl)carbodiimide (EDC) (8.77 g, 45.74 mmol, 2.0 eq.) were added. The reaction was stirred at room temperature for 3-6 h, concentrated *in vacuo* to give **5** as a white solid (5.61 g, 22.41 mmol, 98 %). The product was used immediately for the next step without any purification. **TLC** (Rf = 0.5, PE: EA=1:1, stained by iodine).

#### Synthesis of ManNSe, GalNSe

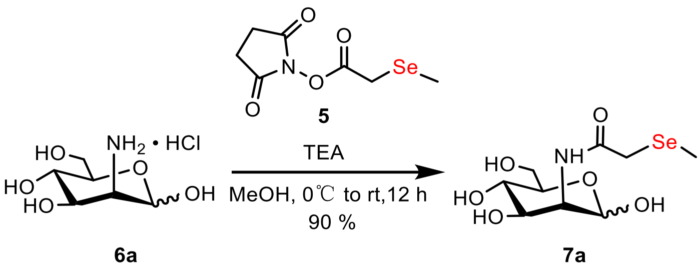


**2-(methylselanyl)-N-((*3S,4R,5S,6R*)-2,4,5-trihydroxy-6-(hydroxymethyl)tetrahydro-*2H*-pyran-3-yl)acetamide (7a, ManNSe):** Under a nitrogen atmosphere, a solution of **6a** (3.0 g, 13.91 mmol, 1.0 eq.) and triethylamine (TEA) (6 mL, 41.73 mmol, 3.0 eq.) in anhydrous MeOH (100 mL) was cooled to 0°C, and then **5** (3.47 g, 13.91 mmol, 1.0 eq.) was added. The reaction was warmed to room temperature while stirred overnight, and concentrated *in vacuo* as a yellow oil. A first purification via silica gel flash column chromatography (5-20 % MeOH in DCM) gave **7a** as a light-yellow solid (3.93 g, 12.52 mmol, 90%). The crude product was further purified by normal phase HPLC (XBridge Prep amide column,19 mm×250 mm, 5 μm) to give the product as white solid, eluted with 0-95% water in ACN at the flow rate of 7 mL/min. **TLC** (Rf = 0.4, DCM:MeOH=3:1, stained by 10% $H_2SO_4$ in MeOH). **$^1$H NMR:** α: β=1:0.78 (400 MHz, $D_2O$) δ 5.14 (d, J = 1.4 Hz, 1H), 5.04 (d, *J* = 1.6 Hz, 0.78H), 4.44 (dd, *J* = 4.4, 1.4 Hz, 0.78H), 4.31 (dd, *J* = 4.6, 1.5 Hz, 1H), 4.07 (dd, *J* = 9.8, 4.7 Hz, 1H), 3.94 – 3.74 (m, 6H), 3.62 (t, *J* = 9.7 Hz, 1H), 3.52 (t, *J* = 9.8 Hz, 1H), 3.42 (ddd, *J* = 9.9, 5.0, 2.3 Hz, 1H), 2.84 (s, 4H), 2.13 (d, *J* = 9.7 Hz, 5H). **$^{13}$C NMR:** (anomers, 101 MHz, $D_2O$) δ 175.35, 174.63, 92.98, 92.88, 76.37, 72.01, 71.94, 68.69, 66.76, 66.53, 60.42, 54.46, 53.60, 25.79, 25.48, 4.82, 4.76. **ESI-HRMS**: Calcd for $C_9H_{17}NO_6SeNa$ $[M+Na]^+$= 338.0114, found 338.0101

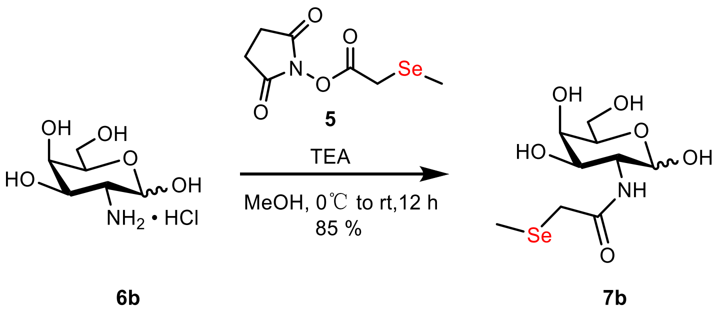


**2-(methylselanyl)-N-((*3R,4R,5R,6R*)-2,4,5-trihydroxy-6-(hydroxymethyl)tetrahydro-*2H*-pyran-3-yl)acetamide (7b, GalNSe):** **7b** was obtained with the same procedure for synthesis of **7a** as a white solid. **TLC** (Rf = 0.4, DCM: MeOH=3:1, stained by 10% $H_2SO_4$ in MeOH). **$^1$H NMR:** α: β=1:0.68 (400 MHz, $D_2O$) δ 5.25 (d, *J* = 3.7 Hz, 1H), 4.67 (d, *J* = 8.4 Hz, 0.68H), 4.18 – 4.08 (m, 2H), 3.99 (d, *J* = 3.0

Hz, 1H), 3.92 (dd, $J$ = 11.2, 3.0 Hz, 2H), 3.89 – 3.84 (m, 0.68H), 3.80 – 3.66 (m, 5H), 3.32 – 3.15 (m, 6H), 2.20 – 2.03 (m, 5H). **$^{13}$C NMR**: (anomers, 101 MHz, $D_2O$) δ174.71, 174.53, 95.31, 90.92, 75.13, 70.96, 70.51, 68.64, 68.42, 67.92, 67.26, 61.23, 60.98, 54.00, 50.62, 25.92, 25.59, 4.52, 4.44. **ESI-HRMS**: Calcd for $C_9H_{17}NO_6SeNa$ $[M+Na]^+$ = 338.0114, found 338.0102

#### Synthesis of Ac4ManNSe, Ac4GalNSe

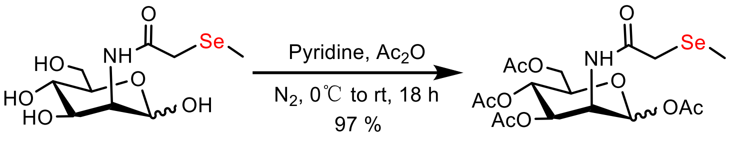


**7a** **8a**

**(*3S,4R,5S,6R*)-6-(acetoxymethyl)-3-(2-(methylselanyl)acetamido)tetrahydro-*2H*-pyran-2,4,5-triyl triacetate (8a, Ac4ManNSe): 7a** (1.0 g, 3.18 mmol) was dissolved in pyridine (50 mL), cooled to 0°C, and acetic anhydride (25 mL) was added. The reaction was warmed to room temperature and stirred for 18 h. The reaction mixture was diluted with ethyl acetate, and sequentially washed with 1 M HCl, sat. aq. $NaHCO_3$, and brine. The organic phase was dried over $Na_2SO_4$, filtered, and concentrated in vacuum. The residue was first purified by silica gel flash column chromatography (5-50% EA in PE). Further purification via reversed phase HPLC (0-90% ACN in water at the flow rate of 7 mL/min) gave **8a** as a white solid (1.49 g, 3.09 mmol, 97%). **TLC** (Rf = 0.5, PE: EA =1:1, stained by 10% $H_2SO_4$ in MeOH). **$^1$H NMR**: α: β=1:0.84 (400 MHz, $CDCl_3$) δ 6.04 (d, $J$ = 1.7 Hz, 1H), 5.90 (d, $J$ = 1.6 Hz, 0.84H), 5.35 (dd, $J$ = 10.3, 4.1 Hz, 1H), 5.31 – 5.18 (m, 2H), 5.06 (dd, $J$ = 10.0, 3.8 Hz, 0.84H), 4.75 (ddd, $J$ = 9.3, 3.8, 1.5 Hz, 0.84H), 4.63 (ddd, $J$ = 9.7, 4.0, 1.9 Hz, 1H), 4.28 (dd, $J$ = 12.4, 3.9 Hz, 2H), 4.17 – 4.03 (m, 4H), 3.83 (ddd, $J$ = 9.8, 4.3, 2.4 Hz, 0.84H), 3.28 (d, $J$ = 14.0 Hz, 4H), 2.22 – 1.99 (m, 32H). **$^{13}$C NMR**: (anomers, 101 MHz, $CDCl_3$) δ 170.68, 170.14, 169.63, 168.27, 91.71, 90.64, 73.46, 71.79, 70.38, 69.35, 65.10, 65.00, 61.75, 61.70, 50.13, 49.74, 27.79, 27.68, 20.92, 20.89, 5.53, 5.35. **ESI-HRMS**: Calcd for $C_{17}H_{25}NO_{10}SeNa$ $[M+Na]^+$ = 506.0536, found 506.0527

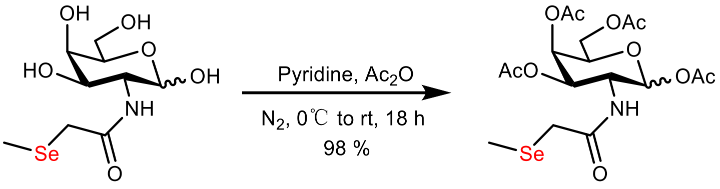


**7b** **8b**

**(*3R,4R,5R,6R*)-6-(acetoxymethyl)-3-(2-(methylselanyl)acetamido)tetrahydro-*2H*-pyran-2,4,5-triyl triacetate (8b, Ac4GalNSe)**: **8b** was obtained from **7b** with the same procedure for synthesis of **8a** as a white solid. **TLC** (Rf = 0.5, PE:EA =1:1, stained by 10% $H_2SO_4$ in MeOH). **$^1$H NMR**: α: β=1:0.43 (400 MHz, $CDCl_3$) δ 6.46 – 6.29 (m, 1.44H), 6.24 (d, $J$ = 3.5 Hz, 1H), 5.79 (d, $J$ = 8.8 Hz, 0.43H), 5.42 (dd, $J$ = 21.3, 3.2 Hz, 1.49H), 5.29 – 5.16 (m, 1.55H), 4.74 – 4.65 (m, 1H), 4.43 – 4.32 (m, 0.68H), 4.25 (dt, $J$ = 10.9, 5.5 Hz, 1.44H), 4.17 – 4.03 (m, 3H), 3.15 (dd, $J$ = 23.5, 11.9 Hz, 3H), 2.27 – 1.98 (m, 25H). **$^{13}$C NMR**: (anomers, 101 MHz, $CDCl_3$) δ 171.05, 170.50, 170.35, 169.78, 168.95, 92.82, 91.16, 70.25, 68.75, 67.89, 66.78, 66.50, 61.40, 47.27, 27.16, 21.04, 20.82, 5.71, 5.62. **ESI-HRMS**: Calcd for $C_{17}H_{25}NO_{10}SeNa$ $[M+Na]^+$= 506.0536, found 506.0527

#### Synthesis of 1,6-Pr2ManNSe, 1,6-Pr2GalNSe

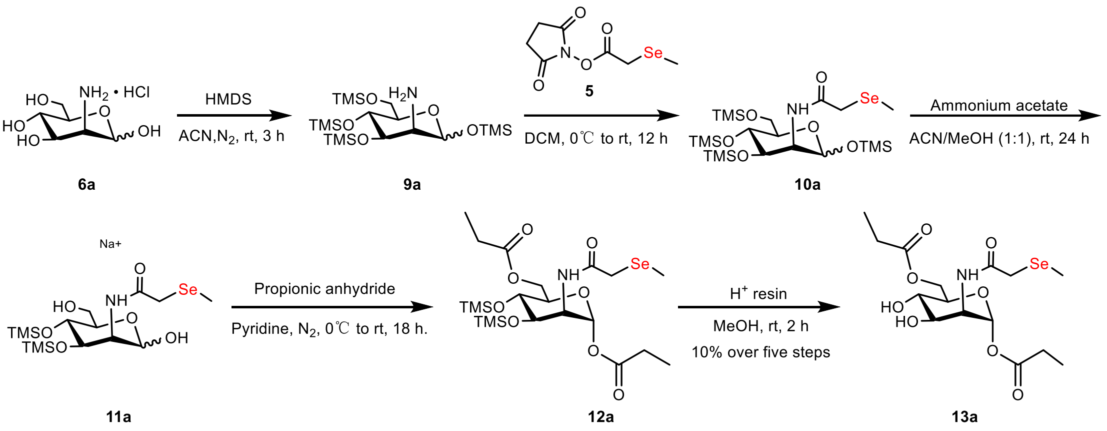

**(*3S,4R,5R,6R*)-2,4,5-tris((trimethylsilyl)oxy)-6-(((trimethylsilyl)oxy)methyl)tetrahydro-*2H*-pyran-3-amine (9a)**: Under a nitrogen atmosphere, to a suspension of **1a** (*D*-Mannosamine·HCl, 5.0 g, 23.19 mmol, 1.0 eq.) in anhydrous ACN (100 mL), hexamethyldisilazane (HMDS) (12.3 mL, 57.97 mmol, 2.5 eq.) was added dropwise. The reaction was stirred at room temperature for 3 h, filtered to remove white precipitates, and the residue was concentrated to a colorless oil. The product was used immediately for the next step without further purification. **TLC** (Rf = 0.6, PE:EA =3:1, stained by 10% $H_2SO_4$ in MeOH)

**2-(methylselanyl)-N-((*3S,4R,5R,6R*)-2,4,5-tris((trimethylsilyl)oxy)-6 (((trimethylsilyl)oxy)methyl)tetrahydro-*2H*-pyran-3-yl)acetamide (10a)**: A solution of **5** (5.8 g, 23.19 mmol, 1.0 eq) was cooled to 0°C and added to freshly-prepared **9a**. The reaction was warmed to room temperature and stirred overnight. The mixture was diluted with DCM, and washed with sat. aq. $NaHCO_3$. The organic phase was collected, dried over $Na_2SO_4$ and concentrated to a yellow oil. The product was used immediately for the next step without further purification. **TLC** (Rf = 0.5, PE:EA =8:1, stained by 10% $H_2SO_4$ in MeOH)

**N-((*3S,4R,5R,6R*)-2-hydroxy-6-(hydroxymethyl)-4,5-bis((trimethylsilyl)oxy)tetrahydro-*2H*-pyran-3-yl)-2-(methylselanyl)acetamide (11a): 10a** obtained in the previous step was re-dissolved in the mixed solvent of ACN/MeOH (100 mL, 1:1 v/v), ammonia acetate (3.58 g, 46.38 mmol, 2.0 eq) was added. The reaction was stirred at room temperature for 24 h, concentrated *in vacuo*, diluted with EtOAc and washed four times with brine. The organic phase was collected, dried over $Na_2SO_4$, and concentrated to **11a** as a light-yellow oil. The product was used immediately for the next step without further purification. **TLC** (Rf = 0.6, DCM:MeOH =9:1, stained by 10% $H_2SO_4$ in MeOH). **ESI-HRMS**: Calcd for $C_{15}H_{33}NO_6SeSi_2Na$ $[M+Na]^+$= 482.0904, found 482.0897

**(*2R,3S,4R,5R,6R*)-3-(2-(methylselanyl)acetamido)-6-((propionyloxy)methyl)-4,5-bis((trimethylsilyl)oxy)tetrahydro-*2H*-pyran-2-yl propionate (12a): 11a** obtained in the previous step was re-dissolved in pyridine (80 mL), cooled to 0°C, and then propionic anhydride (45 mL) was added. The reaction was warmed to room temperature and stirred for 18 h. The reaction mixture was diluted sequentially with ethyl acetate, washed with 1 M HCl, sat. aq. $NaHCO_3$, and brine. The organic phase was dried over $Na_2SO_4$, filtered, and concentrated to a yellow oil. The product was used immediately for the next step without further purification. **TLC** (Rf = 0.6, PE:EA =3:1, stained by 10% $H_2SO_4$ in MeOH). **ESI-HRMS**: Calcd for $C_{12}H_{41}NO_8SeSi_2Na$ $[M+Na]^+$= 574.1429, found 594.1436

**(*2R,3S,4R,5S,6R*)-4,5-dihydroxy-3-(2-(methylselanyl)acetamido)-6-((propionyloxy)methyl)tetrahydro-*2H*-pyran-2-yl propionate (13a, 1,6-Pr$_2$ManNSe): 12a** obtained in the previous step was re-dissolved in MeOH (100 mL), and then Dowex H$^+$ resin was added. The reaction was stirred at room temperature for 2 h. The mixture was filtrated and concentrated in vacuum. The residue was first purified by silica gel flash column chromatography (0-2% MeOH in DCM). Then further purification via reversed phase HPLC (10-100% ACN in water at the flow rate of 7 mL/min) gave **13a** as a white solid (1.03 g, 2.42 mmol). Total yield:10% over five steps. **TLC** (Rf = 0.5, DCM:MeOH =10:1, stained by 10% $H_2SO_4$ in MeOH). **$^1$H NMR**: (500 MHz, MeOD) δ 6.02 (d, *J* = 1.7 Hz, 1H), 4.41 (dd, *J* = 11.9, 2.2 Hz, 1H), 4.33 – 4.28 (m, 2H), 4.03 (dd, *J* = 9.4, 4.9 Hz, 1H), 3.88 – 3.83 (m, 1H), 3.67 (t, *J* = 9.7 Hz, 1H), 3.36 (dt, *J* = 3.3, 1.6 Hz, 2H), 3.28 (q, *J* = 12.5 Hz, 3H), 2.48 (q, *J* = 7.5 Hz, 3H), 2.41 (dd, *J* = 15.1, 7.6 Hz, 3H), 2.19 (s, 4H), 1.21 (d, *J* = 7.5 Hz, 3H), 1.19 (d, *J* = 4.8 Hz, 3H), 1.16 (d, *J* = 7.6 Hz, 3H). **$^{13}$C NMR**: (126 MHz, MeOD) δ 176.05, 174.38, 173.69, 93.23, 73.89, 70.13, 68.54, 64.70, 53.83, 28.21, 26.38, 9.41, 9.26, 5.13. **ESI-HRMS**: Calcd for $C_{15}H_{25}NO_8SeNa$ $[M+Na]^+$= 450.0638, found 450.0620

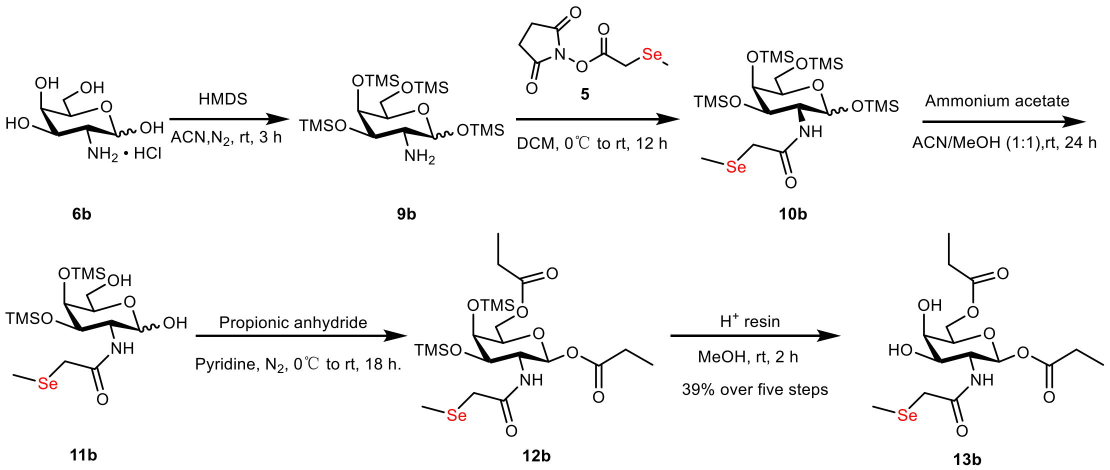


**(*2S,3R,4R,5R,6R*)-4,5-dihydroxy-3-(2-(methylselanyl)acetamido)-6-((propionyloxy)methyl)tetrahydro-*2H*-pyran-2-yl propionate (13b, 1,6-Pr$_2$GalNSe): 13b** (3.9 g,9.14 mmol, 1.0 eq.) was synthesized from **6b** following the procedure for the synthesis of **13a**. The total yield of **13b** was 39% over five steps. **TLC** (Rf = 0.5, DCM: MeOH =10:1, stained by 10% $H_2SO_4$ in MeOH). **$^1$H NMR**: (400 MHz, $D_2O$) δ 6.17 (d, *J* = 3.8 Hz, 1H), 4.40 – 4.23 (m, 4H), 4.14 – 4.04 (m, 2H), 3.27 (d, *J* = 12.6 Hz, 1H), 3.21 (t, *J* = 10.6 Hz, 1H), 2.57 – 2.47 (m, 2H), 2.47 – 2.38 (m, 2H), 2.09 (d, *J* = 4.3 Hz, 2H), 1.15 (t, *J* = 7.5 Hz, 3H), 1.10 (t, *J* = 7.6 Hz, 3H). **$^{13}$C NMR**: (101 MHz, $D_2O$) δ 177.25, 176.01, 174.81, 90.62, 70.66, 68.12, 66.90, 63.82, 49.01, 27.25, 25.26, 8.31, 8.23, 4.54. **ESI-HRMS**: Calcd for $C_{15}H_{25}NO_8SeNa$ $[M+Na]^+$= 450.0638, found 450.0620

#### Synthesis of SiaNSe

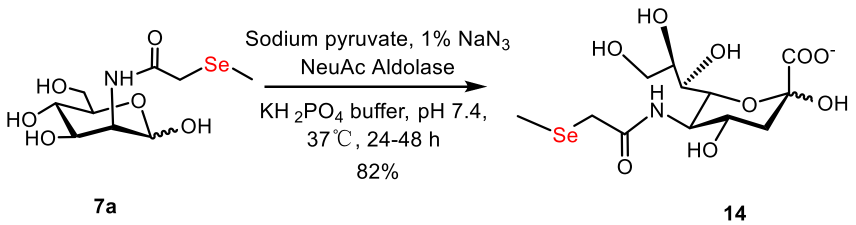


**(*4S,5R*)-2,4-dihydroxy-5-(2-(methylselanyl)acetamido)-6-((*1R,2R*)-1,2,3-**

**trihydroxypropyl)tetrahydro-*2H*-pyran-2-carboxylic acid (14, SiaNSe): 7a** (1.0 g, 3.18 mmol, 1.0 eq.) was dissolved in 40 mL of 0.050 M potassium phosphate buffer (pH 7.4), followed by addition of sodium pyruvate (3.5 g, 31.8 mmol, 10.0 eq.), $NaN_3$ (final concentration of 1% w/v) and NeuAc aldolase (60-80 U). The mixture was reacted at 37°C for 24-48 h, concentrated *in vacuo*, and purified by anion-exchange chromatography with AG1X2 resin, formate form (Bio-Rad). The product was eluted with a gradient of 0.5 M to 1.0 M formic acid at 1.0 mL/min. The fractions containing the desired product were combined and concentrated in vacuum. Further purification via normal phase HPLC (XBridge Prep amide column,19 mm×250 mm, 5 µm) gave **14** as a white solid (1.0 g, 2.61 mmol, 82%), eluted with 0-95% water in ACN at the flow rate of 7 mL/min. **TLC** (Rf = 0.3, DCM:MeOH:$H_2O$ = 3:2:0.5, stained by 10% $H_2SO_4$ in MeOH). **$^1$H NMR:** (400 MHz, $D_2O$) δ 4.12 – 4.03 (m, 2H), 3.97 – 3.89 (m, 1H), 3.83 (dd, *J* = 11.9, 2.6 Hz, 1H), 3.74 (ddd, *J* = 9.0, 6.2, 2.6 Hz, 1H), 3.64 – 3.57 (m, 2H), 3.23 (d, *J* = 1.1 Hz, 2H), 2.31 (dd, *J* = 13.0, 4.9 Hz, 1H), 2.15 – 2.09 (m, 3H), 1.87 (dd, *J* = 13.0, 11.5 Hz, 1H).**$^{13}$C NMR:** (101 MHz, $D_2O$) δ 174.89, 173.37, 95.34, 70.45, 70.21, 68.32, 66.49, 63.13, 52.24, 39.01, 25.78, 4.95. **ESI-HRMS**: Calcd for $C_{12}H_{20}NO_9Se^-$ $[M-H]^-$= 402.0309, found 402.0302

#### Synthesis of 9AzSiaNSe

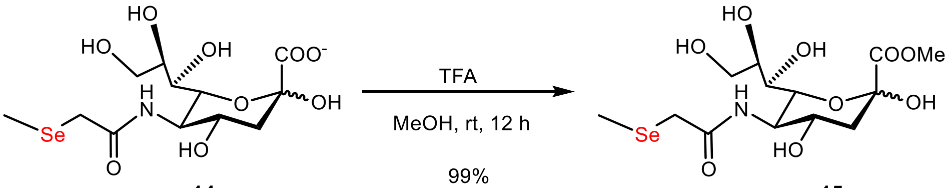


14 15

**methyl (*4S,5R*)-2,4-dihydroxy-5-(2-(methylselanyl)acetamido)-6-((*1R,2R*)-1,2,3-trihydroxypropyl)tetrahydro-*2H*-pyran-2-carboxylate (15):** To a solution of **14** (500 mg, 1.25 mmol,1.0 eq.) in 50 mL MeOH, 0.5 mL trifluoroacetic acid (TFA) was added. The reaction was stirred at room temperature overnight. The solvent was removed in vacuo, and the residue was purified by silica gel flash chromatography eluted with DCM: MeOH gradually from 15:1 to 9:1 to give the desired product as a white solid (515 mg, 1.24 mmol, 99%). **TLC** (Rf = 0.4, DCM: MeOH =4:1, stained by 10% $H_2SO_4$ in MeOH). **$^1$H NMR:** (400 MHz, $D_2O$) δ 4.09 (ddd, *J* = 16.8, 9.1, 2.9 Hz, 2H), 3.94 (t, *J* = 10.3 Hz, 1H), 3.88 – 3.79 (m, 4H), 3.77 – 3.70 (m, 1H), 3.62 (dd, *J* = 11.2, 6.3 Hz, 2H), 3.25 (d, *J* = 0.9 Hz, 2H), 2.33 (dd, *J* = 13.1, 4.9 Hz, 1H), 2.21 – 2.07 (m, 3H), 1.93 (dd, *J* = 13.1, 11.5 Hz, 1H).**$^{13}$C NMR:** (101 MHz, $D_2O$) δ 174.90, 171.40, 95.35, 70.41, 70.17, 68.32, 66.46, 63.15, 53.48, 52.26, 38.85, 25.79, 4.95. **ESI-HRMS:** Calcd for $C_{13}H_{23}NO_9SeNa$ $[M+Na]^+$= 440.0431, found 440.0421

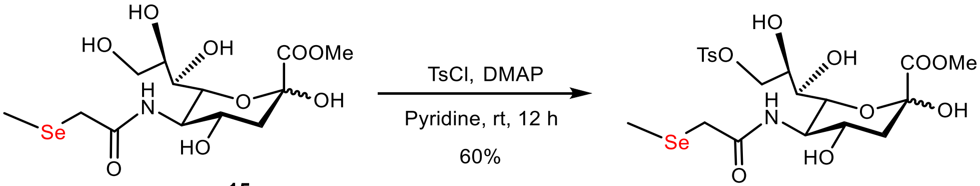


15 16

**Methyl (*4S,5R*)-6-((*1R,2R*)-1,2-dihydroxy-3-(tosyloxy)propyl)-2,4-dihydroxy-5-(2-(methylselanyl)acetamido)tetrahydro-*2H*-pyran-2-carboxylate (16):** A solution of **15** (500 mg, 1.20 mmol, 1.0 eq.) in anhydrous pyridine (50 mL) was cooled to 0°C, then TsCl (274.5 mg, 1.44 mmol, 1.2 eq.) and DMAP (14.7 mg, 0.12 mmol, 0.1 eq.) was added. The reaction mixture was stirred at room temperature overnight under a nitrogen atmosphere. The solvent was removed in vacuum, and the residue was purified by silica gel flash chromatography with DCM:MeOH gradually from 20:1 to 15:1 to give **16** as a white solid (411 mg, 0.71 mmol, 60%). **TLC** (Rf=0.6, DCM: MeOH =9:1, stained by 10% $H_2SO_4$ in MeOH). **$^1$H NMR:** (400 MHz, MeOD) δ 7.82 (d, *J* = 8.3 Hz, 2H), 7.46 (d, *J* = 8.0 Hz, 2H), 4.28 (dd,

$J$ = 9.9, 2.2 Hz, 1H), 4.11 – 4.01 (m, 2H), 3.99 (dd, $J$ = 10.5, 1.4 Hz, 1H), 3.90 – 3.83 (m, 1H), 3.81 – 3.75 (m, 4H), 3.59 (dd, $J$ = 9.2, 1.4 Hz, 1H), 3.33 (dt, $J$ = 3.2, 1.6 Hz, 1H), 3.33 (dt, $J$ = 3.2, 1.6 Hz, 1H), 3.22 – 3.15 (m, 2H), 2.47 (s, 3H), 2.22 (dd, $J$ = 12.9, 4.9 Hz, 1H), 2.19 – 2.14 (m, 3H), 1.89 (dd, $J$ = 12.8, 11.5 Hz, 1H).**$^{13}$C NMR:** (101 MHz, MeOD) δ 175.91, 171.64, 146.39, 134.18, 131.02, 129.17, 96.65, 73.56, 71.95, 69.79, 69.18, 67.57, 54.41, 53.25, 40.85, 26.49, 21.58, 5.63. **ESI-HRMS**: Calcd for $C_{20}H_{29}NO_{11}SSeNa$ [M+Na]$^+$ = 594.0519, found 594.0512

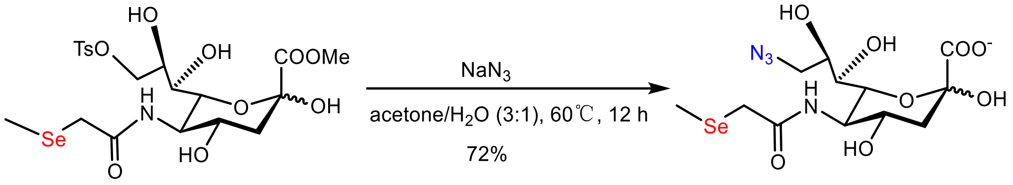


**16** **17**

**(*4S,5R*)-6-((*1R,2R*)-3-azido-1,2-dihydroxypropyl)-2,4-dihydroxy-5-(2-(methylselanyl)acetamido)tetrahydro-*2H*-pyran-2-carboxylate (17): 16** (400 mg, 0.70 mmol, 1.0 eq.) was dissolved in acetone (15 mL) and $H_2O$ (5 mL), and then $NaN_3$ (227.5 mg, 3.5 eq.) was added. The reaction was stirred overnight at reflux at 60°C. After removal of the solvent in vacuum, the residue was purified by silica flash chromatography with DCM: MeOH gradually from 4:1 to 1:1. Further purification via normal phase HPLC (XBridge Prep amide column,19 mm×250 mm, 5 µm) gave **17** as a light yellow solid (215 mg, 0.50 mmol, 72%), eluted with 0-95% water in ACN at the flow rate of 7 mL/min. **TLC** (Rf=0.4, DCM:MeOH:$H_2O$ =3:2:0.5, stained by 10% $H_2SO_4$ in MeOH). **$^1$H NMR:** (400 MHz, $D_2O$) δ 4.09 – 3.98 (m, 2H), 3.97 – 3.85 (m, 2H), 3.71 – 3.59 (m, 2H), 3.49 (dd, $J$ = 13.2, 5.6 Hz, 1H), 3.30 – 3.21 (m, 2H), 2.28 – 2.20 (m, 1H), 2.19 – 2.10 (m, 3H), 1.84 (dd, $J$ = 12.8, 11.6 Hz, 1H).**$^{13}$C NMR:** (101 MHz, $D_2O$) δ 176.67, 174.81, 96.39, 70.09, 69.04, 67.09, 53.89, 52.48, 39.56, 25.84, 4.98. **ESI-HRMS**: Calcd for $C_{12}H_{19}N_4O_8Se^-$ [M-H]$^-$ = 427.0373, found 427.0368

#### NMR Spectra

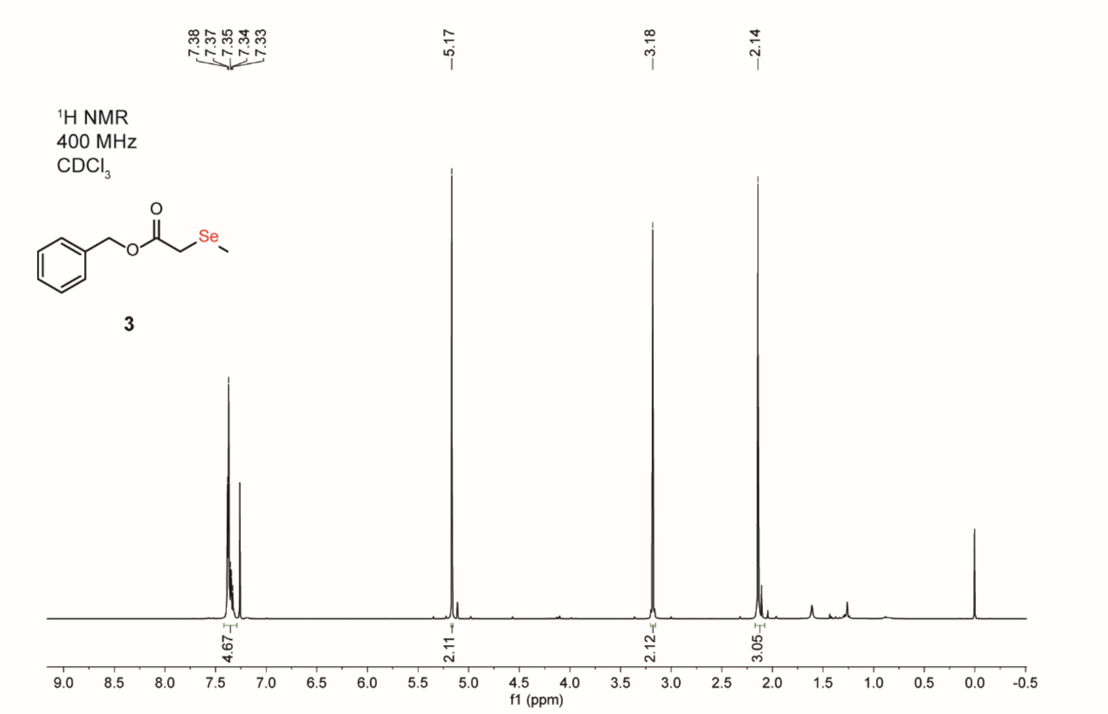

7.38
7.37
7.35
7.34
7.33
5.17
3.18
2.14
1H NMR
400 MHz
CDCl3
O
O
Se
3
4.67
2.11
2.12
3.05
9.0 8.5 8.0 7.5 7.0 6.5 6.0 5.5 5.0 4.5 4.0 3.5 3.0 2.5 2.0 1.5 1.0 0.5 0.0 -0.5
f1 (ppm)


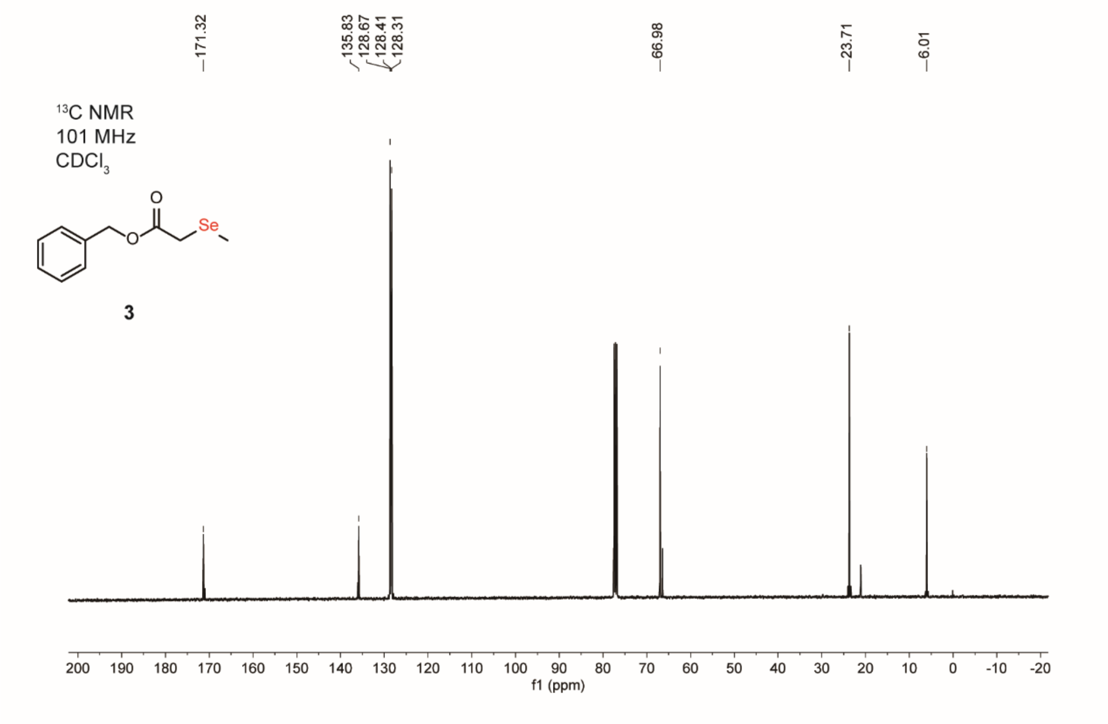

171.32
135.83
128.67
128.41
128.31
66.98
23.71
6.01
13C NMR
101 MHz
CDCl3
O
O
Se
3
200 190 180 170 160 150 140 130 120 110 100 90 80 70 60 50 40 30 20 10 0 -10 -20
f1 (ppm)

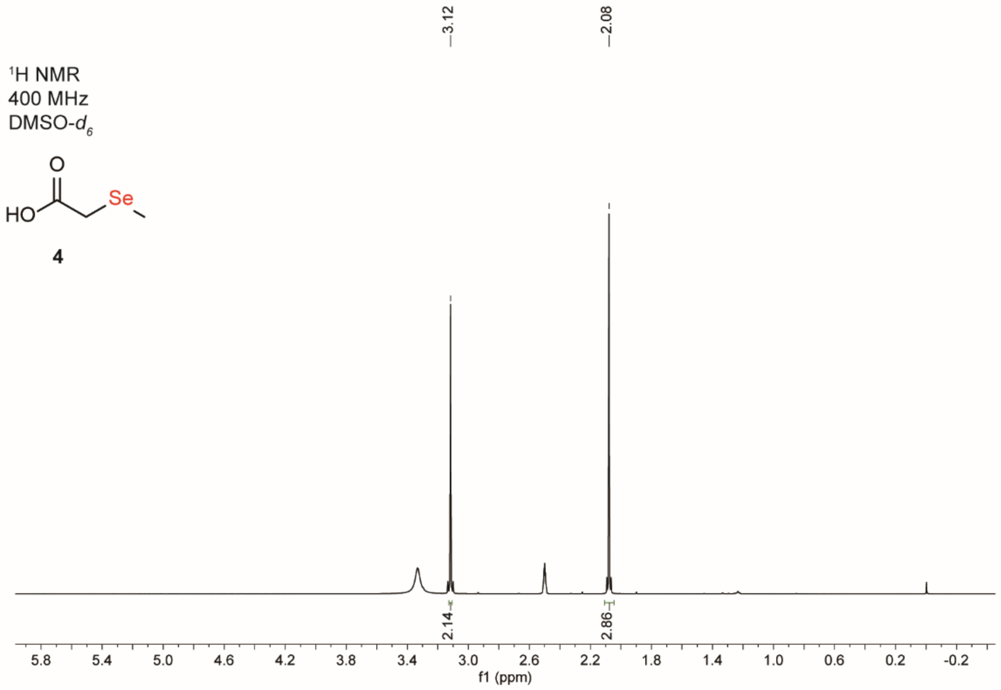

3.12
2.08
1H NMR
400 MHz
DMSO-d6
HO
O
Se
4
2.14
2.86
5.8
5.4
5.0
4.6
4.2
3.8
3.4
3.0
2.6
2.2
1.8
1.4
1.0
0.6
0.2
-0.2
f1 (ppm)


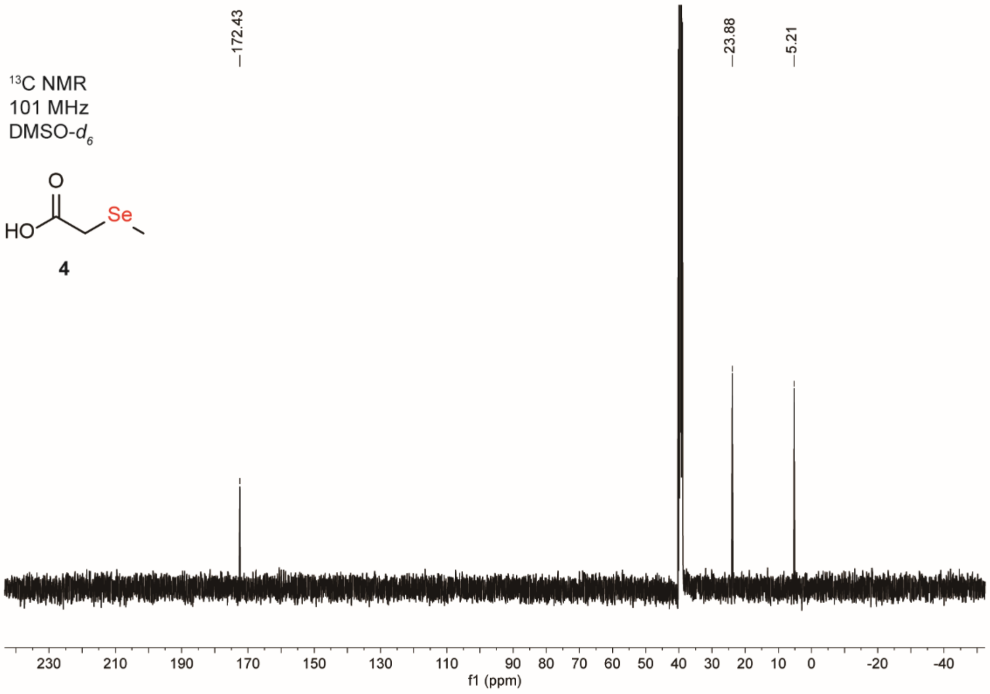

172.43
23.88
5.21
13C NMR
101 MHz
DMSO-d6
HO
O
Se
4
230
210
190
170
150
130
110
90
80
70
60
50
40
30
20
10
0
-20
-40
f1 (ppm)

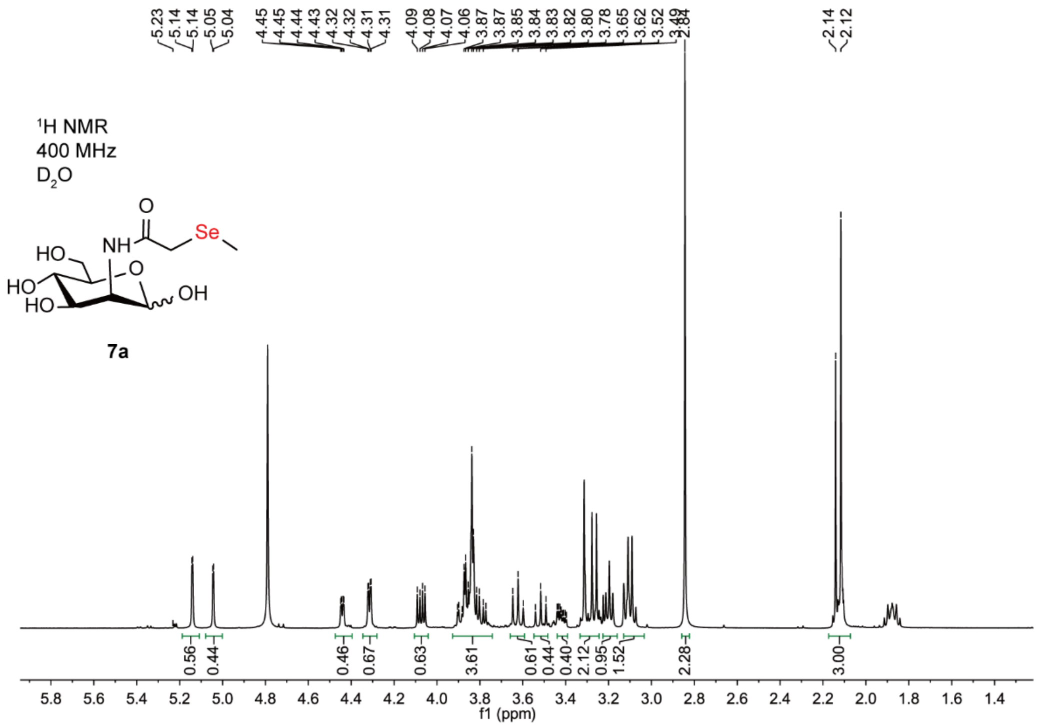
1H NMR
400 MHz
D2O
7a
f1 (ppm)

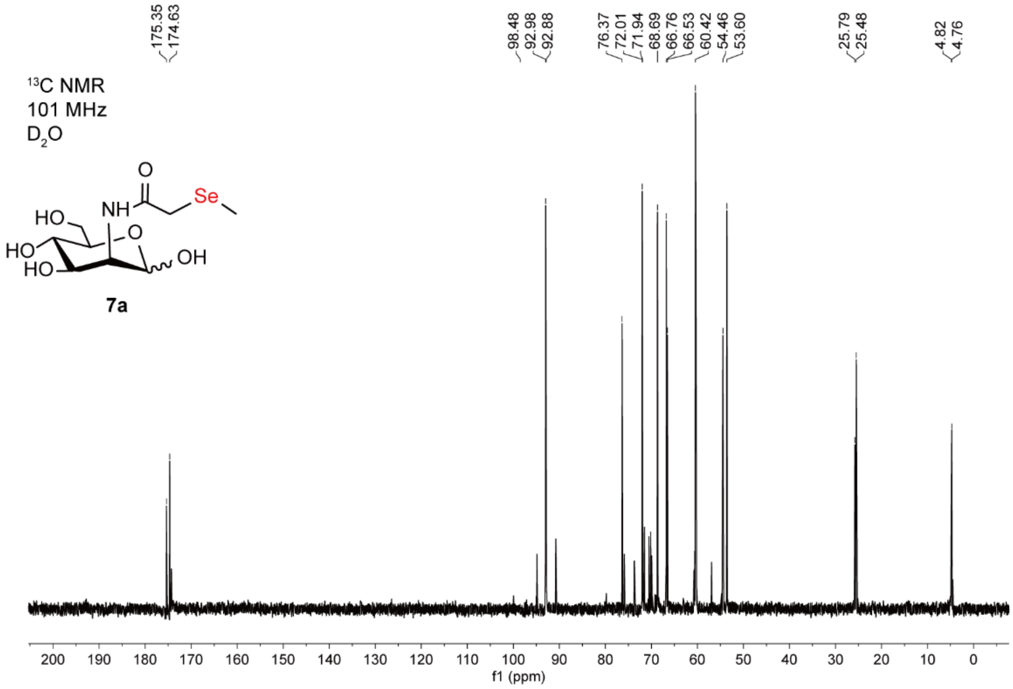
13C NMR
101 MHz
D2O
7a
f1 (ppm)

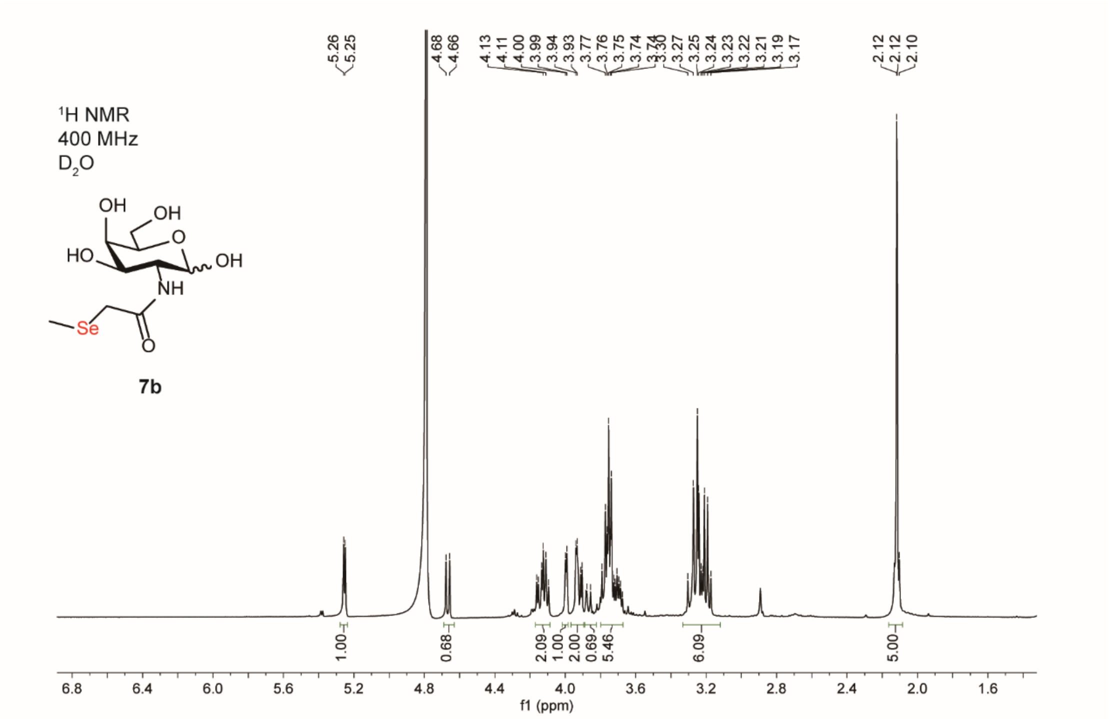

1H NMR
400 MHz
D2O
OH
OH
O
HO
OH
NH
Se
O
7b
5.26
5.25
4.68
4.66
4.13
4.11
4.00
3.99
3.94
3.93
3.77
3.76
3.75
3.74
3.30
3.27
3.25
3.24
3.23
3.22
3.21
3.19
3.17
2.12
2.12
2.10
1.00
0.68
2.09
1.00
2.00
0.69
5.46
6.09
5.00
f1 (ppm)


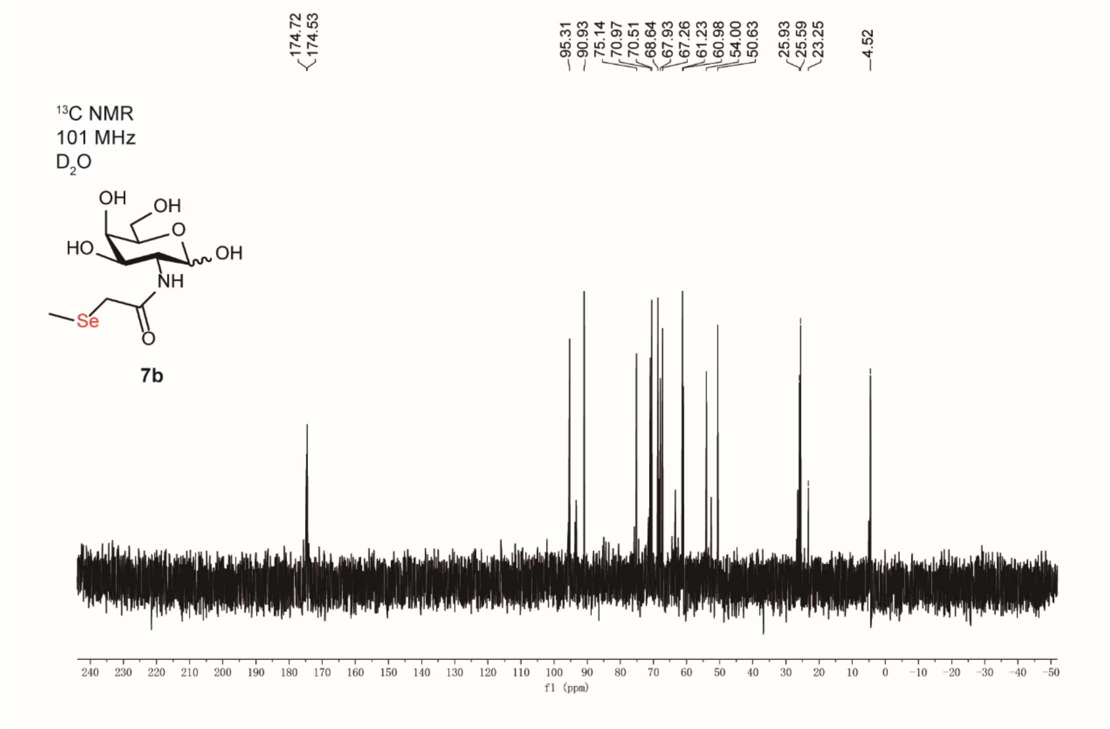

13C NMR
101 MHz
D2O
OH
OH
O
HO
OH
NH
Se
O
7b
174.72
174.53
95.31
90.93
75.14
70.97
70.51
68.64
67.93
67.26
61.23
60.98
54.00
50.63
25.93
25.59
23.25
4.52
f1 (ppm)

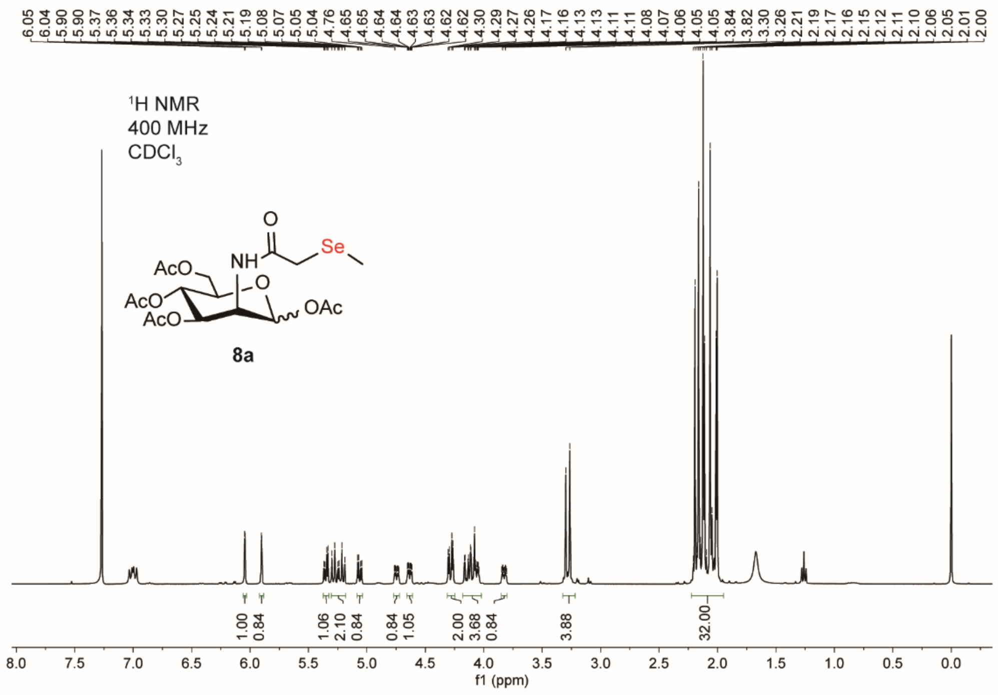

1H NMR
400 MHz
CDCl3
8a
f1 (ppm)


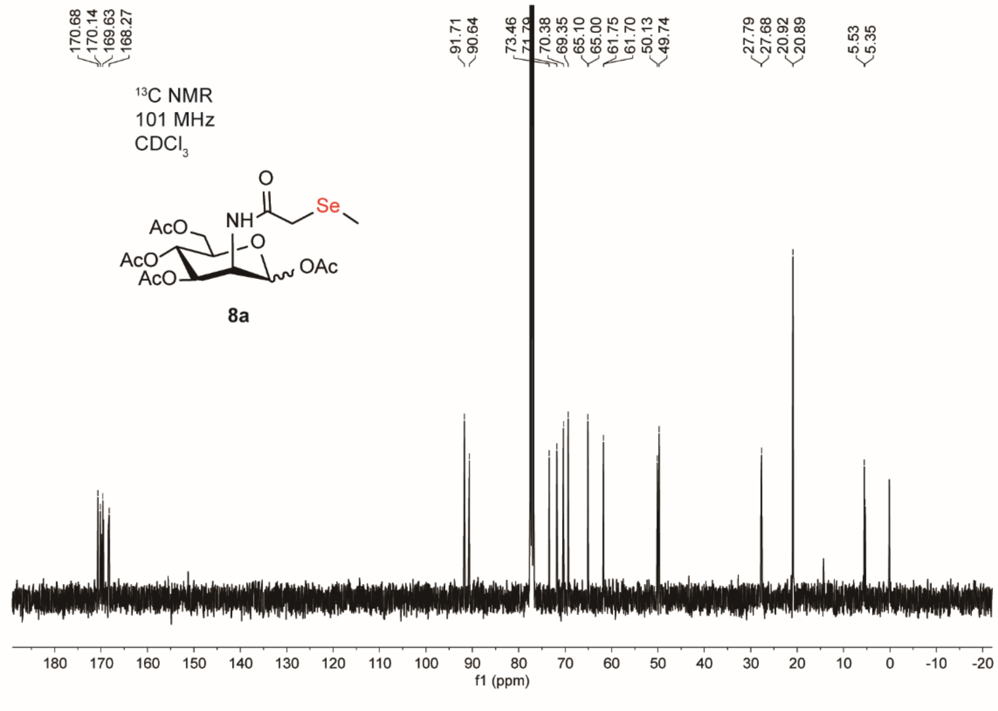

13C NMR
101 MHz
CDCl3
8a
f1 (ppm)

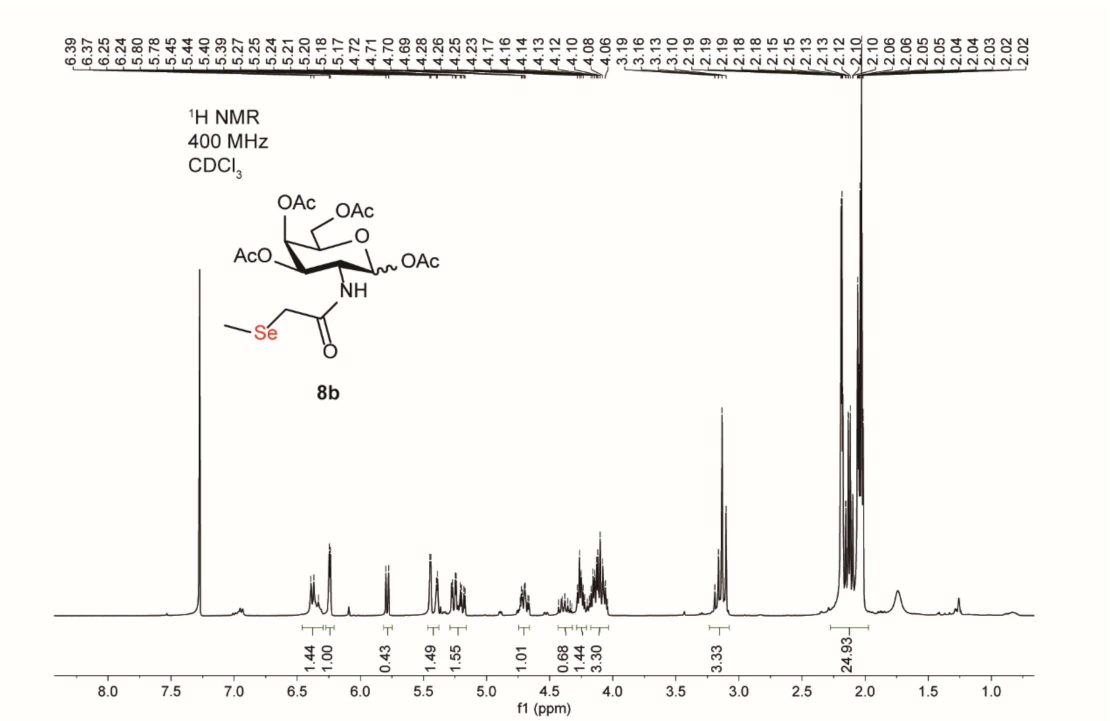
6.39 6.37 6.25 6.24 5.80 5.78 5.45 5.44 5.40 5.39 5.27 5.25 5.24 5.21 5.20 5.18 5.17 4.72 4.71 4.70 4.69 4.28 4.26 4.25 4.23 4.17 4.16 4.14 4.13 4.12 4.10 4.08 4.06 3.19 3.16 3.13 3.10 2.19 2.19 2.19 2.18 2.15 2.15 2.13 2.13 2.12 2.10 2.10 2.06 2.06 2.05 2.05 2.04 2.04 2.03 2.02 2.02
1H NMR
400 MHz
CDCl3
OAc
OAc
O
AcO
OAc
NH
Se
O
8b
1.44 1.00 0.43 1.49 1.55 1.01 0.68 1.44 3.30 3.33 24.93
8.0 7.5 7.0 6.5 6.0 5.5 5.0 4.5 4.0 3.5 3.0 2.5 2.0 1.5 1.0
f1 (ppm)

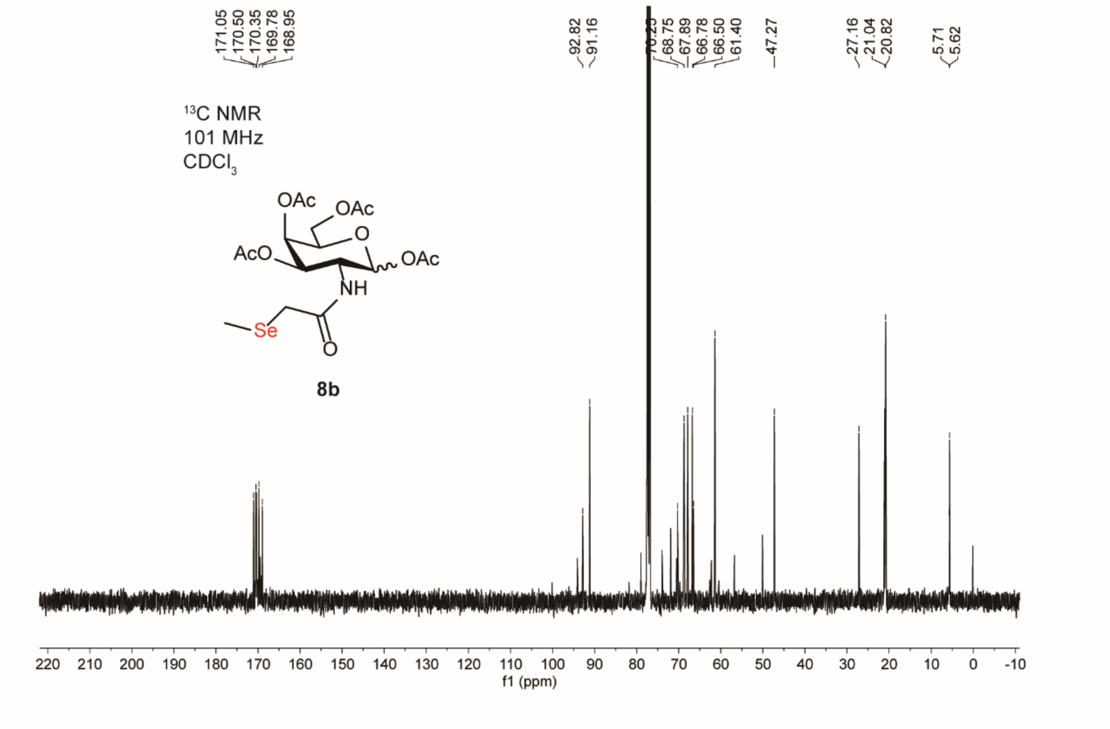
171.05 170.50 170.35 169.78 168.95
92.82 91.16
68.75 67.89 66.78 66.50 61.40
47.27
27.16 21.04 20.82
5.71 5.62
13C NMR
101 MHz
CDCl3
OAc
OAc
O
AcO
OAc
NH
Se
O
8b
220 210 200 190 180 170 160 150 140 130 120 110 100 90 80 70 60 50 40 30 20 10 0 -10
f1 (ppm)

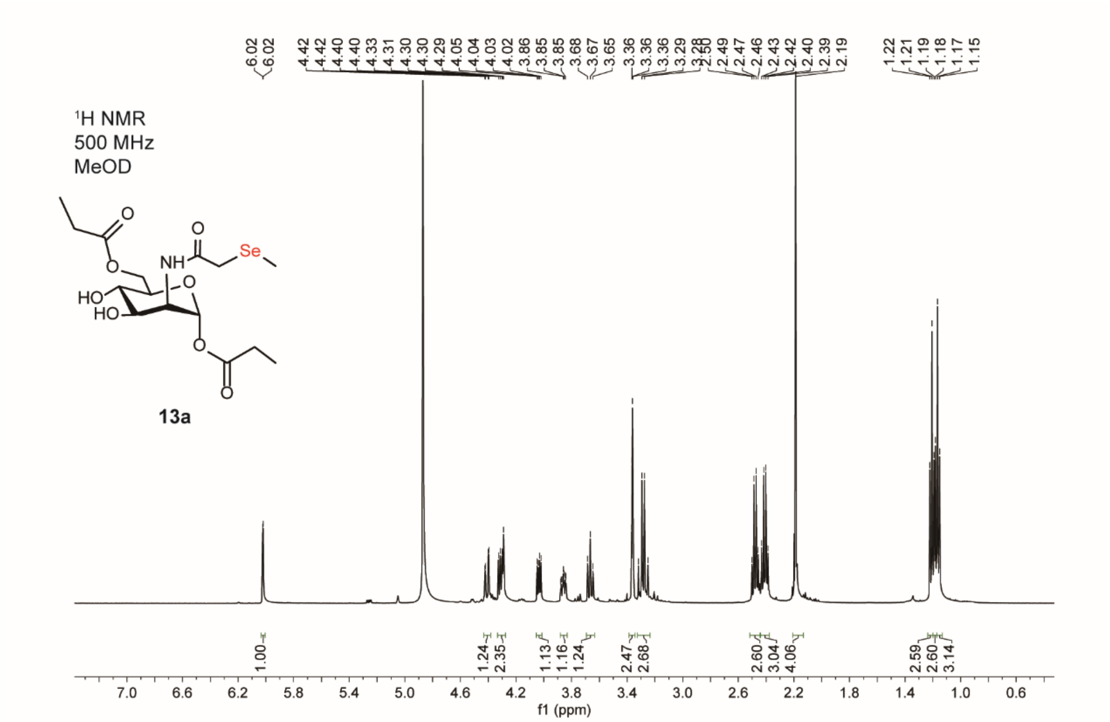
6.02
6.02
4.42
4.42
4.40
4.40
4.33
4.31
4.30
4.30
4.29
4.05
4.04
4.03
4.02
3.86
3.85
3.85
3.68
3.67
3.65
3.36
3.36
3.36
3.29
2.49
2.47
2.46
2.43
2.42
2.40
2.39
2.19
1.22
1.21
1.19
1.18
1.17
1.15
¹H NMR
500 MHz
MeOD
O
NH
O
Se
O
HO
HO
O
O
O
13a
1.00
1.24
2.35
1.13
1.16
1.24
2.47
2.68
2.60
3.04
4.06
2.59
2.60
3.14
7.0 6.6 6.2 5.8 5.4 5.0 4.6 4.2 3.8 3.4 3.0 2.6 2.2 1.8 1.4 1.0 0.6
f1 (ppm)

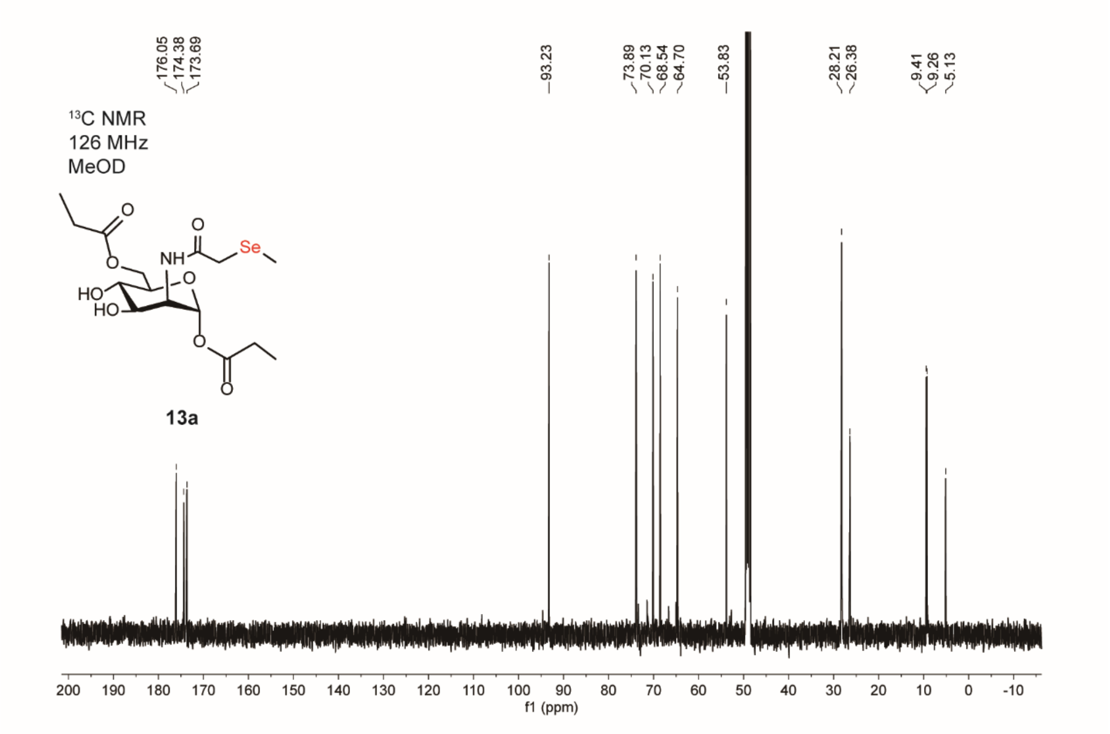
176.05
174.38
173.69
93.23
73.89
70.13
68.54
64.70
53.83
28.21
26.38
9.41
9.26
5.13
¹³C NMR
126 MHz
MeOD
O
NH
O
Se
O
HO
HO
O
O
O
13a
200 190 180 170 160 150 140 130 120 110 100 90 80 70 60 50 40 30 20 10 0 -10
f1 (ppm)

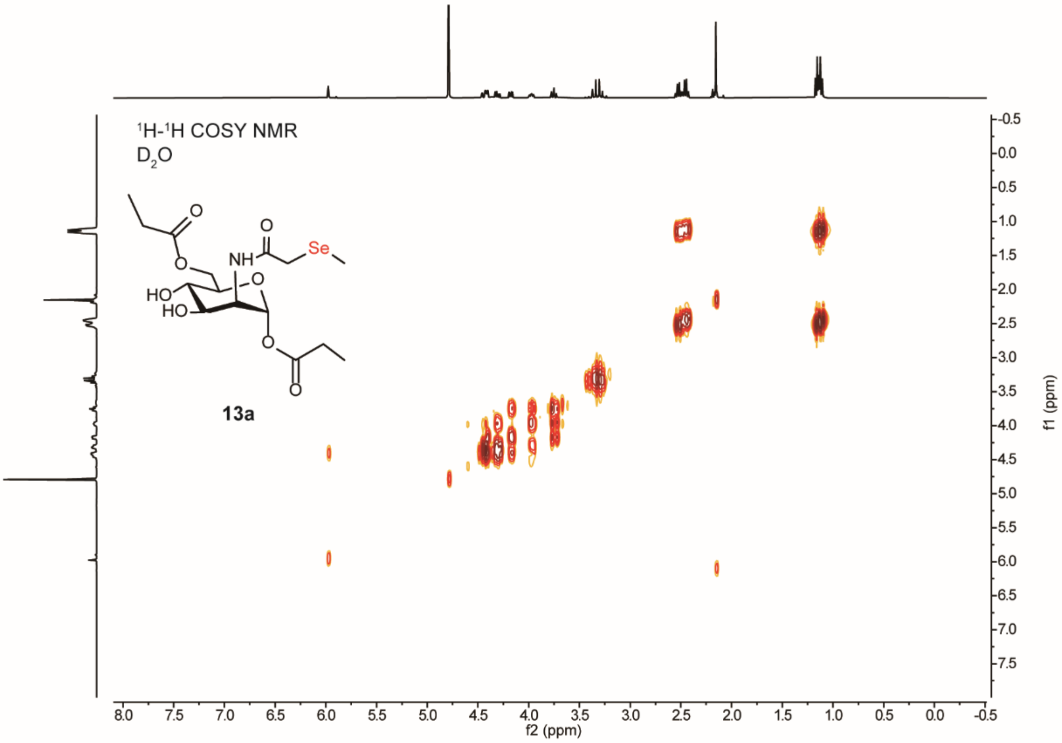

¹H-¹H COSY NMR
D₂O
NH
O
Se
HO
HO
13a
f2 (ppm)
f1 (ppm)


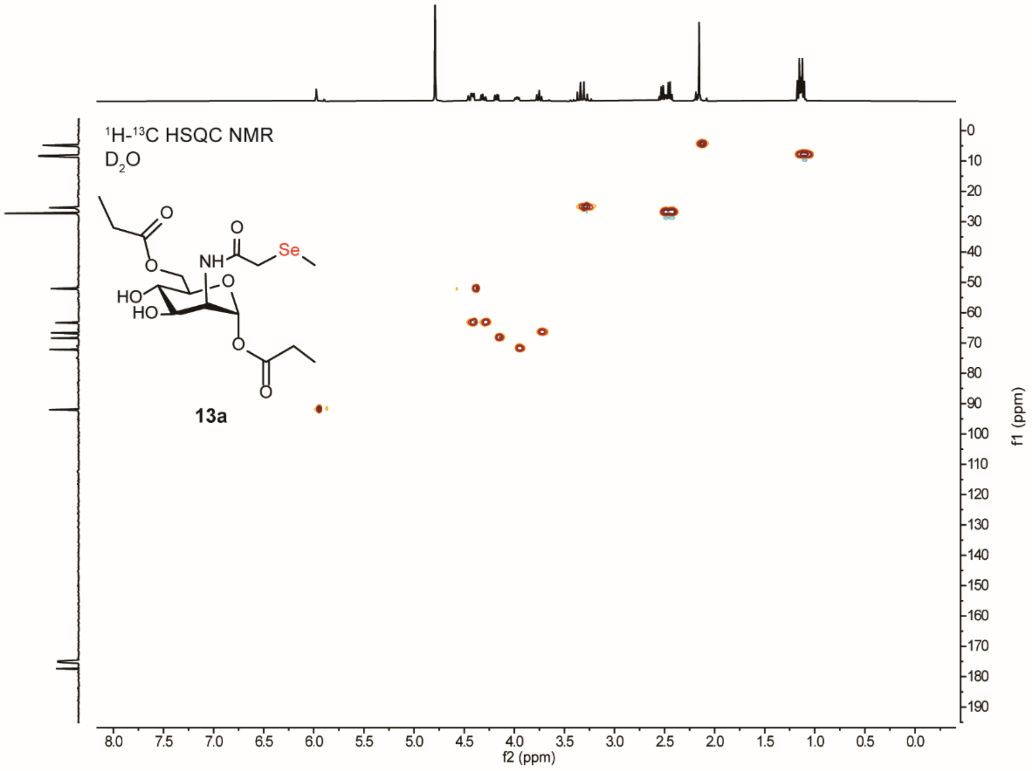

¹H-¹³C HSQC NMR
D₂O
NH
O
Se
HO
HO
13a
f2 (ppm)
f1 (ppm)

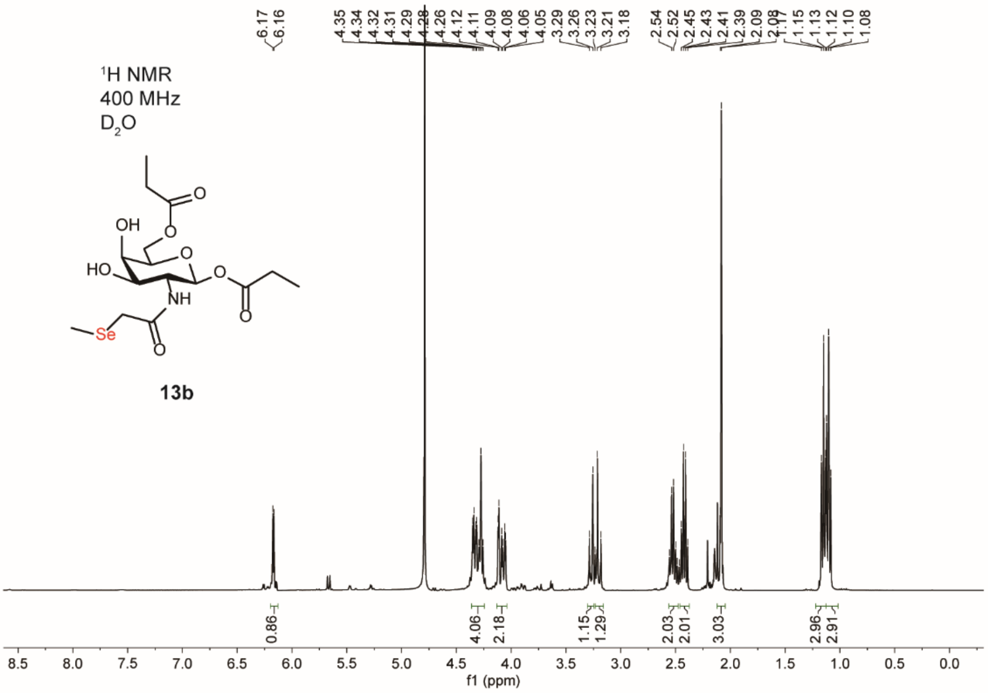
1H NMR
400 MHz
D2O
13b
6.17
6.16
4.35
4.34
4.32
4.31
4.29
4.28
4.26
4.12
4.11
4.09
4.08
4.06
4.05
3.29
3.26
3.23
3.21
3.18
2.54
2.52
2.45
2.43
2.41
2.39
2.09
2.08
1.17
1.15
1.13
1.12
1.10
1.08
0.86
4.06
2.18
1.15
1.29
2.03
2.01
3.03
2.96
2.91
f1 (ppm)

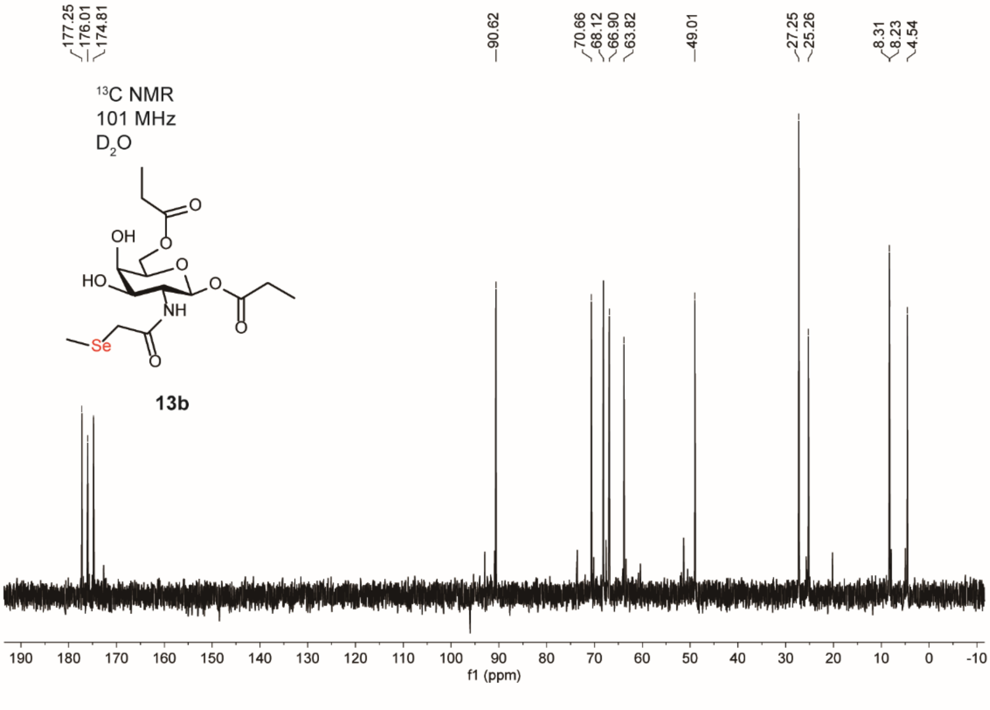
13C NMR
101 MHz
D2O
13b
177.25
176.01
174.81
90.62
70.66
68.12
66.90
63.82
49.01
27.25
25.26
8.31
8.23
4.54
f1 (ppm)

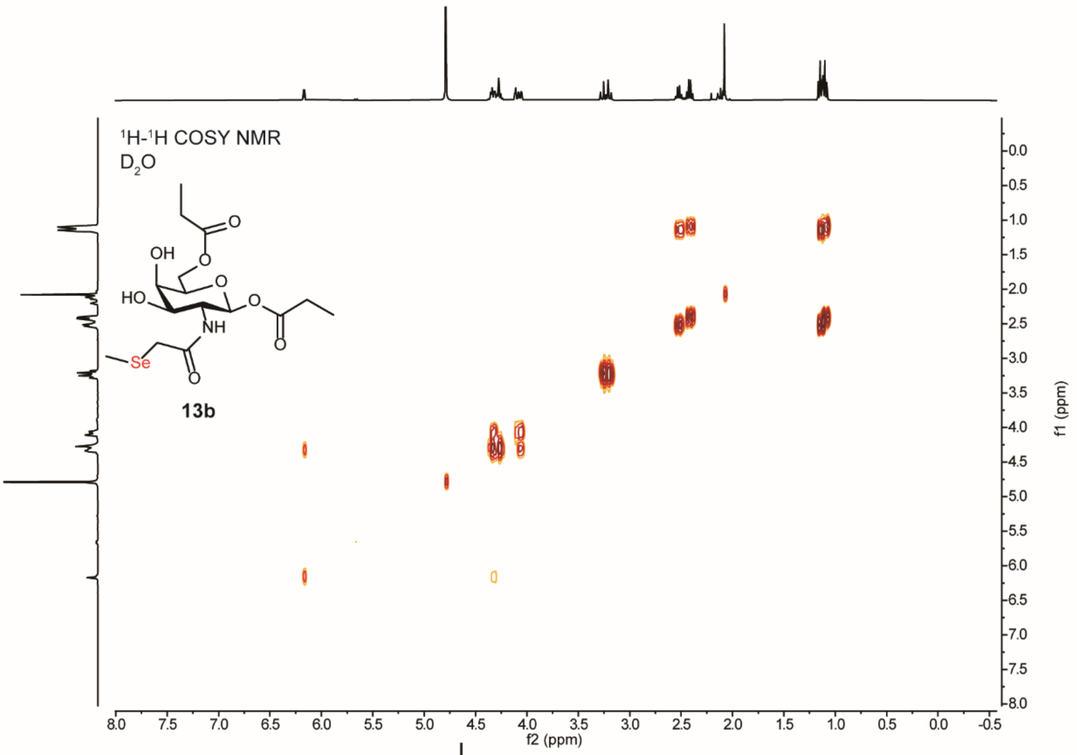
1H-1H COSY NMR
D2O
OH
HO
O
O
O
NH
Se
O
O
13b
f2 (ppm)
f1 (ppm)

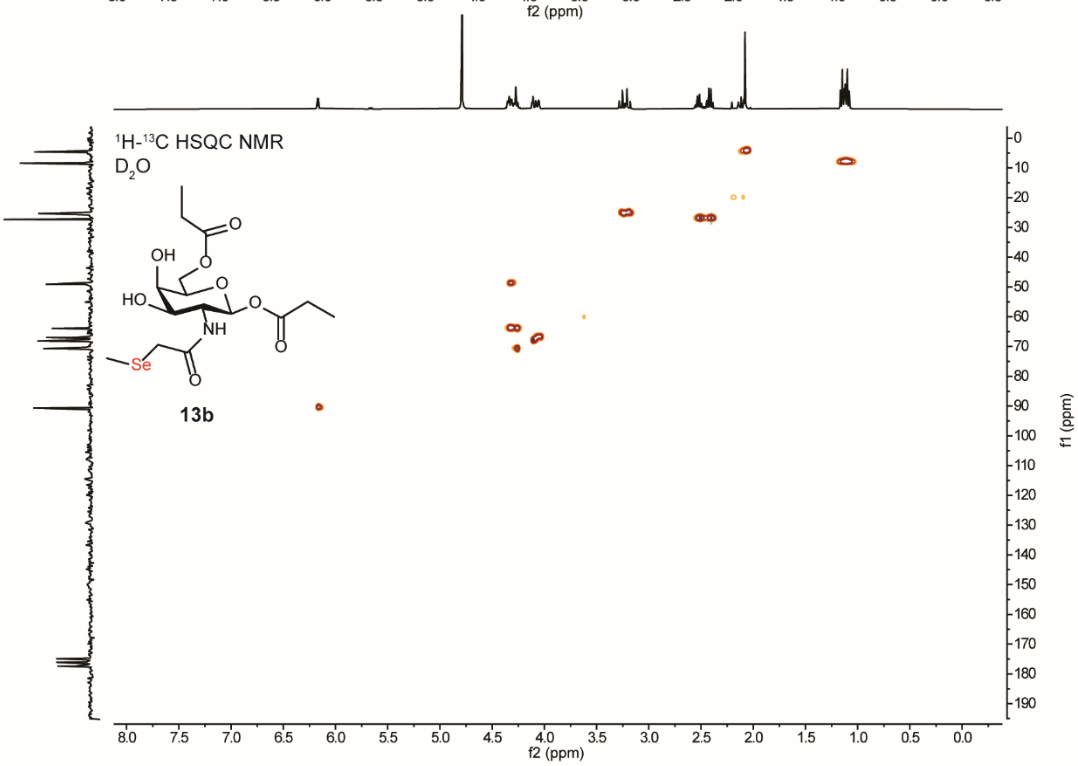
1H-13C HSQC NMR
D2O
OH
HO
O
O
O
NH
Se
O
O
13b
f2 (ppm)
f1 (ppm)

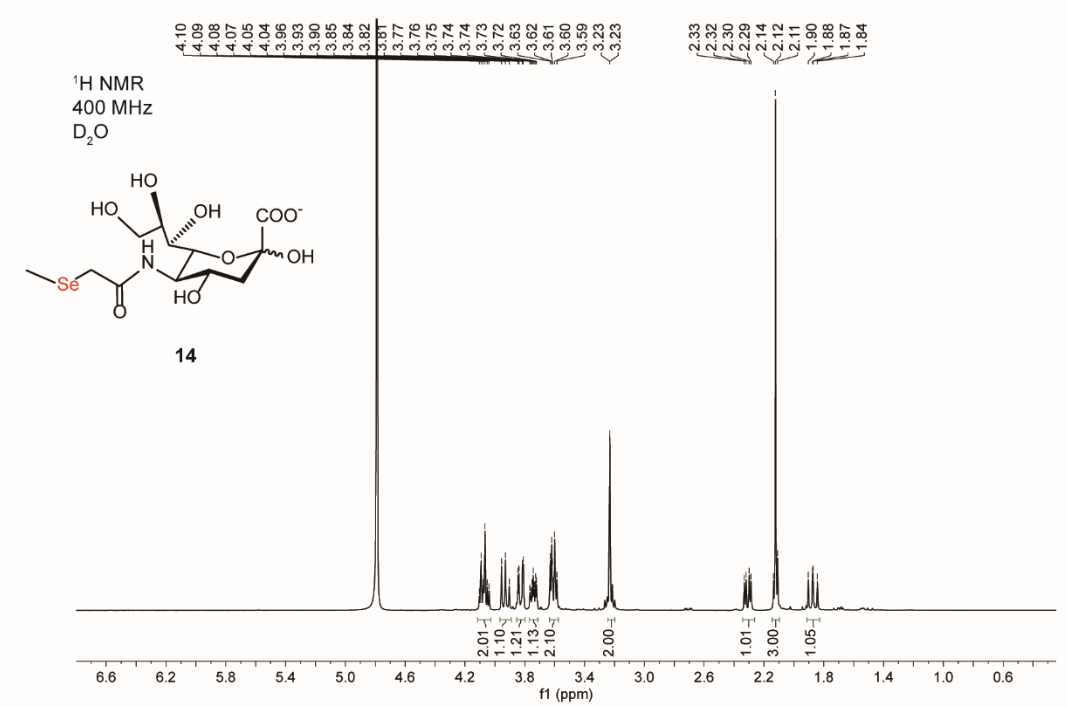
1H NMR
400 MHz
D2O
HO
HO
OH
COO-
H
N
O
OH
Se
HO
O
14
4.10 4.09 4.08 4.07 4.05 4.04 3.96 3.93 3.90 3.85 3.84 3.82 3.81 3.77 3.76 3.75 3.74 3.74 3.73 3.72 3.63 3.62 3.61 3.60 3.59 3.23 3.23
2.33 2.32 2.30 2.29 2.14 2.12 2.11 1.90 1.88 1.87 1.84
2.01 1.10 1.21 1.13 2.10 2.00 1.01 3.00 1.05
f1 (ppm)

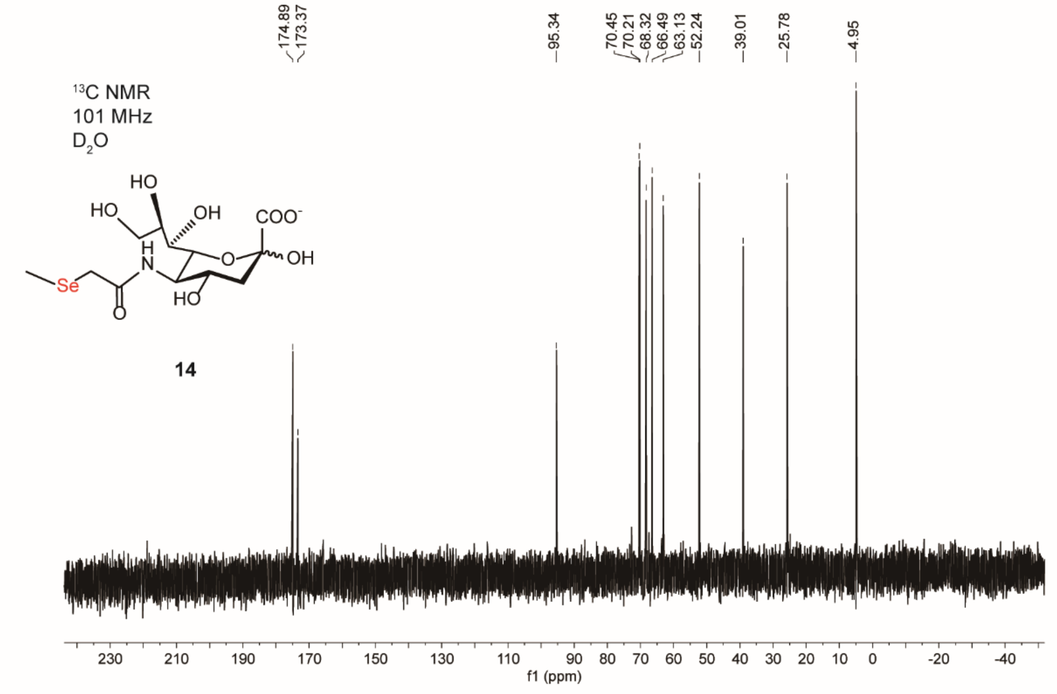
13C NMR
101 MHz
D2O
HO
HO
OH
COO-
H
N
O
OH
Se
HO
O
14
174.89 173.37
95.34
70.45 70.21 68.32 66.49 63.13 52.24
39.01
25.78
4.95
f1 (ppm)

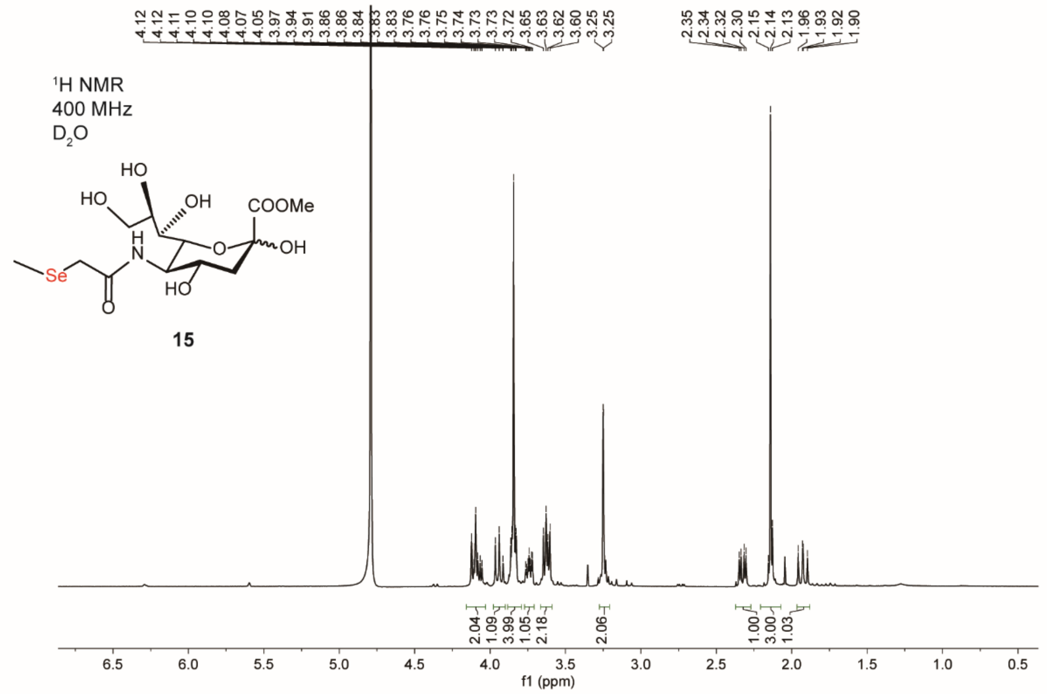
4.12 4.12 4.11 4.10 4.10 4.08 4.07 4.05 3.97 3.94 3.91 3.86 3.86 3.84 3.83 3.83 3.76 3.75 3.74 3.73 3.73 3.72 3.65 3.63 3.62 3.60 3.25 3.25
2.35 2.34 2.32 2.30 2.15 2.14 2.13 1.96 1.93 1.92 1.90
1H NMR
400 MHz
D2O
HO
HO
OH
COOMe
H
N
O
OH
Se
O
HO
15
2.04 1.09 3.99 1.05 2.18 2.06 1.00 3.00 1.03
6.5 6.0 5.5 5.0 4.5 4.0 3.5 3.0 2.5 2.0 1.5 1.0 0.5
f1 (ppm)

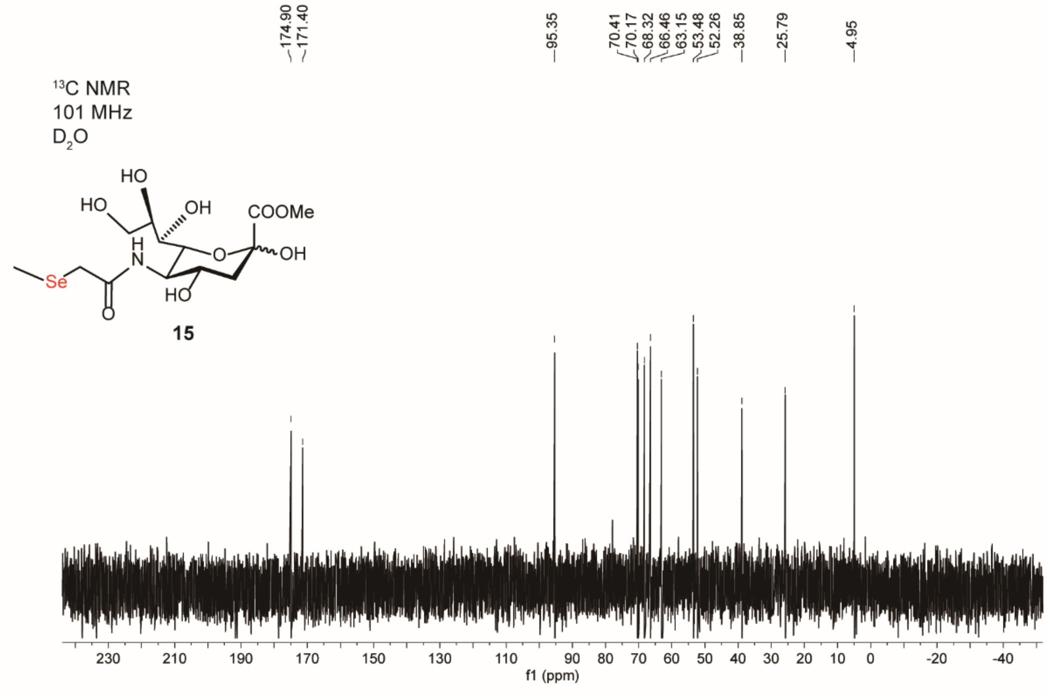
174.90 171.40 95.35 70.41 70.17 68.32 66.46 63.15 53.48 52.26 38.85 25.79 4.95
13C NMR
101 MHz
D2O
HO
HO
OH
COOMe
H
N
O
OH
Se
O
HO
15
230 210 190 170 150 130 110 90 80 70 60 50 40 30 20 10 0 -20 -40
f1 (ppm)

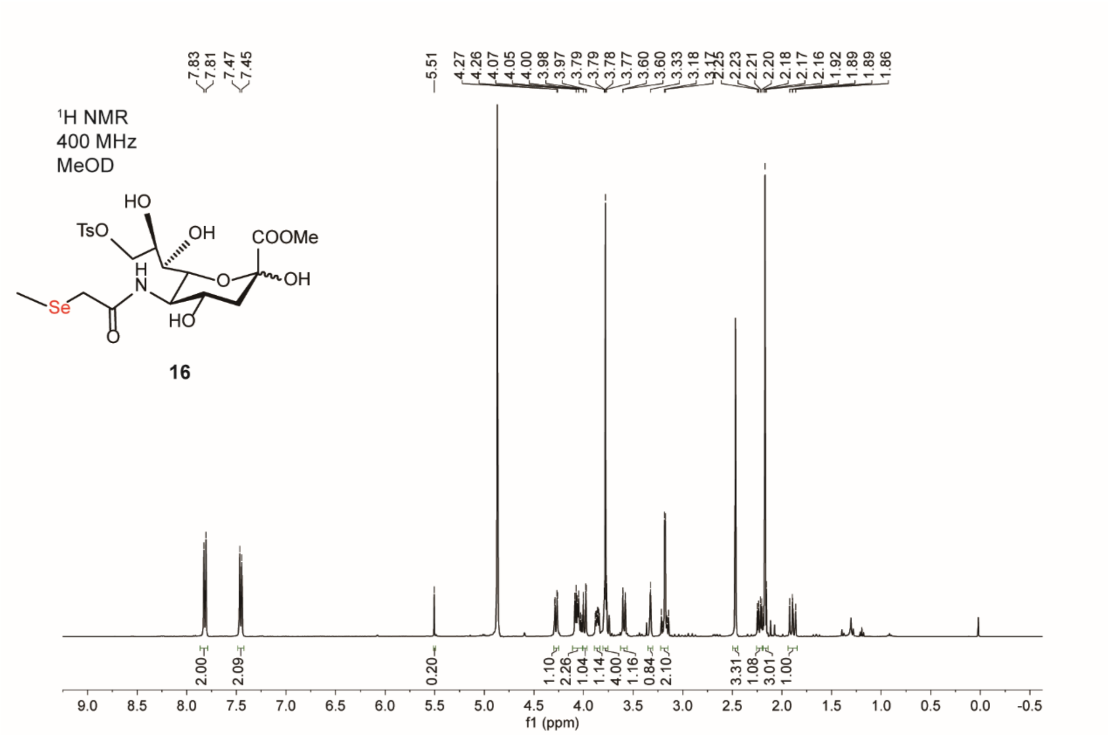

¹H NMR
400 MHz
MeOD
16
f1 (ppm)


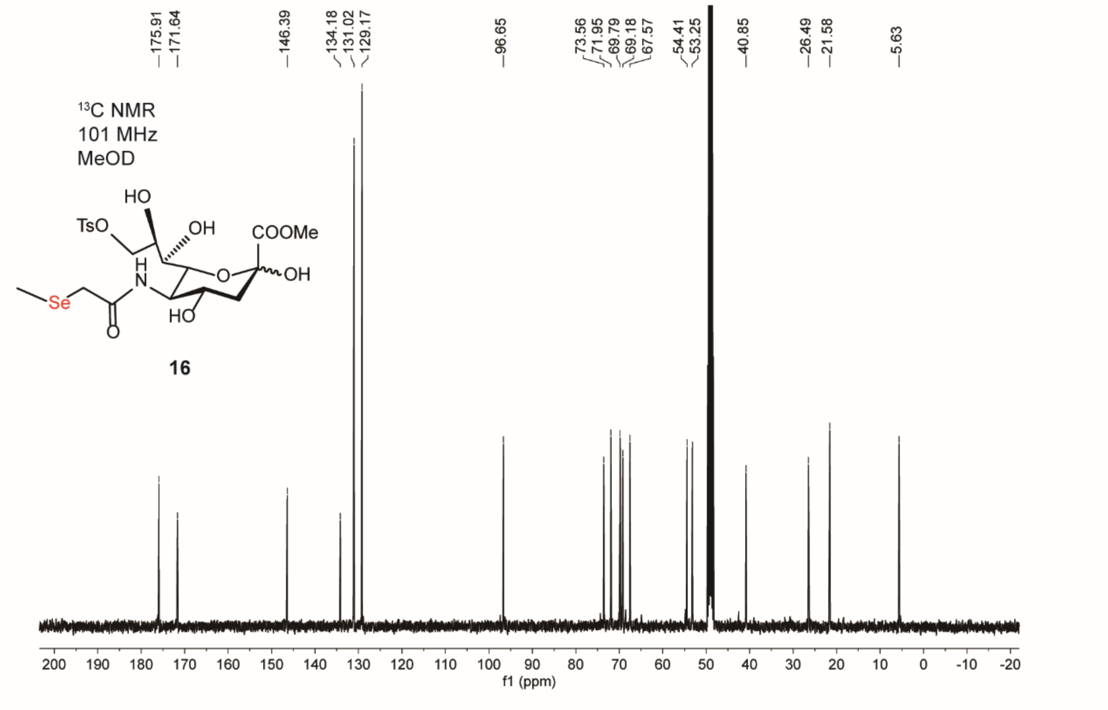

¹³C NMR
101 MHz
MeOD
16
f1 (ppm)

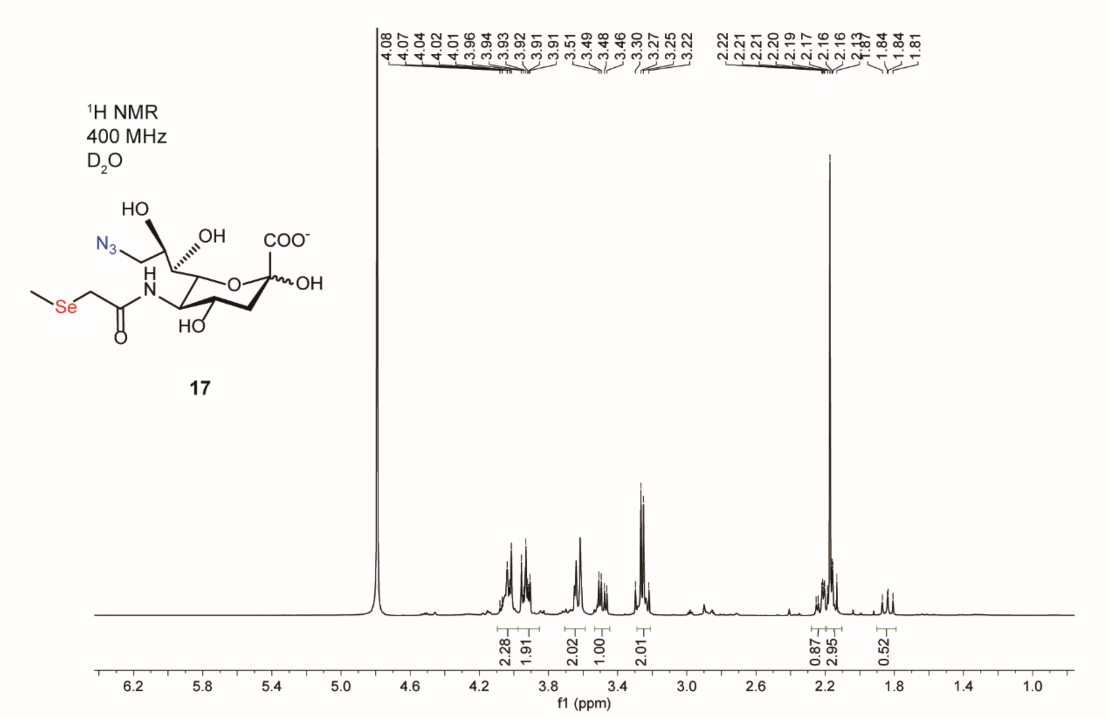

1H NMR
400 MHz
$D_2O$
HO
OH
$N_3$
COO-
H
N
O
OH
Se
HO
O
17
4.08 4.07 4.04 4.02 4.01 3.96 3.94 3.93 3.92 3.91 3.91 3.51 3.49 3.48 3.46 3.30 3.27 3.25 3.22 2.22 2.21 2.21 2.20 2.19 2.17 2.16 2.16 2.13 1.87 1.84 1.84 1.81
2.28 1.91 2.02 1.00 2.01 0.87 2.95 0.52
6.2 5.8 5.4 5.0 4.6 4.2 3.8 3.4 3.0 2.6 2.2 1.8 1.4 1.0
f1 (ppm)


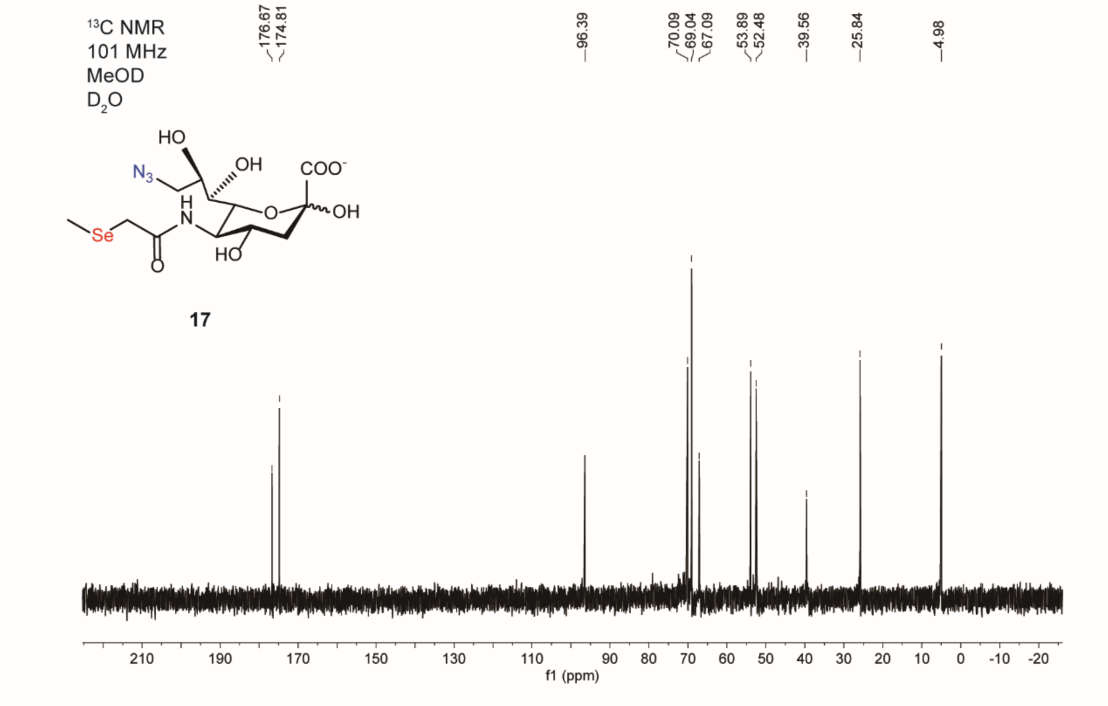

13C NMR
101 MHz
MeOD
$D_2O$
HO
OH
$N_3$
COO-
H
N
O
OH
Se
HO
O
17
176.67 174.81 96.39 70.09 69.04 67.09 53.89 52.48 39.56 25.84 4.98
210 190 170 150 130 110 90 80 70 60 50 40 30 20 10 0 -10 -20
f1 (ppm)

### Supplementary Note 2: Flow cytometry gating strategy

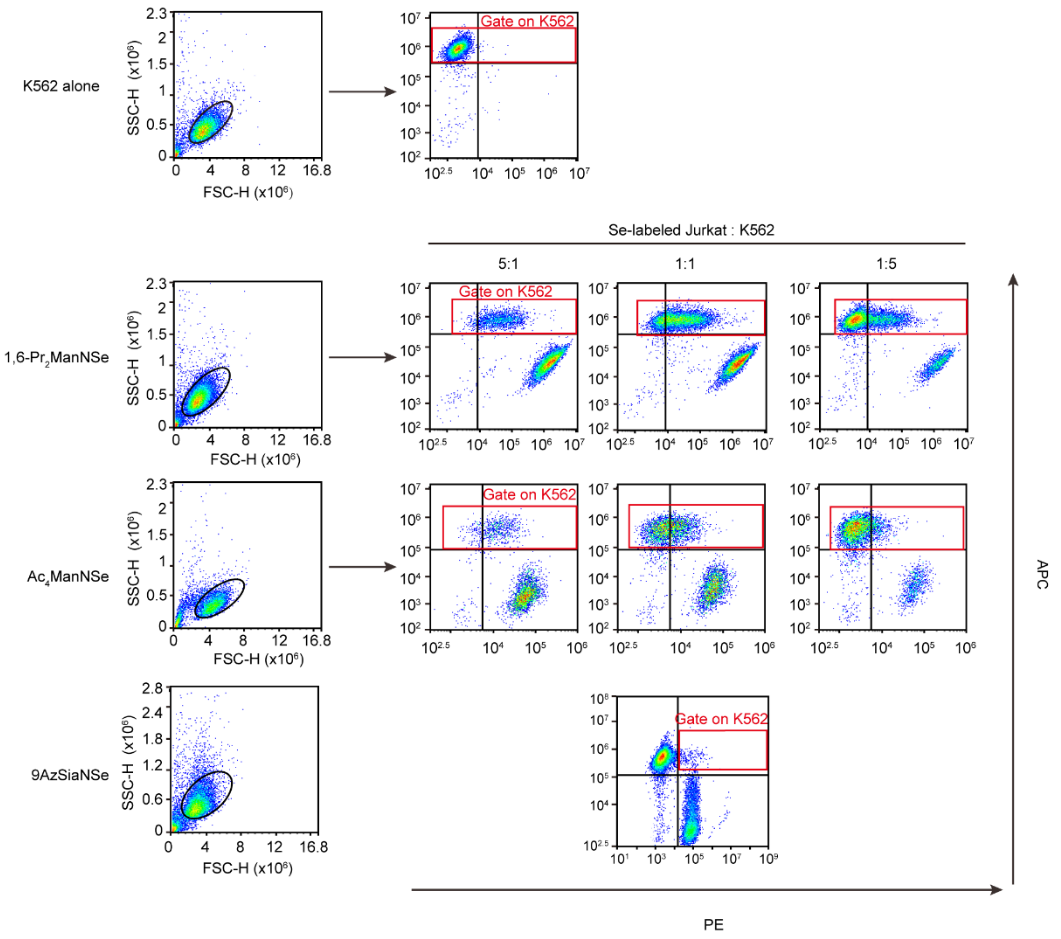

**Supplementary Table 1: The instrumental and operational parameters of ICP-MS solution analysis**

| ICP-MS | Parameter |
|---|---|
| Forward power (W) | 1600 |
| Nebulizer gas (L $min^{-1}$) | 1 |
| Cool gas (L $min^{-1}$) | 18 |
| Auxiliary gas (L $min^{-1}$) | 1.2 |
| Spray chamber | Cyclone |
| Interface | Ni Cone |
| Nebulizer | Concentric |
| Isotope | $^{78}Se$ |
| Uptake Rate (mL $min^{-1}$) | 0.3 |

**Supplementary Table 2: Instrumental parameters of LA-ICP-MS for selenoprotein analysis on PVDF membrane**

| | Parameter |
|---|---|
| **ICP-MS** | |
| RF power | 1600W |
| Nebulizer gas | 1.2 L/min |
| Auxiliary gas | 1.2 L/min |
| Plasma gas | 18.0 L/min |
| Measurement mode | Standard mode |
| Cone materials | Ni |
| Dwell time | 10 ms |
| Isotope | $^{78}Se$ |
| **Laser ablation system** | |
| Laser wavelength | 193 nm |
| Sample introduction system | ARIS |
| Ablation mode | Linear ablation |
| Spot size | 100 µm |
| Laser fluency | 0.90 J/cm$^2$ |
| Laser repetition rate | 100 Hz |
| Line scan velocity | 1 mm/s |
| He flow of inner cell | 0.3 L/min |
| He flow of outer cell | 0.3 L/min |

**Supplementary Table 3: Instrumental parameters of LA-ICP-TOF MS for mouse tissue sialoglycan imaging**

| | **Parameter** |
|---|---|
| **ICP-TOFMS** | |
| RF power | 1550W |
| Nebulizer gas | 0.89 L/min |
| Auxiliary gas | 0.8 L/min |
| Plasma gas | 14.0 L/min |
| Measurement mode | CCT mode |
| CCT flow | 4.5 mL/min |
| m/z range | 14-256 |
| Cone materials | Ni |
| Notch mass | 28, 32, 40, 80 |
| Injector diameter | 2 mm |
| TOF extraction time | 46 µs |
| Waveform | 516 |
| **Laser ablation system** | |
| Laser wavelength | 193 nm |
| Sample introduction system | ARIS |
| Ablation mode | Spot ablation |
| Spot size | 20 µm |
| Laser fluency | 0.90 J /$cm^2$ |
| Stage movement | 5000 µm/s |
| He flow of inner cell | 0.45 L/min |
| He flow of outer cell | 0.15 L/min |
| Laser repetition rate | 250 Hz |

**Supplementary Table 4: Instrumental parameters of LA-ICP-MS for mouse tissue sialoglycan quantification *in situ***

| | Parameter |
|---|---|
| **ICP-MS** | |
| RF power | 1600W |
| Nebulizer gas | 1.2 L/min |
| Auxiliary gas | 1.2 L/min |
| Plasma gas | 18.0 L/min |
| Measurement mode | Standard mode |
| Cone materials | Ni |
| Dwell time | 10 ms |
| Isotope | $^{78}Se$ |
| **Laser ablation system** | |
| Laser wavelength | 193 nm |
| Sample introduction system | ARIS |
| Ablation mode | Spot ablation |
| Spot size | 20 μm |
| Offset of Scans | 40 μm |
| Laser fluency | 0.90 J /cm$^2$ |
| Stage movement | 2000 μm/s |
| He flow of inner cell | 0.3 L/min |
| He flow of outer cell | 0.3 L/min |
| Laser repetition rate | 100 Hz |
